# Allosteric activation and inhibition of glycogen phosphorylase share common transient structural features

**DOI:** 10.1101/2023.06.03.543583

**Authors:** Monika Kish, Dylan P. Ivory, Jonathan J. Phillips

## Abstract

It remains a major challenge to ascertain the specific structurally dynamic changes that underpin protein functional switching. There is a growing need to complement structural models with the ability to determine the dynamic structural changes that occur as these proteins are regulated and function. The archetypal allosteric enzyme, glycogen phosphorylase is one of the most studied proteins and is a clinical target of much interest to treat type II diabetes and metastatic cancers. However, a lack of understanding of its complex regulation, mediated by dynamic structural changes, hinder its exploitation as a drug target. Here, we precisely locate dynamic structural changes upon allosteric activation and inhibition of glycogen phosphorylase, by developing a time-resolved non-equilibrium millisecond hydrogen/deuterium-exchange mass spectrometry (HDX-MS) approach. We resolved obligate transient changes in localized structure that are absent when directly comparing active/inactive states of the enzyme and show that they are common to allosteric activation by AMP and inhibition by caffeine, operating at different sites. This indicates that opposing allosteric regulation by inhibitor and activator ligands is mediated by pathways that intersect at a common structurally dynamic motif. This approach has broad application to determine the structural kinetic mechanisms by which proteins are regulated.

## INTRODUCTION

Allostery refers to the transfer of a signal between two sites of a protein resulting in a change in the catalytic activity to a substrate or binding affinity to a ligand at a remote site of the protein. These sophisticated structural transitions are fundamental to receptor transduction^1^, cell signalling^2^, and metabolic regulation^3^. While allostery is typically associated with ligand binding to the allosteric site, connected to conformational change^4^, these structural transitions can also be triggered by covalent modification (e.g., phosphorylation)^5^, or proteolytic cleavage^6^. Despite extensive studies since its inception^7-9^, very little is known about the core dynamic process of allosteric communication, precisely how these signals are transmitted long distances through a protein molecule. This is in large part due to the lack of biophysical approaches to measure these signals at high resolution in both time and space. Recently, important advances have been made towards this goal in single molecule FRET^10,11,12,13^, NMR^14^, time resolved cryo-EM^15^, time resolved crystallography^16^, dynamic simulations^17^ and Double Electron Electron Resonance (DEER)^18,19^. Perhaps the most direct evidence for an intramolecular mechanism comes from infrared laser absorbance and emission experiments of conjugated peptides, which strongly argues that the vibrational energy transfer (VET) is mediated by hydrogen-bonds. Collectively, these studies begin to reveal non-equilibrium protein structural dynamics at high structural and temporal resolution, first described by Monod and colleagues in 1965^20^. Though impressive, they still require considerable adaptation per sample and are not yet broadly applicable to cases of ligand-induced allostery in proteins.

Glycogen phosphorylase (GlyP) is the archetypal allosteric enzyme whose regulation is tightly coupled to solid tumor metastasis^21^, type II diabetes^22^ and adaptive immunity (early memory CD8+ T-cell recall response)^23^. GlyP catalyzes the first committed energetic step of carbohydrate metabolism from glycogen stores and as such is one of the most highly regulated enzymes known, with at least six allosteric sites that sense natural ligands and drugs. GlyP has been intensively studied since the 1930s^24^, thus there is abundance of structural and kinetic data on this protein system^25-34^, with detailed studies on the activation mechanism^35^ and stabilization of R/T-states (active/inactive-states) by allosteric ligands^36^ and phosphorylation^37^. GlyP is, therefore, an ideal system to challenge our ability to discern dynamic structural mechanisms of allosteric regulation as it is alternately activated by AMP at the ‘nucleotide site’ and inhibited by caffeine at the ‘inhibitor site’^38,39^, whilst also being constitutively active by Ser 14 phosphorylation^20^. Binding of AMP at the allosteric effector site brings similar changes to phosphorylation but believed to be through somewhat different mechanisms^40^. How a single protein can support allosteric activation and inhibition by multiple intersecting structural pathways has not been elucidated.

Hydrogen-deuterium exchange mass spectrometry (HDX-MS) is a sensitive measure of solvent accessible hydrogen-bonding of polypeptide backbone amide groups. The amide H-bonds directly relate to local structural dynamics and now can be routinely determined in large protein systems^41^, with millisecond time resolution^42^ and at or near single amino acid structural resolution^43^. Here, we used a non-equilibrium millisecond HDX-MS approach to identify specific dynamic changes that follow ligand binding. The difference in deuterium labeling before, during and after ligand binding provides a highly structurally resolved measurement of dynamic reconfiguration during allosteric regulation. Equilibrium experiments reflect local minima of R/T-states, while non-equilibrium data reveal transition state ensembles between local minima on the energy landscape^44^. Here, we clearly identify amino acids that participate in transient conformational dynamics and we propose a structural pathway for those changes upon allosteric ligand binding. This approach stands to uniquely discover local structural dynamics and could be used broadly to identify such features in almost any ligand-induced allosteric transition of a protein.

## RESULTS

### Non-equilibrium allosteric activation/inhibition of GlyP

To observe the kinetically populated transition state of glycogen phosphorylase activation and inhibition, we first established the chemical conditions that fully activate or inhibit the enzyme. The catalytic activity for the fully active form phosphorylated at Ser14 (GlyPa) was used as a positive control reference in a glycogen hydrolysis colorimetric assay of GlyPb activation (**Figure 1A,E**; **Figure S2A)**. Against this level, the maximal extent of GlyPb allosteric activation by AMP was then established^26,40^. This showed that full activation is achieved with AMP ≥25 mM, the lowest concentration of AMP tested (range 25-100 mM), given no significant difference between the activity of GlyPb+AMP compared with GlyPa (**Figure 1A,E**). We also sought to verify the requirement of ammonium sulfate (AS) for full activation, known to mimic the presence of phosphate by binding GlyP at the AMP effector site, catalytic site and Ser14 phosphorylation site^20^. There are conflicting reports on the requirement of AS for full activation of GlyPb cooperatively with AMP: whilst GlyPb affinity for AMP is reported to increase 50-fold and activity to increase by 50% with sulfate^25,45-47^, here we found activation by sulfate alone was not achieved (15% of maximum), nor was a cooperative effect observed on activity (**Figure 1E,C** and **Figure S2A**). However, there was structural cooperativity between AMP and AS, as evidenced by concentration-dependent changes in hydrogen-exchange kinetics for amino acids site-localized to the nucleotide site and tower helix (**Figure 1D**). Local structural perturbations were observed for GlyPb samples supplemented with 0-100 mM AMP and/or 0-100 mM AS by measuring HDX-MS at equilibrium. Binding of AMP at the nucleotide site results in local protection against D-labeling that is positively cooperative with AS (**Figure 1D**). Corresponding changes were observed in the tower helix, which shows only weak protection with AS alone, but stronger protection with AMP (similar at all AMP concentrations tested) that increases with [AS] (**Figure 1D)**. The 380 loop shows cooperativity between AS and AMP, resulting in deprotection against D-labeling as the loop becomes disordered in the active state^20^ (**Figure 1D**). This identified cooperativity in GlyPb structural activation by AMP/AS that is not readily observed in the activity assay (**Figure 1D,E**)^25,40,48^. One-way ANOVA tests proved that higher concentrations of AMP or AS were not significant for either catalytic activity or structural perturbation. Therefore, the lowest concentrations tested of 25 mM AMP and 25 mM AS were considered to fully activate GlyPb catalytically and structurally and these conditions were used for the experiments to study non-equilibrium activation of GlyPb. Caffeine inhibits the active GlyPa R-state, stabilising the T-state, therefore we also sought to determine the minimal caffeine concentration required to fully inhibit 12 μM GlyPa – the concentration for subsequent non-equilibrium HDX-MS experiments. This was achieved with 32 mM caffeine, given by equivalence in specific activity to inactive GlyPb by ANOVA these conditions were used for the experiments to study non-equilibrium inhibition of GlyPa (**Figure 1F** and **Figure S2B**).

**Figure 1:**
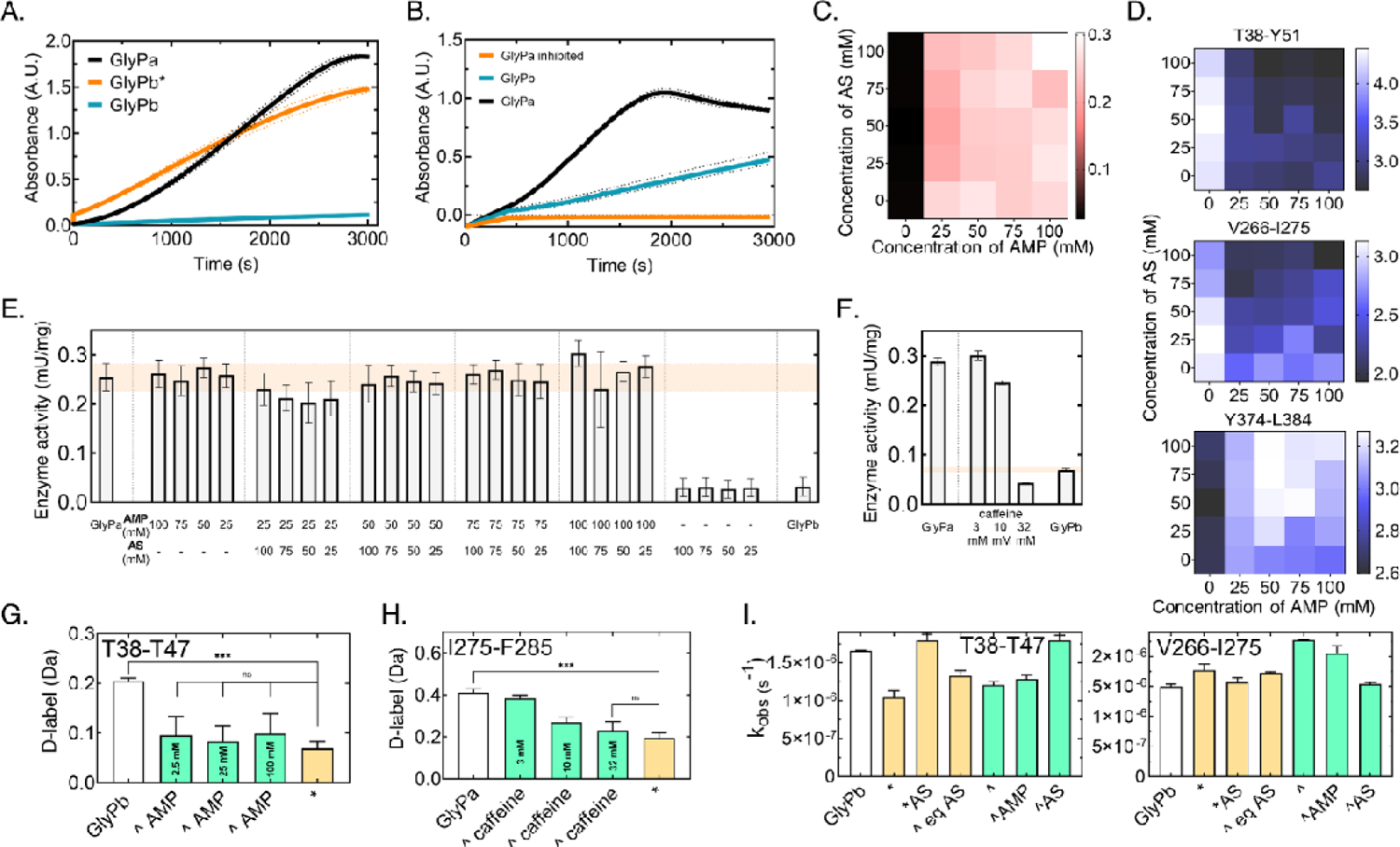
Allosteric activation of glycogen phosphorylase b and inhibition of glycogen phosphorylase a. **(A)** Glycogen hydrolysis kinetics for GlyPa (black), GlyPb (blue) and GlyPb* (orange; GlyPb equilibrated with 25 mM AMP and 25 mM AS). Absorbance (A.U.) at 30 °C for +60 min at 450 nm. Mean and SD from n=2. Plots for all conditions in Figure S2A. **(B)** Glycogen hydrolysis kinetics for GlyPa (black), GlyPb (blue) and GlyPa* (orange; GlyPa equilibrated with 32 mM caffeine. Absorbance (A.U.) at 30 °C for +60 min at 450 nm. Mean and SD from n=3. Plots for all conditions in Figure S2B. **(C)** Heat map of measured activity for combinations of AMP/AS concentrations in range 0-100 mM. Inactive GlyPb (black); highest activity achieved (white). Activity was calculated by a standard curve for n=2. **(D)** Heat maps for structural perturbations in the presence of AMP/AS, detected by equilibrium HDX-MS for the nucleotide site **(T38-Y51)**, tower helix **(V266-I275)** and 380s loop **(Y274-L384)**. Deuterium uptake data from the HDX screen was used to produce these heat maps, n=3. The relative deuteration level for each sample is color coded shown on the scale right; note different normalization. **(E)** Catalytic activity of GlyPb supplemented with AS and AMP. Shaded region – GlyPa 95% confidence intervals. **(F)** Catalytic activity of GlyPa supplemented with caffeine. Shaded region – GlyPb 95% confidence intervals. **(G)** AMP binding saturation during the dead time of the HDX labelling (50 ms). Non-equilibrium addition of 25 mM AMP/25 mM AS is optimal to fully saturate the nucleotide site. **(H)** Caffeine binding saturation during the dead time of the HDX labelling (50 ms). Non-equilibrium addition of 32 mM caffeine is optimal to fully saturate the inhibitor site. **(I)** Ammonium sulfate influence on the tower helix and nucleotide site. HDX labelling data was fitted and plotted versus the state (Figure S3). States annotated include: GlyPb -apo, * -equilibrium activated with 25 mM AMP and 25 mM AS, *AS -equilibrium activated with 25 mM AS, ^ eq AS – equilibrated with 25 mM AS and then non-equilibrium activated with AMP, ^ -non-equilibrium activated with 25 mM AMP and 25 mM AS, ^AMP – non-equilibrium activated with 25 mM AMP, ^AS – non-equilibrium activated with 25 mM AS.

In order to observe any kinetically populated transient structural ensembles, it is critical for the non-equilibrium experiments that the protein:ligand complex is fully saturated within 50 ms (dead time of the non-equilibrium experiment), governed by the second-order rate constant of association with units M^-1^s^-1^. While T-state GlyPb is allosterically activated by the binding of AMP at the nucleotide site, active phosphorylated R-state GlyPa is allosterically inhibited by the binding of caffeine between the 280s active site gate loop and the L40 (Tyr 613) loop. Therefore, the degree of AMP saturation at the nucleotide site was determined by protection against D-labeling at amino acids T38-T47, whereas the degree of caffeine saturation was determined by protection against D-labelling at amino acids N274-F285. In separate experiments, a range of AMP concentrations (2.5, 25 and 100 mM) and a range of caffeine concentrations (3, 10 and 32 mM) were assessed by rapid mixing with GlyPb or GlyPa, respectively, under non-equilibrium conditions (**Figure 1G-H)**. To confirm that a 1:1 complex with ligand had been reached, these were compared with the end-state of the respective process: activated GlyPb fully-equilibrated with 25 mM AMP, or inhibited GlyPa fully-equilibrated with caffeine. After 50 ms of incubation with ligand in deuterated buffer, 32 mM of caffeine achieved equivalent protection against D-labelling compared with the fully-equilibrated inhibited state (GlyPa + 32 mM caffeine) within the 50 ms dead-time of the HDX-MS experiment. The case of GlyPb activation was more complicated, as it was important to consider the contribution of both AMP and AS to ternary complex formation under these rapid-mixing conditions (**Figure S3**Error! Reference source not found.). Incubation of GlyPb with AS fully equilibrated (*AS) or at non-equilibrium (^AS) showed a lack of protection against D-labeling relative to GlyPb alone, signifying absence of binding to the putative sites for AS interaction. 25 mM AMP alone did not fully bind in 50 ms, whereas full protection is achieved within error for the non-eq addition of 25 mM AMP and AS. This confirms that the presence of AS enhances the *k*_*a*_ for GlyP:AMP association and these conditions yield full complex formation inside the dead time of the non-eq HDX-MS experiment. As we and others have recently reported, the presence of salts can significantly alter the intrinsic rates of HDX, thus masking – or artefactually creating – protein structural dynamics that are inferred by the HDX data^49^. This is not a systemic alteration of HDX kinetics by the presence of AS, but rather it is confirmed to be a local effect at the nucleotide site, as peptides covering other regions of the protein in the same HDX-MS data set do not show the same behavior: for example the tower helix (reported by peptide segment V266-I275) shows the opposite relationship with the *k*_*obs*_ for D-labeling being significantly faster in the presence of 25 mM AS/AMP under equilibrium and non-eq conditions (**Figure 1I, Figure S3**Error! Reference source not found.). We concluded that the ligand binding sites for inhibition and activation of GlyP by caffeine and AMP, respectively, are achieved within the dead-time of the HDX-MS experiment at times post-mixing >= 50 ms. Therefore, we proceeded to evaluate the transient structural kinetics that occur upon allosteric regulation across the whole enzyme.

### Transient structural dynamics during GlyPb allosteric activation by AMP

To identify specific dynamic changes associated with long-range allosteric communication induced by ligand binding, we developed a non-equilibrium approach to millisecond hydrogen/deuterium-exchange that reveals the local structural perturbations that result from the conditions established above (**Figure S1**). As the D-labeling of polypeptide backbone amide groups is a sensitive, but convoluted measure of H-bonding and solvent accessibility, first we sought to categorize the local differences in the protein ensemble during the non-equilibrium phase following allosteric activation by AMP/AS. Subtle changes in dynamics and structural changes can be derived from HDX-MS data, reflected in the observed kinetics of deuterium uptake plots at specific positions within the protein, given by short peptide segments in the ‘bottom-up’ experiments used here^44^. The data comprised measurements at 31 D-labeling time points over four orders of magnitude from 50 ms – 300 s for 219 peptide segments that cover 90.5% of the GlyP amino acid sequence (**Figure S4A**). This was done for three conditions: (i) inactive GlyPb (apo), (ii) fully activated GlyPb (termed GlyPb*) equilibrated for one hour with 25 mM AMP, 25 mM AS and (iii) GlyPb activated at non-equilibrium by rapid mixing with 25 mM AMP, 25 mM AS (termed GlyPb^). It is clear that for many peptides in a variety of regions of the protein the D-labeling kinetics under non-eq conditions are qualitatively different from both the apo and the activated states (**Figure S5-6** and **Figure S16**). There are peptides for which the D-labeling values fall outside of the range measured for either the initial (apo) or equilibrated active (GlyPb*) forms, which indicates a degree of non-linearity in the structural interpolation (**Figure 3, Figure S16;17**). However, the sub-molecular location, magnitude and nature of relative (de)protection of these show a variety of differences. Therefore, we next sought to classify these structural dynamic changes occurring during allosteric activation to better identify clear behaviors in each part of the enzyme.

To categorize the local structural changes that occur during non-equilibrium activation/inhibition into correlated transient dynamics, we developed a simple, but robust quantitative analysis of the HDX-MS data that served to represent the non-equilibrium HDX kinetics for the pool of peptide segments and to act as input to a clustering algorithm^50,51^. Each amino acid of the protein was represented spatially in a manner that reflects the relationship of its non-equilibrium state to the starting and ending states of the protein (**Figure 2**). We identified the archetypes of non-equilibrium structural kinetics that might be possible to observe in this plot (**Figure S9)**. In order to categorize the continuum of diverse D-labeling kinetic differences that we observed, we performed a k-means clustering analysis to identify amino acids with correlated HDX kinetics. Seven clusters of amino acids within GlyP comprises a common structural response to activation by AMP/AS (**Figure 2, Figure S10**). **Figure 3A** shows the D-labeling uptake plot for representative amino acids from each cluster, alongside the structural location of the cluster.

**Figure 2:**
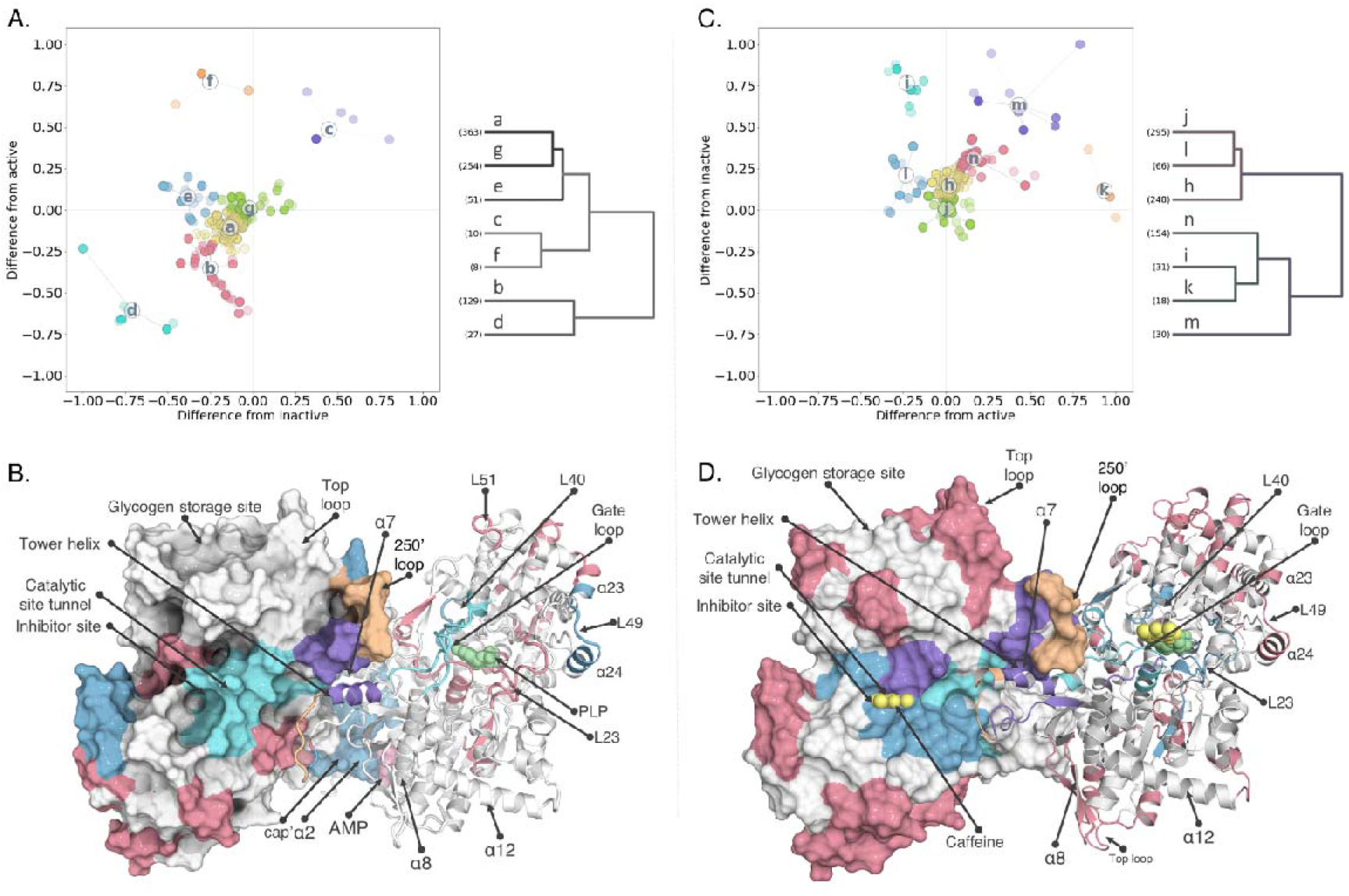
Clusters of structural kinetic behavior in GlyP. **(A)** The sum difference of the D uptake per amino acid between non-equilibrium and GlyPb/active state plotted against x/y axes, respectively. Amino acids were grouped into seven clusters (a-g) resulting from a k-means hierarchical analysis. The centroid of each cluster is denoted by the letter connected to each member by an edge. Raw D-labelling uptake data was centroided, corrected for maximum exchangeable amide hydrogens, normalized and the different states were pairwise subtracted. Dendrogram of clustering in A is shown on the right. Y-axis is Euclidean distance between the clusters (arbitrary units). The number of amino acid members in each cluster is indicated at the bottom of each cluster (truncated for clarity; Figure S15A). **(B)** Clusters (excluding g and a) represented on GlyPb active state structure (PDB:3E3N). Notable features are indicated on the left subunit molecular surface; Notable secondary structural elements indicated. **(C)** The sum difference of the D uptake per amino acid between non-equilibrium and GlyPa/inactive state plotted against x/y axes, respectively. Amino acids were grouped into seven clusters (h-n) resulting from a k-means hierarchical analysis. The centroid of each cluster is denoted by the letter connected to each member by an edge. Raw D-labelling uptake data was centroided, corrected for maximum exchangeable amide hydrogens, normalized and the different states were pairwise subtracted. Dendrogram of clustering in C is shown on the right. Y-axis is Euclidean distance between the clusters (arbitrary units). The number of amino acid members in each cluster is indicated at the bottom of each cluster (truncated for clarity; Figure S15B). **(D)** Clusters (excluding h and j) represented on GlyPa inactive state structure (PDB:1GFZ). Notable features are indicated on the left subunit molecular surface; Notable secondary structural elements indicated.

The hierarchical relationship between the seven clusters was determined based on the observed transient structural changes (**Figure 2A**)^52^. This reveals that there are two closely related clusters (**a** and **g)** that show little or no observable transient structural change during allosteric activation (i.e. close to the origin on **Figure 2A** and **Figure SI15**). We calculated the local hydrogen exchange rate (*k*_*obs*_) by fitting HDX-MS data to a stretched exponential model^49^. For most peptides, *k*_*obs*_ was low (k_obs_ < 1 × 10^−5^ s^-1^ for 130 of 219 peptides measured) in all three conditions (Error! Reference source not found.**Table S2** and **Table S3**), as GlyP is highly structured with many amino acids buried in the core of the enzyme with low solvent accessible surface area (SASA) and stable H-bond networks, consistent with previous studies^20,30,49,53-56^. This validates that much of the enzyme is structurally identical before, during and after allosteric activation, **Figure S5-6** and **Figure S16**. A third related cluster (**e**), representing the nucleotide binding site for AMP, shows no deviation from the active state and only transient relative protection compared to the inactive state that is on-pathway from the apo-to AMP-bound state (i.e. on y=0 in **Figure 2A**). This group includes residues from α2 (residues 47-78), the cap’ region (residues 38-47) and residues from the nucleotide binding domain (**Figure 3A** and **Figure S16**). The α8 helix is also essential for AMP binding, but unfortunately our coverage was lacking peptides in that region. There are two closely related clusters which represent the 250’ loop (**f**) and the tower helix (**c**) that show transiently increased D-labeling compared to one or both equilibrium states. There are also two related clusters, representing the pyridoxal phosphate cofactor binding residues (**b**) and the 606 loop that are distinct from all others which show protection against D-labeling compared to both the inactive and the active state. The amino acids surrounding the PLP cofactor retain apo GlyPb HDX kinetics, indicating either that this region has a structural transition with a low probability or that the induced protection against HDX occurs too slowly to prevent the observed labeling.

**Figure 3:**
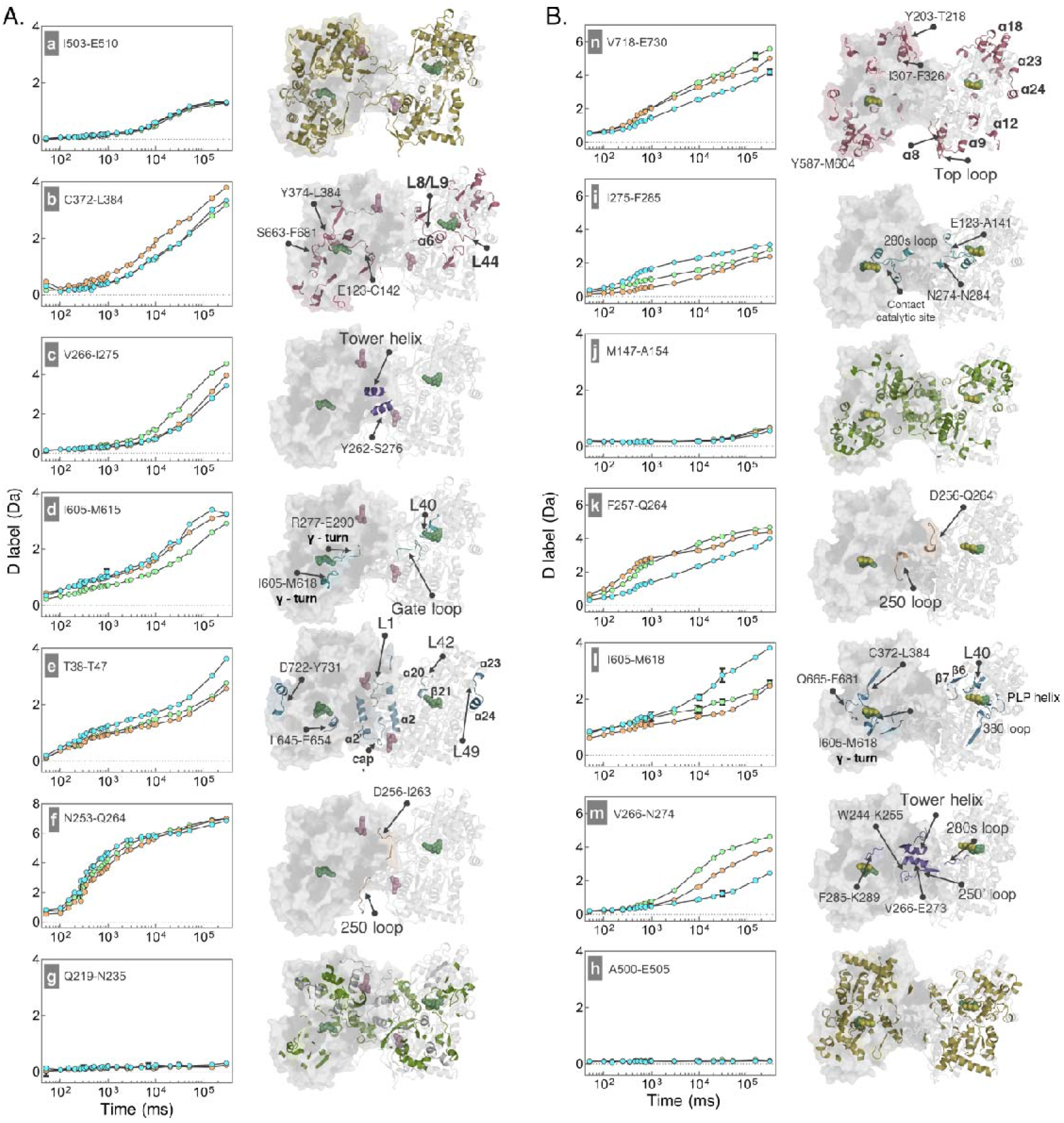
Local structural kinetics upon allosteric activation and inhibition revealed by non-equilibrium D-labeling. **(A)** Hydrogen exchange kinetics of representative peptide segments for each cluster. Mean of n=3; error bars mostly contained within data points. GlyPb (blue); fully equilibrated activated with 25 mM AMP and 25 mM AS (orange); non-equilibrium activated with 25 mM AMP and 25 mM AS (green). **(B)** Clusters mapped on the crystal structure of inactive GlyPb (3E3N.pdb). Detailed annotation is represented on each structure per cluster. **(C)** Hydrogen exchange kinetics of representative peptide segments for each cluster. Mean of n=3; error bars mostly contained within data points. GlyPa (blue); fully equilibrated inactivated with 32 mM caffeine (orange); non-equilibrium inactivated with 32 mM caffeine (green). **(D)** Clusters mapped on the crystal structure of inactive GlyPa (1GFZ.pdb). Detailed annotation is represented on each structure per cluster.

A feature of the HDX-MS data during non-equilibrium AMP activation is that there are transient structural changes that are distinct from both the inactive/active states (i.e. amino acids that do not lie on either x=0 or y=0 in **Figure 2A**) and therefore are not represented by high-resolution structural models derived from the equilibrium states. Activation of GlyPb by AMP binding to the nucleotide site induces the well-established structural changes in the 250’ and 280s loops, together with rotation of the tower helix (α7) (**Figure S12**). The non-equilibrium HDX-MS reveals previously unobserved transient changes that facilitate this. The 250’ loop (A248-G260) exhibits unique behavior during allosteric activation (cluster **f** in **Figure 3A)**. The D-labeling kinetics interpolate smoothly between the GlyPb state (at t=50 ms) and the active GlyPb* state (at t=10 s). This indicates a likely two-state process for helical lengthening and that a gain of structure has fully equilibrated in the molecular population by 10 s of D-labeling, consistent with the tower helix in the active state lengthening at the N-terminal end by one turn in the GlyPb* state (3E3N.pdb^30^). Clusters **c** (the tower helix) and **d** (the 606 loop) show opposite structural behaviors: the 606 loop shows transient protection (increased H-bonding; reduced solvent accessibility), whereas the tower helix shows transient deprotection (i.e. local unfolding). The tower helix (G261-S276) is a contiguous alpha helical structure in all known experimental models of apo, active and inactive forms of GlyP. However, unexpectedly here we observe that it transiently loses protection against hydrogen/exchange, with increased observed D-labeling during allosteric activation following AMP binding at the nucleotide site (Error! Reference source not found.**A** and **Figure S12**).

We sought an atomistic rationale for this transient unfolding, so an all-atom interpolation method was used to generate structures on a non-linear, energy minimized trajectory between apo GlyPb and GlyPb*. The Climber method has been shown to sample intermediate structures effectively, though has been difficult to validate until now, owing to the experimental challenge of determining transient local structural changes^57^. Here a trajectory was created which interpolates between two crystal models that have intact tower helices with predicted hydrogen-bonds in conventional geometries, yet several of these hydrogen-bonds are shown to undergo dynamic remodeling as the helices refold and rotate by ∼40° relative to each other to adopt the established active R-state, GlyPb* (**Figure 4A** and **Figure S12**; **Supplementary movies**)^58^. The interpolation is consistent with the non-equilibrium HDX-MS data in which there is relative deprotection compared to both GlyPb and GlyPb* precisely at I263-D268, resolved by multiple overlapping peptide segments in the data, supporting an interpretation that there is a transient unfolding in the tower helix during allosteric activation.

**Figure 4:**
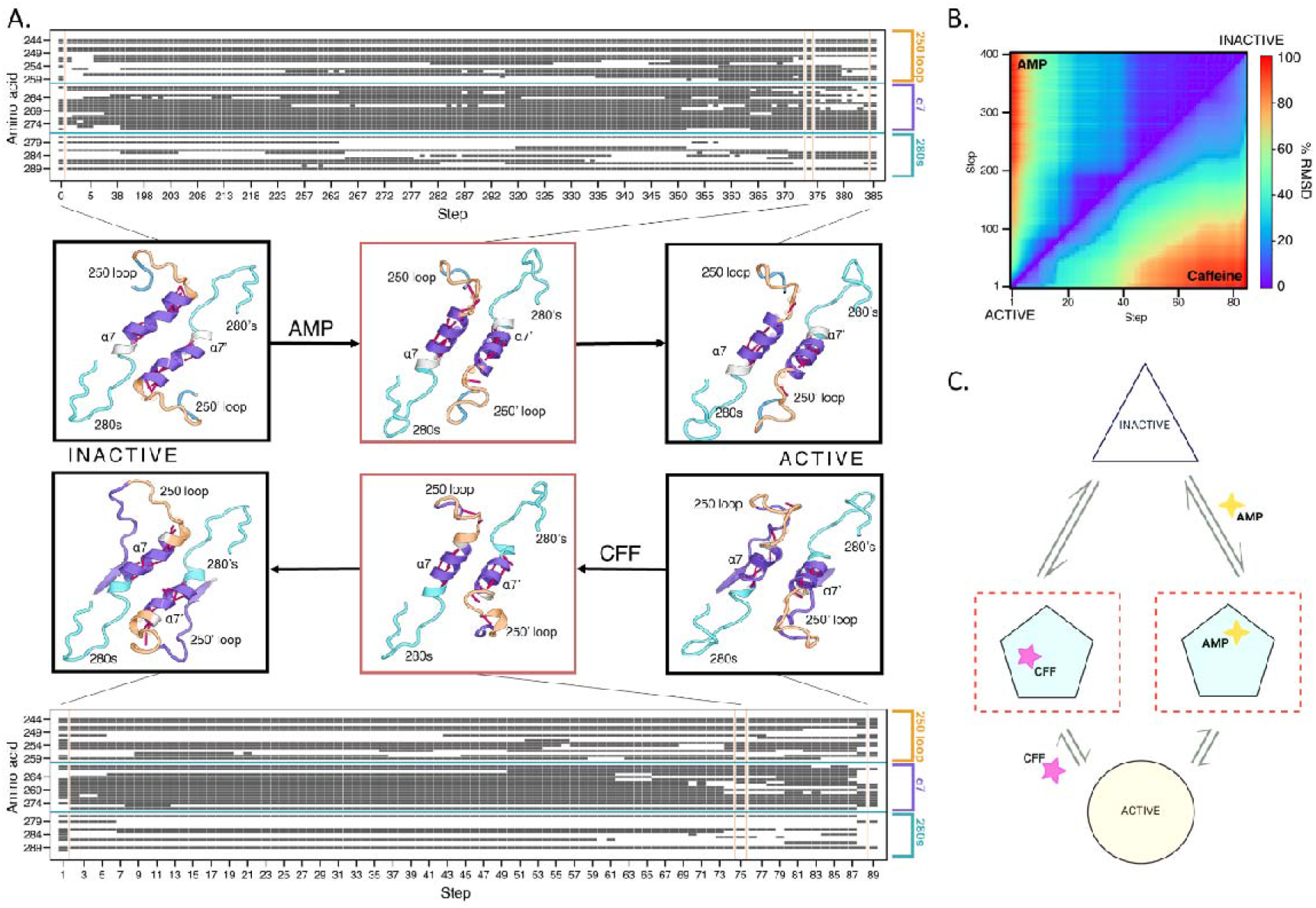
Structural interpolation between (in)active states of GlyP. **(A)** Tower helix and adjacent loops in inactive (left panel), transitional (middle) and active (right) conformations. Transitional structures with multiple broken hydrogen-bonds (pink lines) in the α7 helix are observed upon interpolation between apo GlyPb – GlyPb-AMP (activation) and between apo GlyPa -GlyPa-caffeine (inhibition). Representative structure shown; full trajectories provided as supplementary files. Bottom and top heat maps represent the hydrogen-bond network of each structure upon activation (top) and inhibition (bottom) in this region of the enzyme, gray: H-bond; white: no H-bond. **(B)** Interpolation trajectory represented as rmsd between each structure. Calculated only for the tower helix region (amino acids 256:288). Normalized to maximum values from the AMP-activation (lower triangle) and caffeine-inhibition (upper triangle), respectively. **(C)** Proposed model of the allosteric modification of GlyP. Both modifications, activation (AMP) and inhibition (CFF) of GlyPb and GlyPa, respectively, undergo similar transition mechanism, with common structural features and proximal to the active state.

### Transient structural dynamics during GlyPa allosteric inhibition by caffeine

We next considered whether this transient unfolding of the tower helix seen during activation is a feature common to the GlyP inhibition pathway. GlyPa phosphorylated at serine14 is basally active but is itself allosterically inhibited by binding of caffeine, which stabilizes a T-state that is similar to the GlyPb apo state. The caffeine binding site is formed between the 280s and 606 loops, 3.5 nm from the nucleotide site where AMP binding stabilizes the reverse transition to the R-state. Therefore, we next sought to identify the transient structural features on the inhibition pathway by performing non-equilibrium HDX-MS of GlyPa in the presence of caffeine. As for the AMP-activation experiments, this comprised measurements at 20 D-labeling time points 50 ms – 300 s resulting in 171 peptide segments that cover 91.4% of the GlyP amino acid sequence (Error! Reference source not found.**B**). This was done for three conditions: (i) active GlyPa (apo), (ii) fully inhibited GlyPa (termed GlyPa’) equilibrated for one hour with 32 mM caffeine and (iii) GlyPa inhibited at non-equilibrium by rapid mixing with 32 mM caffeine (termed GlyPa^).

From a comparison of the observed HDX kinetics between GlyPa^ and each of the two equilibrated states, it is clear that the inhibition pathway involves a number of amino acids that show a transient structure distinct from both the start/end states (**Figure 2C-D; Figure S7-8**). Based on this, a clustering analysis identified amino acids with correlated HDX kinetics. As for AMP-activation, caffeine-inhibition resulted in seven clusters (**Figure S11;15**) of amino acids within GlyP with a common structural response to ligand (**Figure 2C; Figure 3B**). Again, much of the protein is relatively unchanged before, during and after allosteric inhibition (clusters **h** and **j, Figure 2C**), which is consistent with the available crystal structures. The 280s and 606 loops form the binding site for caffeine which intercalates between them in the GlyPa T-state crystal model. These amino acids are co-located in cluster **l** which shows HDX equivalent to the GlyPa’ and protected compared to the GlyPa states, consistent with direct binding at this site without any observed transitional behavior (cluster **l, Figure 2C-D;Figure 3B**). During caffeine-inhibition these three clusters (**h/j/l**) have minimal transient structural features and are found to be hierarchically unrelated to all other amino acids in the other four clusters (**Figure 2C**). Furthermore, the 250’ loop is highly deprotected against hydrogen-exchange and labels more than the initial GlyPa state, but similarly to the inhibited GlyPa’ state (cluster **k**). This is consistent with an interpretation that it is a two-state process, where the caffeine-inhibited T-state is observed immediately upon binding, in which one turn of the tower helix unwinds to extend the 250’ loop, as seen in the crystal models. The tower helix itself (cluster **m**) is labeled more with deuterium under non-equilibrium conditions (GlyPa^) than in either GlyPa or GlyPa’ states. As this is a highly solvent-exposed alpha helix in the initial R-state, it is most consistent with an interpretation of transient unfolding (**Figure S13**), again resolved by overlapping peptide segments to be most pronounced at amino acids L254-L271. The 280s loop shows a diverged behavior, with the more C-terminal part forming part of the direct ligand binding site (cluster **l**) and with the N-terminal stretch showing transiently reduced hydrogen-exchange compared to the initial apo GlyPa state but increased when compared to GlyPa’. This is consistent with the initial binding of caffeine to F285 and a subsequent propagation of induced helical structure further upstream in the 280s loop as the tower helix lengthens by one turn at the C-terminal end.

It is apparent that a number of highly resolved transient structurally dynamic features are observed by non-equilibrium HDX-MS upon allosteric inhibition of GlyP by caffeine.

### Transient unfolding of the tower helix is common to allosteric activation and inhibition pathways

We next sought to compare the transient structural features observed upon AMP-activation and caffeine-inhibition to determine whether these different ligands (and different allosteric sites) appear to act via different pathways, or the same one.

The 250’ and 280s loops show opposite neHDX-MS behaviour in response to AMP and caffeine, which is logical given that these motifs are structurally different in the R/T-states and necessarily interpolate between them. However, there is an unexpected common feature during both allosteric activation and inhibition with transiently increased HDX observed in the tower helix that sits in between these two loops. Non-linear interpolation between the atomistic models for GlyPa R-state (apo) and caffeine-bound (T-state) also predicts breakage, then reforming, of hydrogen-bonds in the tower helix, as it does for AMP-activation (**Figure 4A**). It also predicts that the two structural pathways are closely related with low rmsd in the 250’ loop, tower helix and 280s loop between structures close to the T-state and much larger differences in rmsd close to the R-state (**Figure 4B**). As the tower helix is highly solvent-exposed, the transiently lower protection factor would likely be the result of H-bond breakage which is known to be a major determinant of hydrogen-exchange rate. This is suggested by the structural interpolation, which predicts these hydrogen-bonds in the tower helix to be volatile in both pathways (**Figure 4A**).

These are the major transient structural features observed by non-equilibrium HDX-MS and appear common to both pathways, with a consistent atomistic rationalization given by the Climber structural interpolations.

## DISCUSSION

### Transient features of allosteric pathways revealed

Here we show that allosteric regulation of the central metabolic enzyme, glycogen phosphorylase, involves a number of dynamic changes in local structure that are not observed in static structural models of the R- and T-states. We established a new protocol exploiting millisecond hydrogen/deuterium-exchange at non-equilibrium to characterize transient changes in protection in response to ligand, resolved at near amino acid level throughout a 194 kDa enzyme. This revealed that AMP-activation induces a transient increase in HDX in the tower helix. We rationalized this by interpolating the known structures for this allosteric transition using an atomistic method based on the Energy Calculation and Dynamics (ENCAD) molecular-mechanics force-field, Climber. This predicts that the well-established lengthening and 40° relative rotation of the tower helices by first shortening, then bending it, wholly consistent with the increased hydrogen-exchange observed during the process. Unexpectedly, caffeine-inhibition – binding to a different allosteric site 3.5 nm away from AMP -also resulted in a transient increase in HDX in the tower helix. The structural interpolation predicts that this is due to a similar extent of transient breaking of hydrogen-bonds in the tower helix (**Figure 4A**). So, both experiments are consistent with the interpretation suggested by the simulation that breaking of the tower helix is a common requirement of both AMP-activation and caffeine-inhibition pathways (**Figure 4B-C**).

### Slow (millisecond) labeling can observe transient structural dynamics of the molecular population

It is perhaps unexpected that differences in non-equilibrium HDX-MS kinetics were observed across a range of time domains from milliseconds to seconds. Structural kinetics were observed even at the fastest labeling time of 50 ms (**Figure S5;7**), in several significant regions, including: 250 □ turn loop, β3 and α6, 280s loop and another □ turn with the central residue Tyr 613 said to be forming a hydrophobic sandwich of the nucleotide inhibitor site^59^. At slower deuterium labeling times, significant changes in protection between the three states was observed throughout the α7 tower helix (residues 261-274), the β11b (residues 276-279), α23 (residues 714-725) and α24 (residues 728-735) (**Figure S6;8**). We interpret this range of observed non-equilibrium structural kinetics to indicate that the probability of the local structural changes is very low (i.e. bounded by a high energy transition state), resulting in observation of the populated transient species (presumably transition states), even milliseconds after ligand binding. The wide range of local protection factors against hydrogen-exchange (due to existence of stable – or transient -local structure) would then effectively stretch the observation of structural changes across multiple orders of magnitude, as seen here.

### Hydrogen/deuterium-exchange may directly observe vibrational energy transfer upon ligand binding

The HDX-MS intrinsic rate constants indicate a half time of 100’s of milliseconds for which a protein structural state would need to exist in order for it to be appreciably detectable by this technique^60^. However, our non-equilibrium HDX-MS data appear to observe transient species whose lifetime would normally be considerably shorter than this. One explanation for this is that the vibrational energy transfer (VET), derived from the enthalpy of ligand binding, is anisotropically focused into a ‘hotspot’ which can undergo the HDX reaction faster than would be predicted because of an elevated local temperature. This would agree with our simulated trajectories where hydrogen bond stability is locally reduced in the ‘hotspot’ motif and with recent experimental evidence that VET is mediated by hydrogen-bond networks^61^ and with coarse-grained models of specific energy dissipation pathways in proteins^62^.

We believe that this approach, while critically important to establish rapid saturation of the ligand binding site, provides a straightforward way to directly observe allosteric structural dynamics at non-equilibrium which can be applied broadly to proteins irrespective of size or stability and with, in principle, any chemical or protein ligand. In this manner, we expect it to be important in future to understand not only what has changed in protein structure, but how the changes come about and propagate through the molecule.

## METHODS

### Sample preparation

Glycogen phosphorylase b and glycogen phosphorylase a were dissolved in phosphate buffered saline (PBS), pH 7.4 to 10.3 pmol/µL. For the equilibrium and non-equilibrium activated samples, the required amount of AMP/AS and caffeine were added, in the stock solution or the labelling buffer, respectively. During the screening experiments (HDX screen and enzyme activity assay) each sample was prepared simultaneously, with 10.3 pmol/µL GlyPb/GlyPa and with varying concentrations of AMP, AS or caffeine, **Error! Reference source not found**..

### Screening of conditions to activate GlyP by enzyme activity assay

The enzyme activity of GlyPb and GlyPa was screened for AMP/AS and caffeine dependence**Error! Reference source not found**.. A colorimetric assay kit was used (ab273271), with only one adjustment of the protocol for GlyP sample preparation. A suitable protein concentration was determined by pre-testing a range of concentrations and plate reader settings, then all 25 samples were prepared in parallel at 1 mg/mL GlyP (**Error! Reference source not found**.). Three relevant concentrations of caffeine were screened (3, 10 and 32 mM) to determine the extent of inhibition. Reaction kinetic curves and specific activity (mU/mG) were plotted and calculated according to the suggested protocol.

### Screening of conditions to activate GlyP by HDX-MS

The observable perturbations in HDX were screened for AMP/AS dependence. A CTC PAL sample handling robot (LEAP Technologies, USA) was used to mix samples (**Error! Reference source not found**.) with deuterated PBS buffer in 1:20 ratio, then quench the labeling reaction by 1:1 mixing with 100 mM potassium phosphate to a final pH of 2.55 at 0 °C. Samples were immediately digested online with an Enzymate immobilized pepsin column (Waters) at 12 °C, the derived peptides trapped on a VanGuard 2.1 × 5 mm ACQUITY BEH C18 column (Waters) for 3 minutes at 125 µL/min at 0.5 °C and separated on a 1 × 100 mm ACQUITY BEH 1.7 μm C18 column (Waters) with a 7 min linear gradient of acetonitrile (5-40%) supplemented with 0.1% formic acid. Peptides were eluted into a Synapt G2-Si mass spectrometer (Waters).

### Ammonium sulfate influence and maximum velocity kinetics

The maximum velocity kinetics and the AS influence on the HDX were determined prior continuing with the non-equilibrium experiments. HDX labelling experiments were performed at 11 time points, ranging from 50 ms to 300 s. AMP activation was tested at 3 different concentrations, including 2.5, 25 and 100 mM under non-equilibrium conditions with 25 mM AS. The AS influence on the equilibrated sample and the pre-equilibrated AS sample, was also determined. Analyzed protein states include: GlyPb -apo, * -equilibrium activated with 25 mM AMP and 25 mM AS, *AS -equilibrium activated with 25 mM AS, ^ eq AS – equilibrated with 25 mM AS and then non-equilibrium activated with AMP, ^ -non-equilibrium activated with 25 mM AMP and 25 mM AS, ^AMP – non-equilibrium activated with 25 mM AMP, ^AS – non-equilibrium activated with 25 mM AS.

### Hydrogen-deuterium exchange mass spectrometry

HDX was performed using a fully-automated, millisecond HDX labelling and online quench-flow instrument, ms2min (Applied Photophysics Ltd), connected to an HDX manager (Waters). During the labelling experiments, 14 µL of either apo or equilibrated GlyPb/GlyPa were delivered to the labelling mixer. A 20-fold dilution with labelling buffer at 20°C commenced the HDX. Depending on the desired experiment, the labelling buffer differed significantly. For apo and equilibrium experiments, the labelling buffer was 1 x PBS, pHread =7.00 at 20°C. During non-equilibrium experiments and other assessments (see Error! Reference source not found.), the labelling buffer consisted of 1 x PBS, pH =7.00 at 20°C combined with 25 mM AMP/25 mM AS or 32 mM caffeine. All GlyPb samples were labelled at 30 time points, ranging from 50 ms to 300 s. All GlyPa samples were labelled at 20 time points, ranging from 50 ms to 300 s. Immediately after, the HDX reaction was quenched by mixing with quench buffer (100 mM Potassium phosphate, pH =2.5 at 0°C) at 1:1 ratio, and sample digested online with an Enzymate immobilized pepsin column (Waters), the derived peptides trapped on a VanGuard 2.1 × 5 mm ACQUITY BEH C18 column (Waters) for 3 minutes at 125 µL/min and separated on a 1 × 100mm ACQUITY BEH 1.7 μm C18 column (Waters) with a 7-minute linear gradient of acetonitrile (5-40%) supplemented with 0.1% formic acid. Peptides were eluted into a Synapt G2-Si mass spectrometer (Waters). Mass spectra were obtained using the Waters HDMS^E^ mode within a mass range of 50 to 2000 m/z. Instrument parameters were configured as follows: a capillary voltage of 3.0 kV, a cone voltage of 50 V, a trap collision energy of 4 V, a traveling wave ion mobility separation at a velocity of 475 m/s, a wave amplitude of 36.5 V, and a nitrogen pressure of 2.75 mbar. For low-energy scans, a transfer collision energy of 4 V was applied, while high-energy scans utilized four separate collision energy ramps ranging from 15 to 55 V.

### Data and statistical analysis

PLGS (ProteinLynx Global Server 2.5.1, Waters) was used for the analysis of MS^E^ reference data and identification of all discoverable peptic peptides. DynamX 3.0 (Waters, USA) was used for processing and assignments of isotopic distributions of all raw data files. All HDX-MS experiments were performed in triplicate (technical replicates). The means and standard deviations (SD) of these replicates were used to determine the significance of changes, global significance threshold, as previously described^56^. In-house developed scripts in Python 3.4.0 (Python Software Foundation) were used for all post-processing analyses, clustering analyses, and plotting in PyMOL (Schrödinger, Inc.).

A clustering analysis was employed to investigate the behavior of the peptide during non-equilibrium activation/inhibition. The D-labeling data for each peptide segment was normalized to the theoretical maximum, to account for varying peptide length and presence of proline for which there is no signal. The sum of deuterium uptake over all time points for each peptide was calculated for all three protein conditions (states) of GlyPa or GlyPb, in each case. The values for the non-eq data were then subtracted from both other states for every peptide segment in the data set, resulting in an [n x 2] vector of 2-d coordinates. With the non-equilibrium measurements defined as 2-d coordinates relative to a start/end state, this constitutes a well-formed input for a k-means clustering analysis. The 2-d coordinates for HDX-MS differences were then *k-means* clustered with *k* clusters indicated to be optimal by both an elbow method and a Silhouette analysis.

### Non-linear atomistic structural interpolation

The Climber program was utilized for stepwise and non-linear structural interpolation between the active and inactive conformations using default paramters. This involved dynamic evaluation of RMSD between backbone and side chain atom pairs, with continual updates. Atom pair sets were created for each generated conformation between the initial (A) and final (B) states, based on criteria of distance between alpha carbons (>10 Å) and side chain atom pairs (<10 Å). Harmonic restraints were applied to steer inter-residue distances towards the target conformation. The force constant for conformational transformation was adjusted based on RMSD progress. Morphed conformations underwent iterative minimization using the ENCAD potential energy function. Initial coordinates for the allosteric activation and inhibition were taken from crystallographic structures of glycogen phosphorylase b and glycogen phosphorylase a. PDB ID 1gpb and 3e3n were used for GlyPb transition, and PDB ID 1gpa and 1gfz for GlyP inhibition. Missing residues (251-259, 315-324 in 1gfz; 251-260, 281-287 in 3e3n) were built and optimized using MODELLER (version 10.4). For each loop region, a total of 100 conformations were generated, and the model exhibiting the lowest value for MODELLER’s^63,64^ objective was chosen as the final protein structure.

### Graph network analysis of protein trajectories

The n PDB files output by Climber were compared recursively with each other to calculate [n x n] pairwise rmsd values. For plotting, the desired amino acids (i.e. the 250’ loop – tower helix – 280s loop) were extracted beforehand and the calculation was run only on those and the values normalised to the observed maximum. Plotting of the rmsd trajectory was done in Matplotlib. A graph network was constructed by selecting ten pdb files equidistant on the trajectory (with respect to number of steps taken to complete interpolation in Climber) and a Spring_layout model was used to plot the 2-d projection with the network Python library.

## DATA AVAILABILITY STATEMENT

The authors declare that the data supporting the findings of this study are available in this paper and its supplementary information files. All additional data is available upon reasonable request. Source data are provided with this paper. All mass spectrometry .raw files will be available from the PRIDE repository [accession pending].

## ACKNOWLEDGEMENTS

J.J.P. and M.K. are supported by a UKRI Future Leaders Fellowship [Grant Number MR/T02223X/1]. DI is funded by a BBSRC SWBioDTP studentship. For the purpose of open access, the author has applied a CC BY public copyright license to any Author Accepted Manuscript version arising from this submission.

## AUTHOR INFORMATION

### AUTHORS AND AFFILIATIONS

#### Living Systems Institute, Department of Biosciences, University of Exeter, Stocker Road, Exeter, EX4 4QD, UK

Monika Kish, Dylan P. Ivory, Jonathan J. Phillips

#### Alan Turing Institute, British Library, London, NW1 2DB, UK

Jonathan J. Phillips

## CONTRIBUTIONS

JJP conceived the study. MK and JJP designed the experiments and performed computational structural interpolations. MK, DI and JJP performed mass spectrometry experiments. MK and DI performed spectrophotometry assays, constructed atomistic models. MK performed clustering analysis, analyzed mass spectrometry data, and performed data and statistical analysis. All authors wrote the manuscript.

## CODE AVAILABILITY STATEMENT

This study uses in-house developed scripts in Python. Available upon reasonable request.

