## Supplementary Information for "Allosteric activation and inhibition of glycogen phosphorylase share common transient structural features"

### Table of figures:

Figure S1. Scheme for the continuous non-equilibrium HDX-MS approach. ____ 2
Figure S2: Activation/inhibition of Glycogen phosphorylase ____ 3
Figure S3. Ammonium sulfate influence on the binding site residues 38-47. ____ 3
Figure S4. Sequence coverage of glycogen phosphorylase from rabbit muscle. ____ 4
Figure S5. State D uptake difference maps for fast mixing times (50, 100, 150, 300 and 500 ms). ____ 5
Figure S6. State D uptake difference maps for slow mixing times (1, 3, 10, 30 and 300 s). ____ 6
Figure S7. State D uptake difference maps for fast mixing times (50, 100, 150, 300 and 500 ms). ____ 7
Figure S8. State D uptake difference maps for fast mixing times (1, 3, 10, 30 and 300 s). ____ 8
Figure S9. Theoretical behaviors of peptide segments upon non-equilibrium activation. ____ 9
Figure S10. Assessing the quality of k-means clusters fit on the data of GlyPb. ____ 10
Figure S11. Assessing the quality of k-means clusters fit on the data of GlyPa. ____ 11
Figure S12. Rotation and transition of the tower helix upon AMP activation of GlyPb. ____ 12
Figure S13. Rotation and transition of the tower helix upon caffeine inhibition of GlyPa. ____ 13
Figure S14. Representation of the full transition upon activation/inhibition. ____ 14
Figure S15. Full dendrogram of hierarchical clustering. ____ 15
Figure S16. HDX uptake curves of GlyPb per peptide. ____ 16
Figure S17. HDX uptake curves of GlyPa per peptide. ____ 35

### Table of tables:

Table S1. Screening varying activating concentrations of AMP and Ammonium Sulfate. ____ 2
Table S2. Local hydrogen exchange rate ($k_{obs}$) of GlyPb peptides. ____ 50
Table S3. Local hydrogen exchange rate ($k_{obs}$) of GlyPa peptides. ____ 53

Table S1. Screening varying activating concentrations of AMP and Ammonium Sulfate. In total 25 samples were analyzed plus GlyPa, as an activated control. Both an enzyme activity kit and HDX were employed for the screen.

| | | | | | |
|---|---|---|---|---|---|
| $(NH_4)_2SO_4$ (mM) | 100 | ✓ | ✓ | ✓ | ✓ |
| | 75 | ✓ | ✓ | ✓ | ✓ |
| | 50 | ✓ | ✓ | ✓ | ✓ |
| | 25 | ✓ | ✓ | ✓ | ✓ |
| | 0 | 25 | 50 | 75 | 100 |
| AMP (mM) | | | | | |

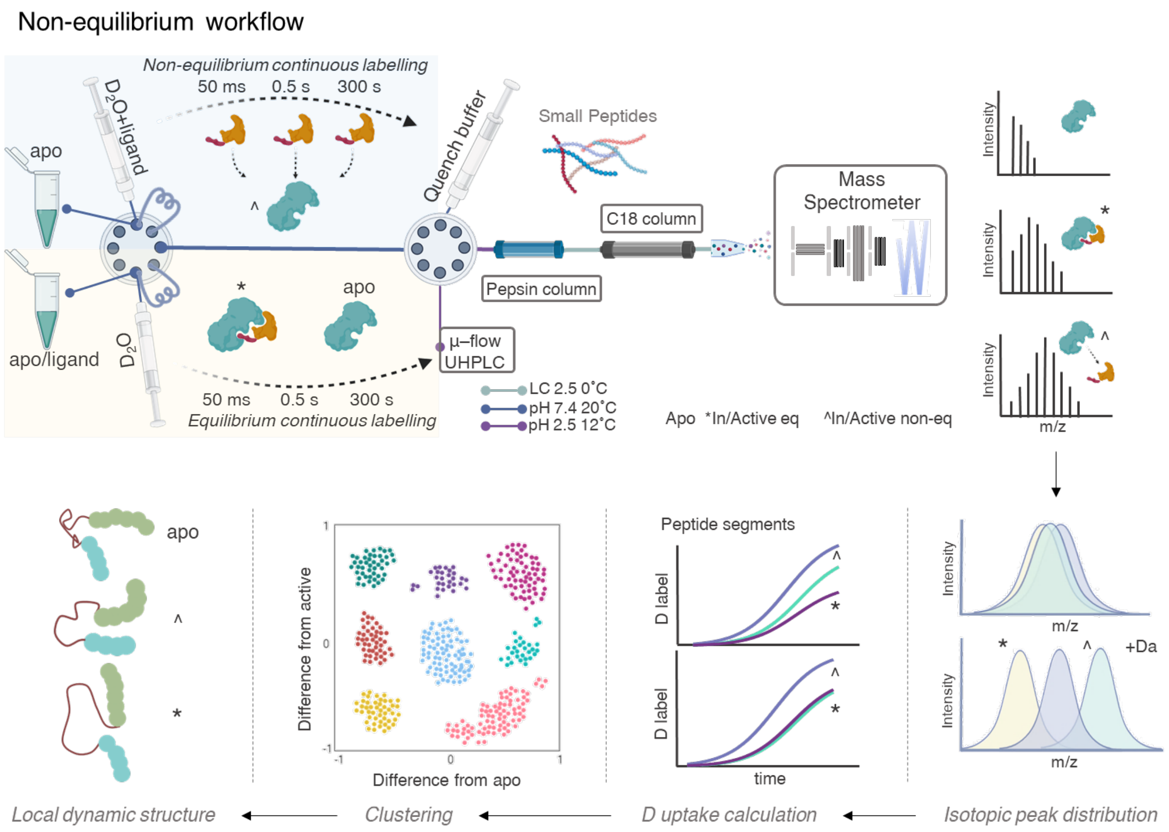


**Figure S1.** Scheme for the continuous non-equilibrium HDX-MS approach. The protein sample is labelled at 20°C at an appropriate pH via three conditions, including apo, apo/ligand at equilibrium and apo/ligand at non-equilibrium. The difference from conventional HDX experiments is the non-equilibrium continuous labelling, achieved by placing the ligand in the labelling buffer, and only allowing them to interact for the preferred time. Upon quenching, digestion, separation and detection, the non-equilibrium data is analysed by clustering the peptide segments by their deuterium uptake difference from active/inactive state. Each cluster is then correlated to specific behaviour upon allosteric activation/inhibition.

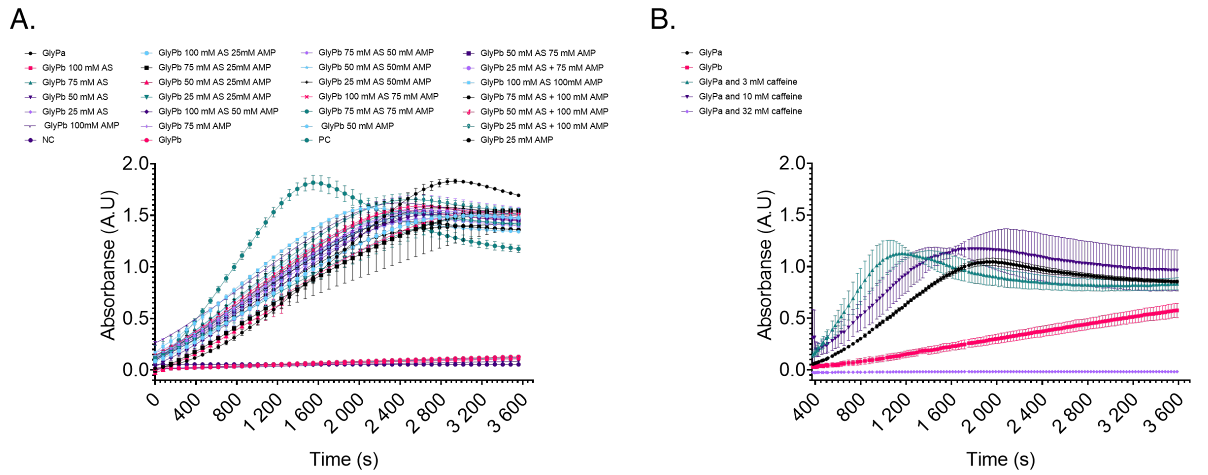


**Figure S2: (A)** Activation of Glycogen phosphorylase b by varying concentrations of AMP/AS. Full enzyme reaction curves for all samples, see legend. For comparability, all samples were tested on the same day with same conditions and reagents. Hence, only two replicates for each sample could be completed, limited by the size of the 96-well plate. Absorbance was measured at 450 nm for 60 minutes. **(B)** Inhibition of Glycogen phosphorylase a by varying concentrations of caffeine. Full enzyme reaction curves for all samples, see legend. All samples were tested on the same day with same conditions and reagents. Absorbance was measured at 450 nm for 60 minutes.

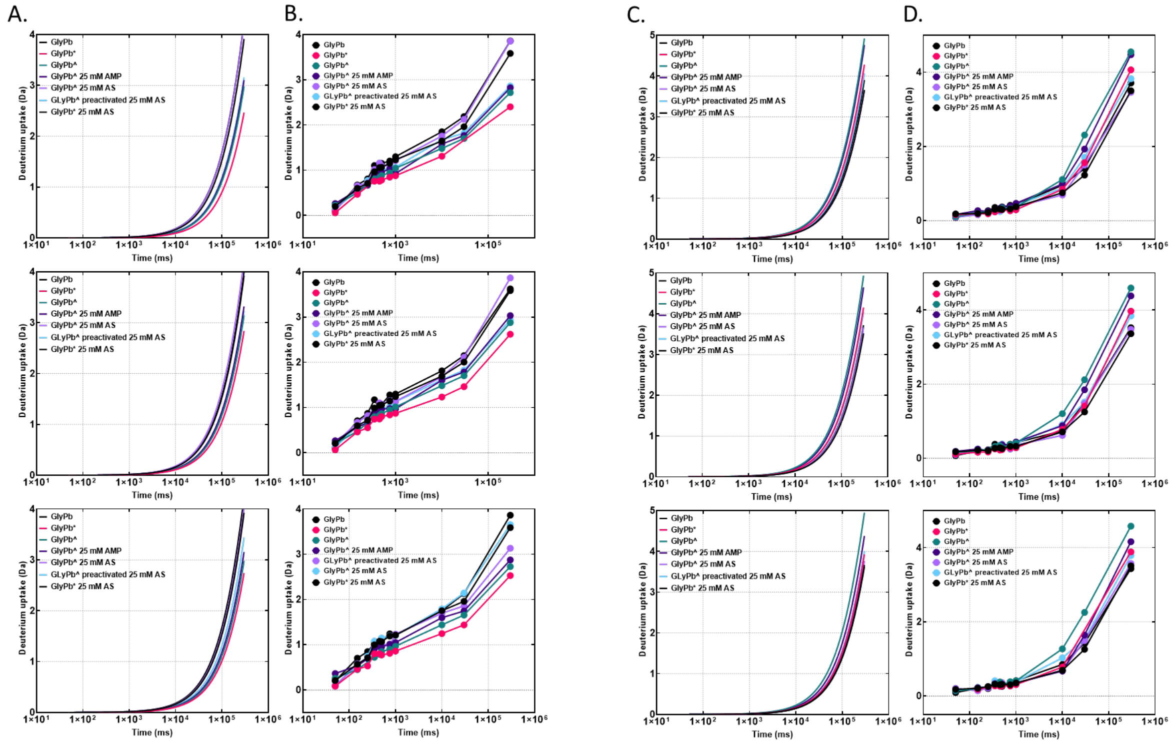


**Figure S3. Ammonium sulfate influence on the binding site residues 38-47. A.** Raw data fits of each state, including GlyPb, GlyPb* - activated with 25/25 mM AMP/AS at equilibrium, GlyPb^ - activated with 25/25 mM AMP/AS at non-equilibrium, GlyPb^ 25 mM AMP - activated with 25 mM AMP at non-equilibrium, GlyPb^ 25 mM AS - activated with 25 mM AS at non-equilibrium, GlyPb^ preactivated 25 mM AS - activated with 25 mM AMP at non-equilibrium, but preactivated with 25 mM AS at equilibrium, GlyPb* 25 mM AS - activated with 25 mM AS at equilibrium. The fitted $k_{obs}$ was used in further analysis (n=3). **B.** Raw uptake data of each state, per replicate, at 11 time points (50-300000 ms). **Ammonium sulfate influence on the tower helix residues 266-275. C.** Raw data fits of each state, including GlyPb, GlyPb* - activated with 25/25 mM AMP/AS at equilibrium, GlyPb^ - activated with 25/25 mM AMP/AS at non-equilibrium, GlyPb^ 25 mM AMP - activated with 25 mM AMP at non-equilibrium, GlyPb^ 25 mM AS - activated with 25 mM AS at non-equilibrium, GlyPb^ preactivated 25 mM AS - activated with 25 mM AMP at non-equilibrium, but preactivated with 25 mM AS at equilibrium, GlyPb* 25 mM AS - activated with 25 mM AS at equilibrium. The fitted $k_{obs}$ was used in further analysis (n=3). **D.** Raw uptake data of each state, per replicate, at 11 time points (50-300000 ms).

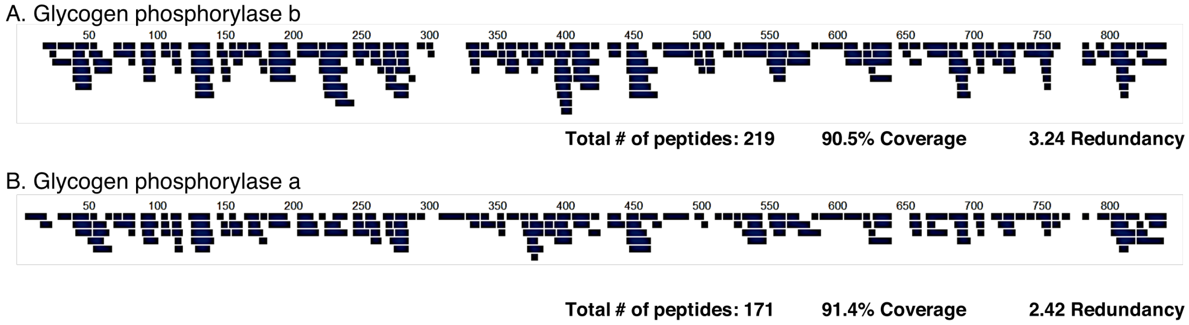


**Figure S4. Sequence coverage of glycogen phosphorylase from rabbit muscle.** The peptides obtained via peptic digestion and LC-IMS/MS analysis are shown as black bars along the sequence numbers. Each bar under the sequence number annotation indicates an identified peptic peptide by eight MSE fragmentation experiments with different collision energy ramps, that was monitored during all HDX-MS experiments. **A.** These 219 peptides of glycogen phosphorylase b cover up to 800 amino acids of the total amino acid residues in the proteins, yielding a linear sequence coverage of up to 90.5%, with 3.24 redundancy. **B.** Similarly, 171 peptides of glycogen phosphorylase a, covering 91.4% of the sequence with 2.42% redundancy were identified and monitored during the HDX-MS experiments.

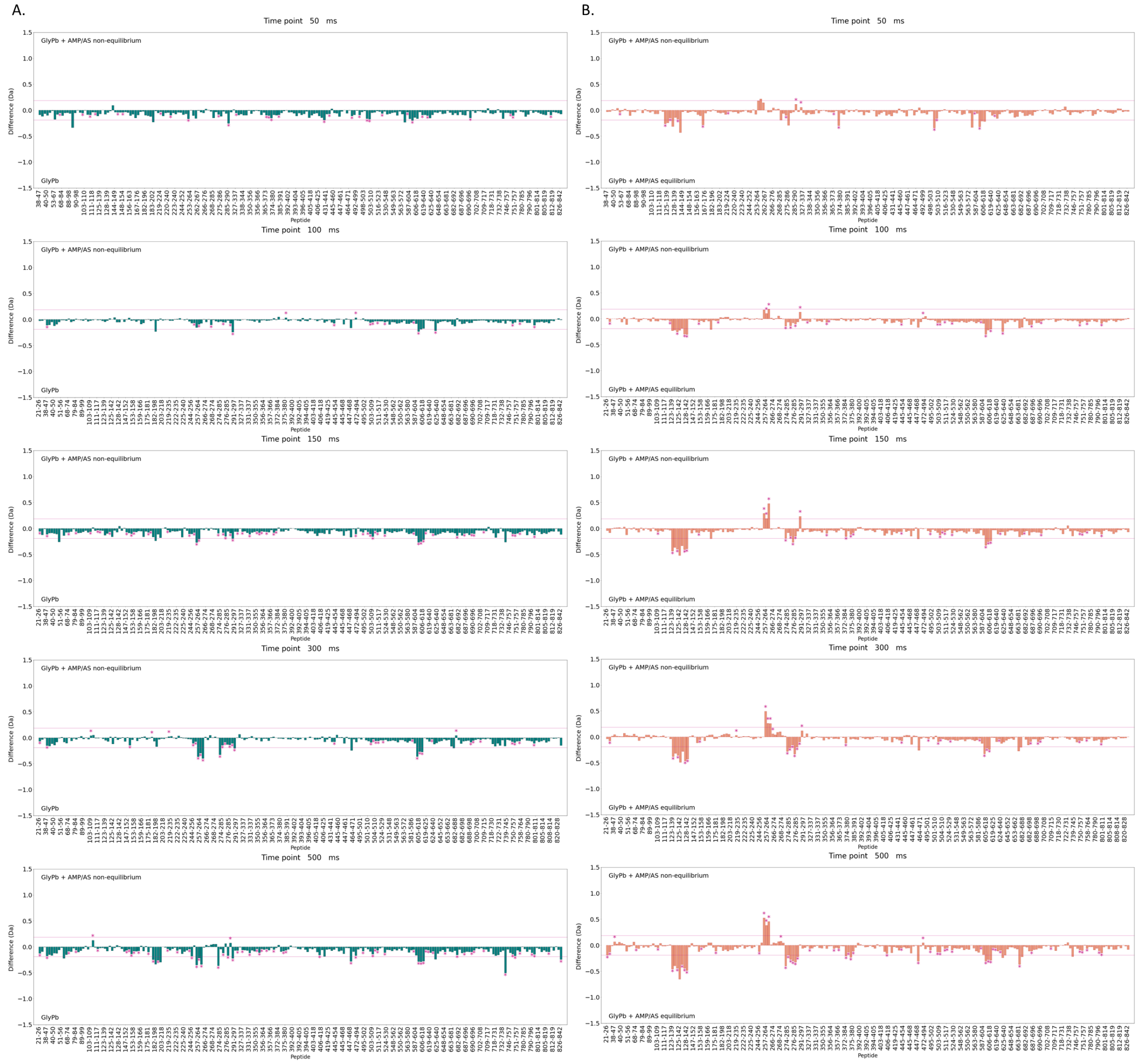


**Figure S5. State D uptake difference maps for fast mixing times (50, 100, 150, 300 and 500 ms).** Difference of the sum of observed deuterium uptake at 50, 100, 150, 300 and 500 ms between three states including GlyPb, GlyPb activated at equilibrium with 25/25 mM AMP/AS, and GlyPb activated at non-equilibrium with 25/25 mM AMP/AS. The deuterium uptake data from the state denoted on the bottom of each plot was subtracted from the data of the state denoted on the top in order to create the deuterium uptake difference plot. Relative protection leads to a more negative value (bar on bottom side); deprotection (e.g., from an exposed domain interface) results in a more positive value (bar on upper side). Each vertical bar represents a single peptide. Horizontal scale is peptides residues, from start residue (left) to end (right). The pink horizontal lines denote the significant difference between the states derived from the global significance threshold, explained elsewhere.[44,51] Pink asterisks denote the significant difference in the D-label per peptide and time point derived from a T-test.

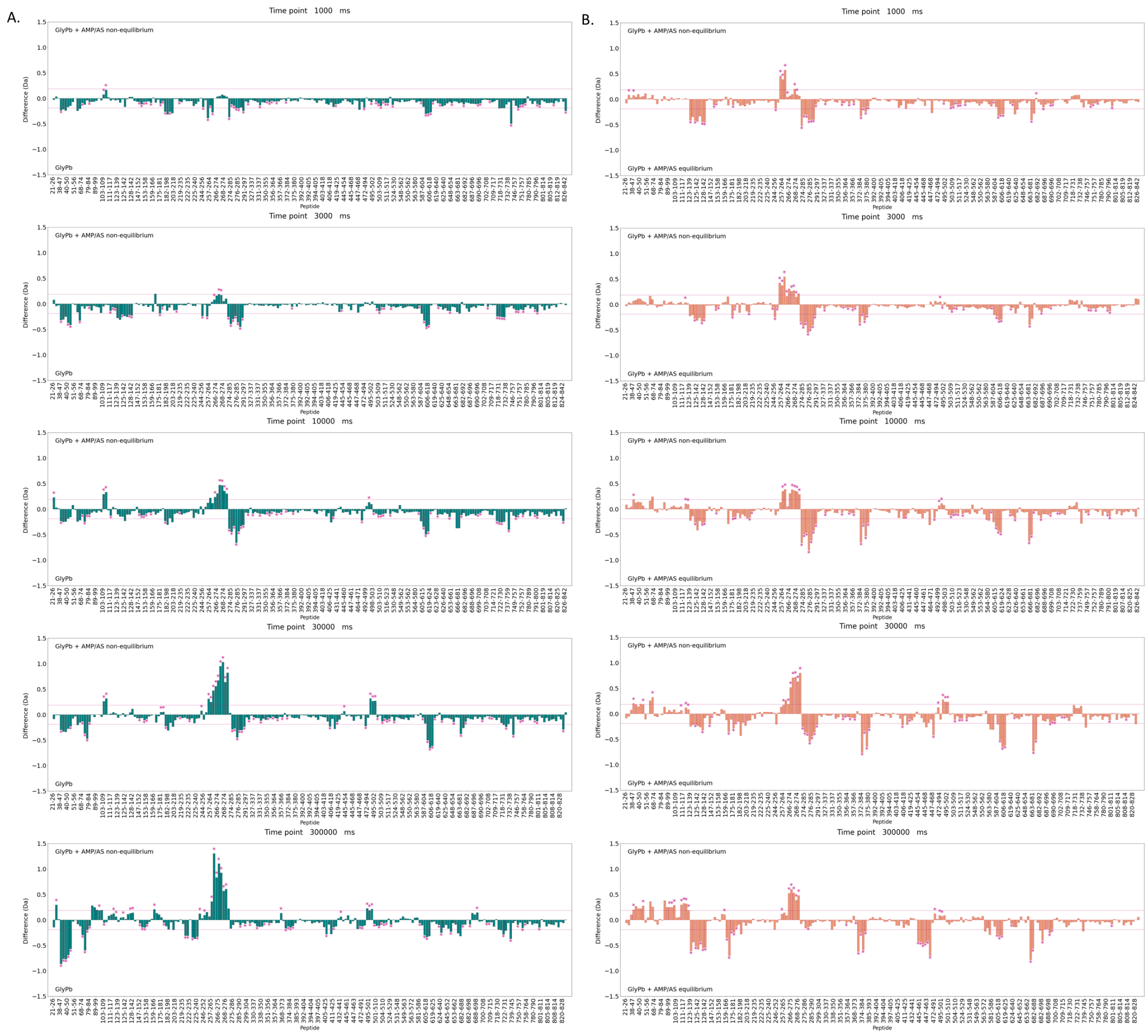


**Figure S6. State D uptake difference maps for slow mixing times (1, 3, 10, 30 and 300 s).** Difference of the sum of observed deuterium uptake at 1,3,10, 30 and 300 s between three states including GlyPb, GlyPb activated at equilibrium with 25/25 mM AMP/AS, and GlyPb activated at non-equilibrium with 25/25 mM AMP/AS. The deuterium uptake data from the state denoted on the bottom of each plot was subtracted from the data of the state denoted on the top in order to create the deuterium uptake difference plot. Relative protection leads to a more negative value (bar on bottom side); deprotection (e.g., from an exposed domain interface) results in a more positive value (bar on upper side). Each vertical bar represents a single peptide. Horizontal scale is peptides residues, from start residue (left) to end (right). The pink horizontal lines denote the significant difference between the states derived from the global significance threshold, explained elsewhere.[44,51] Pink asterisks denote the significant difference in the D-label per peptide and time point derived from a T-test.

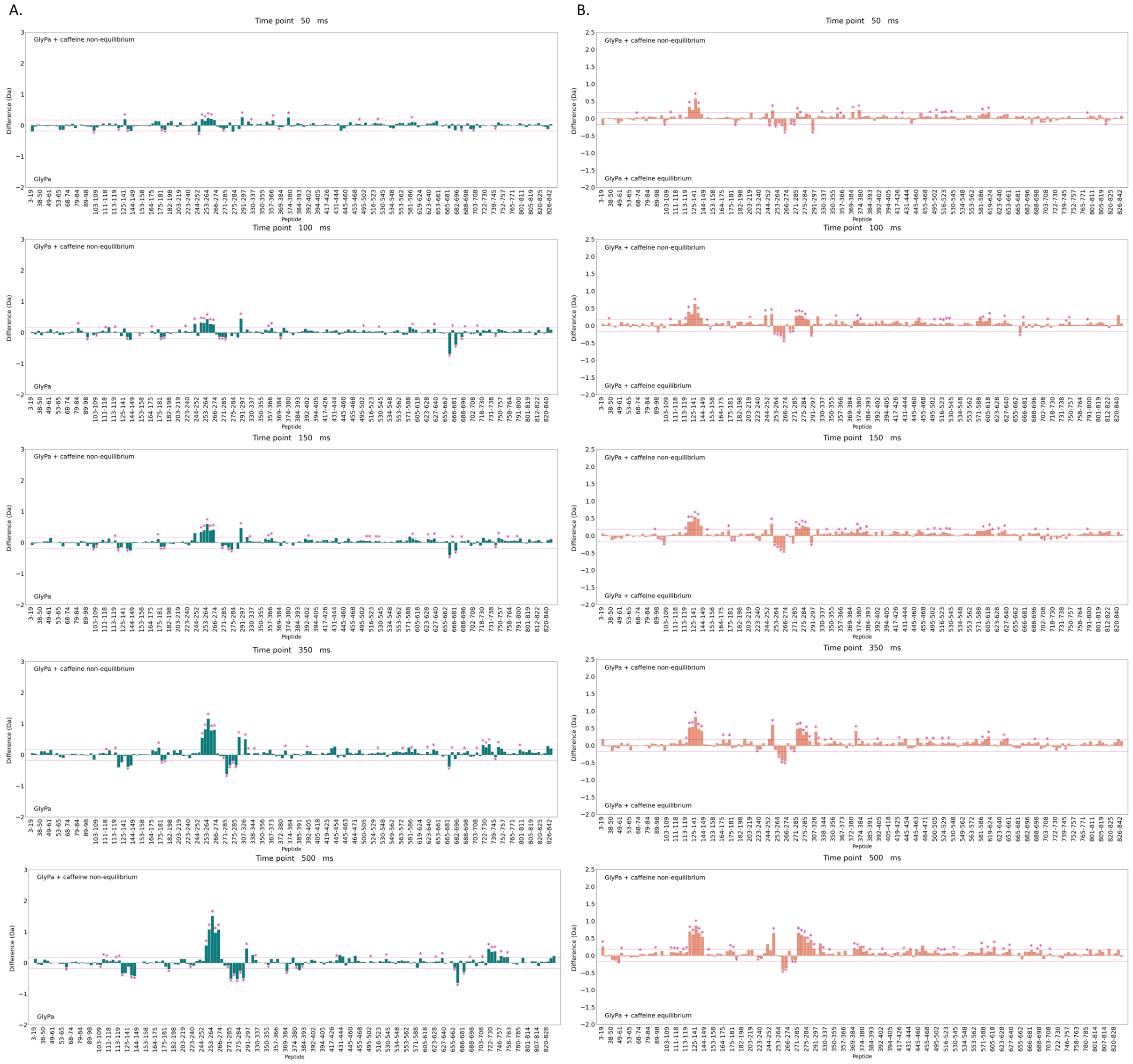


**Figure S7. State D uptake difference maps for fast mixing times (50, 100, 150, 300 and 500 ms).** Difference of the sum of observed deuterium uptake at 50, 100, 150, 300 and 500 ms between three states including GlyPa, GlyPb inhibitd at equilibrium with 32 mM caffeine, and GlyPb inhibitd at non-equilibrium with 32 mM caffeine. The deuterium uptake data from the state denoted on the bottom of each plot was subtracted from the data of the state denoted on the top in order to create the deuterium uptake difference plot. Relative protection leads to a more negative value (bar on bottom side); deprotection (e.g., from an exposed domain interface) results in a more positive value (bar on upper side). Each vertical bar represents a single peptide. Horizontal scale is peptides residues, from start residue (left) to end (right). The pink horizontal lines denote the significant difference between the states derived from the global significance threshold, explained elsewhere.[44,51] Pink asterisks denote the significant difference in the D-label per peptide and time point derived from a T-test.

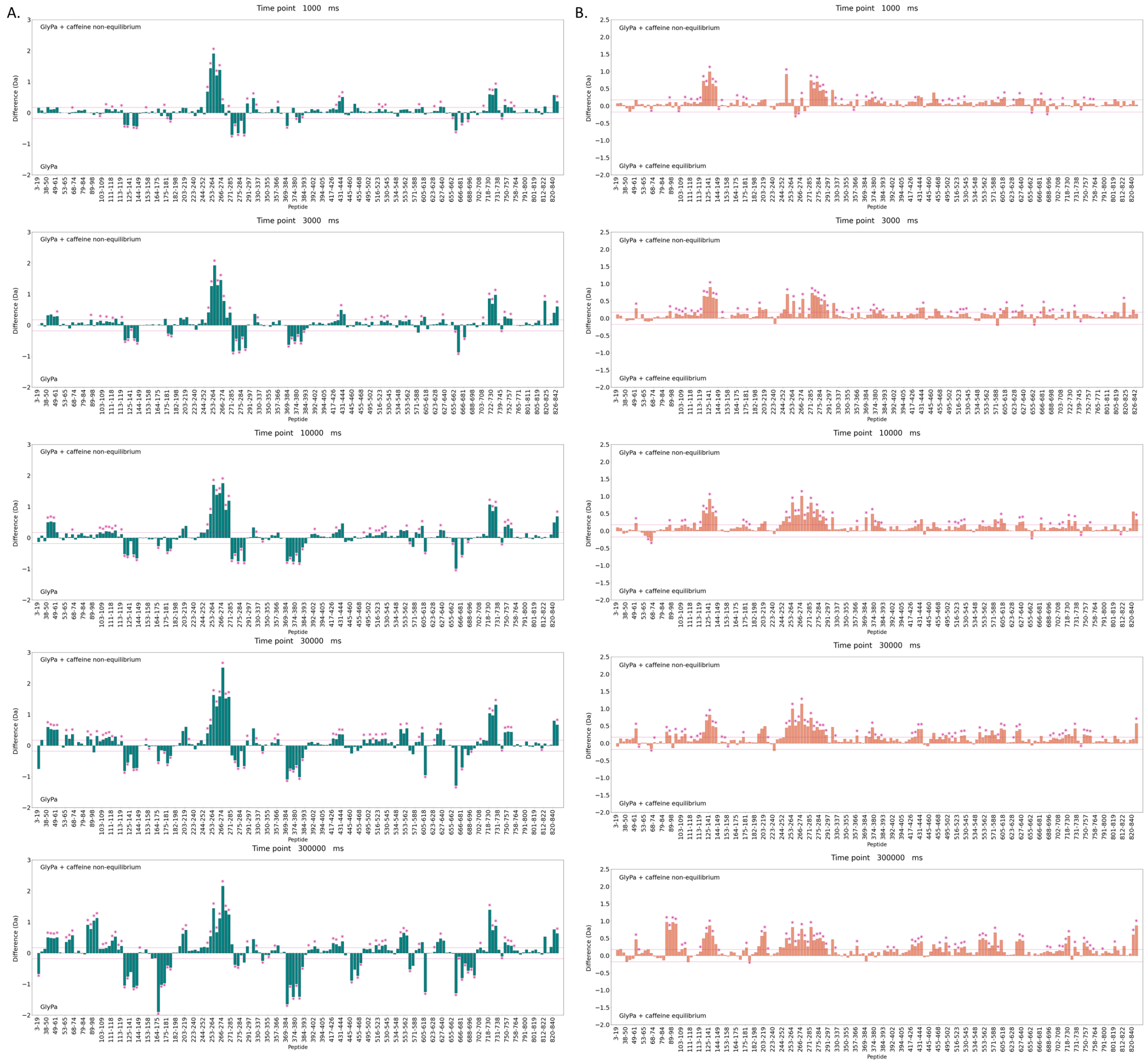


**Figure S8. State D uptake difference maps for fast mixing times (1, 3, 10, 30 and 300 s).** Difference of the sum of observed deuterium uptake at 1, 3, 10, 30 and 300 s between three states including GlyPa, GlyPb inhibitd at equilibrium with 32 mM caffeine, and GlyPb inhibitd at non-equilibrium with 32 mM caffeine. The deuterium uptake data from the state denoted on the bottom of each plot was subtracted from the data of the state denoted on the top in order to create the deuterium uptake difference plot. Relative protection leads to a more negative value (bar on bottom side); deprotection (e.g., from an exposed domain interface) results in a more positive value (bar on upper side). Each vertical bar represents a single peptide. Horizontal scale is peptides residues, from start residue (left) to end (right). The pink horizontal lines denote the significant difference between the states derived from the global significance threshold, explained elsewhere.[44,51] Pink asterisks denote the significant difference in the D-label per peptide and time point derived from a T-test.

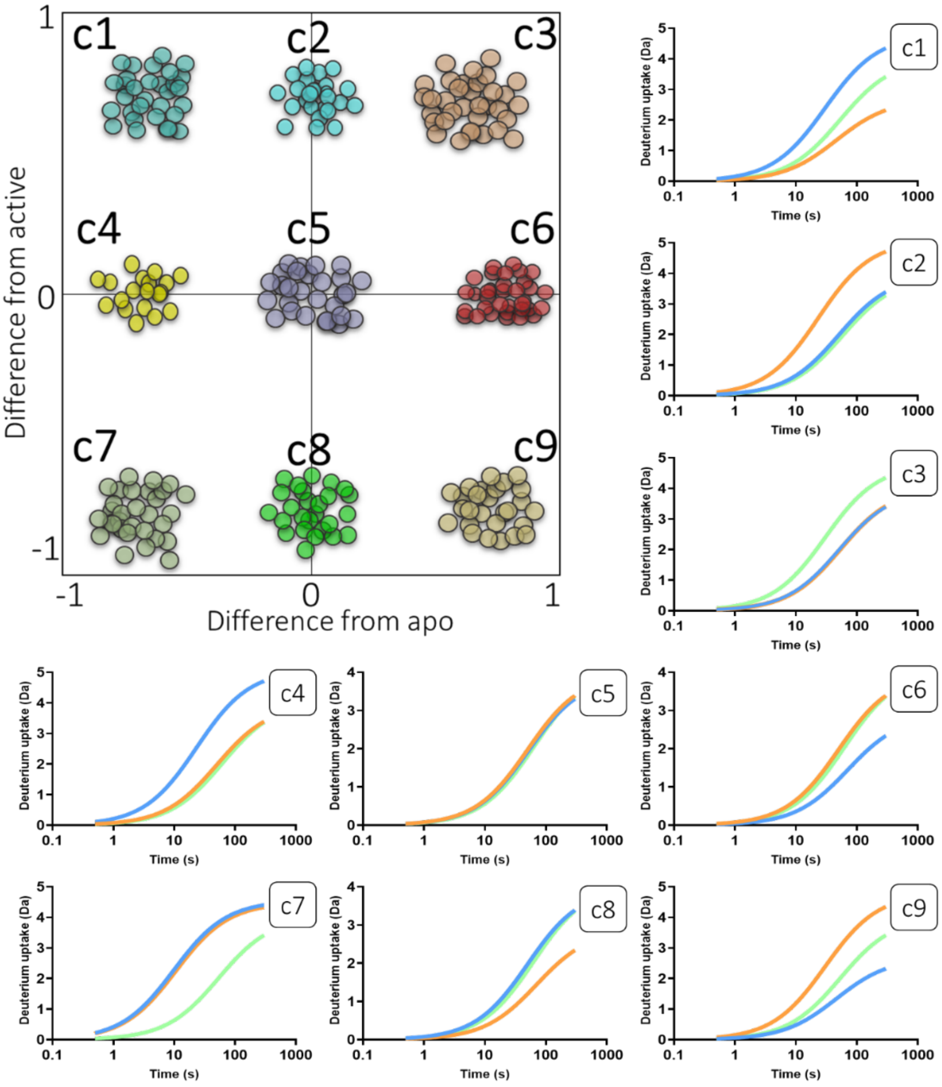


**Figure S9. Theoretical behaviors of peptide segments upon non-equilibrium activation.** Clustering by k means per peptide of the sum difference of the uptake data for the specific states after normalization involving non-equilibrium minus apo (x-axis), and non-equilibrium minus equilibrium (y-axis). Approximately, 9 clusters are anticipated, each cluster representing the location of the non-equilibrium kinetic uptake plot in respect to the apo and equilibrium uptake plots. From top to bottom the clusters include: c1 (x=-1, y=1), c2 (x=0, y=1), c3 (x=1, y=1), c4 (x=-1, y=0), c5 (x=0, y=0), c6 (x=1, y=0), c7 (x=-1, y=-1), c8 (x=0, y=-1), c9 (x=1, y=-1). Uptake kinetic plots are also represented for each cluster, where blue denotes apo state, orange ligand-bound at equilibrium and green the non-equilibrium state.

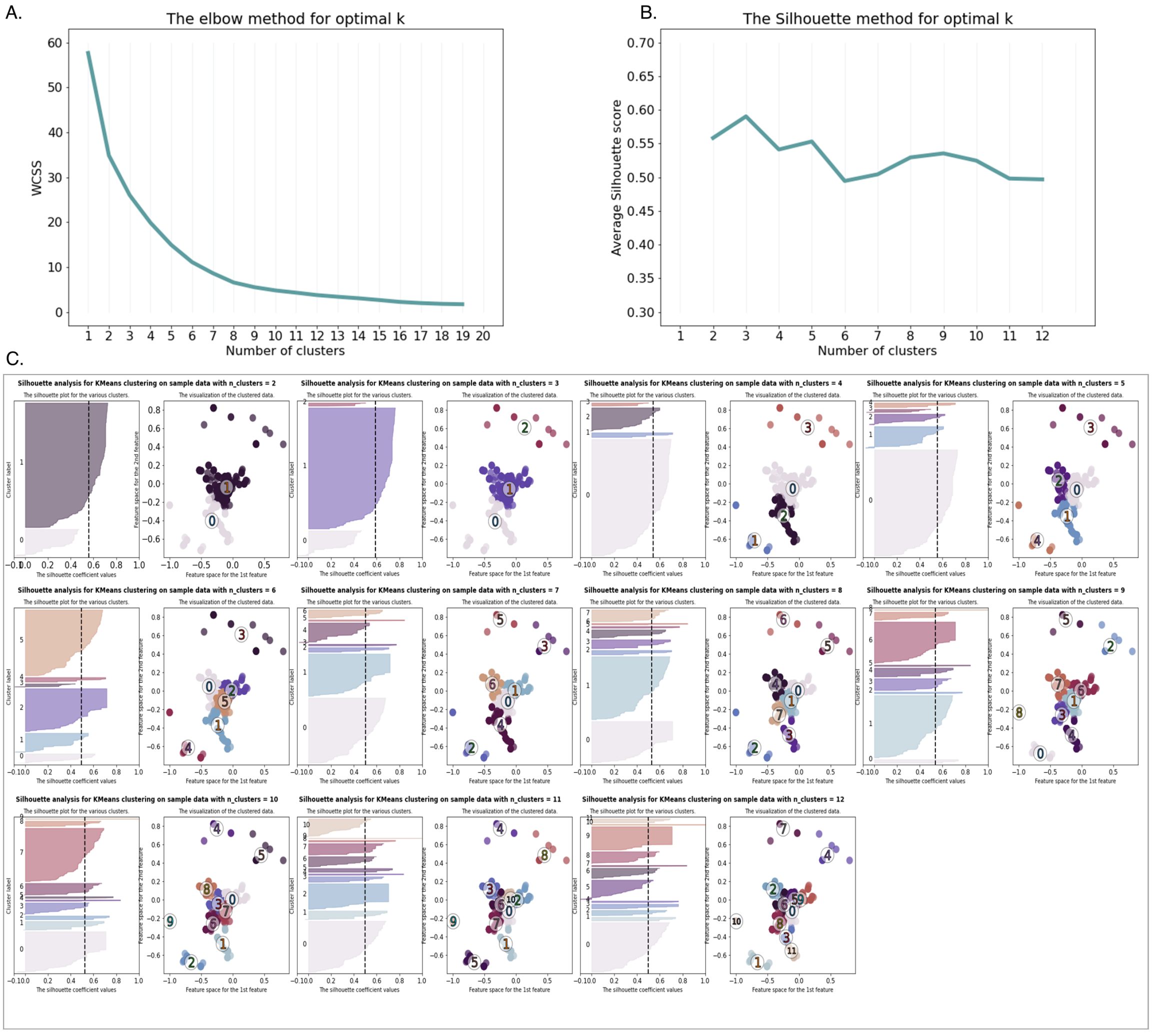


**Figure S10. Assessing the quality of k-means clusters fit on the data of GlyPb. A.** Elbow method for optimal value of clusters. **B.** Silhouette score analysis for clustering quality. The Silhouette score was calculated for the whole data set for 2-7 clusters. **C.** Silhouette analysis for 2-12 clusters.

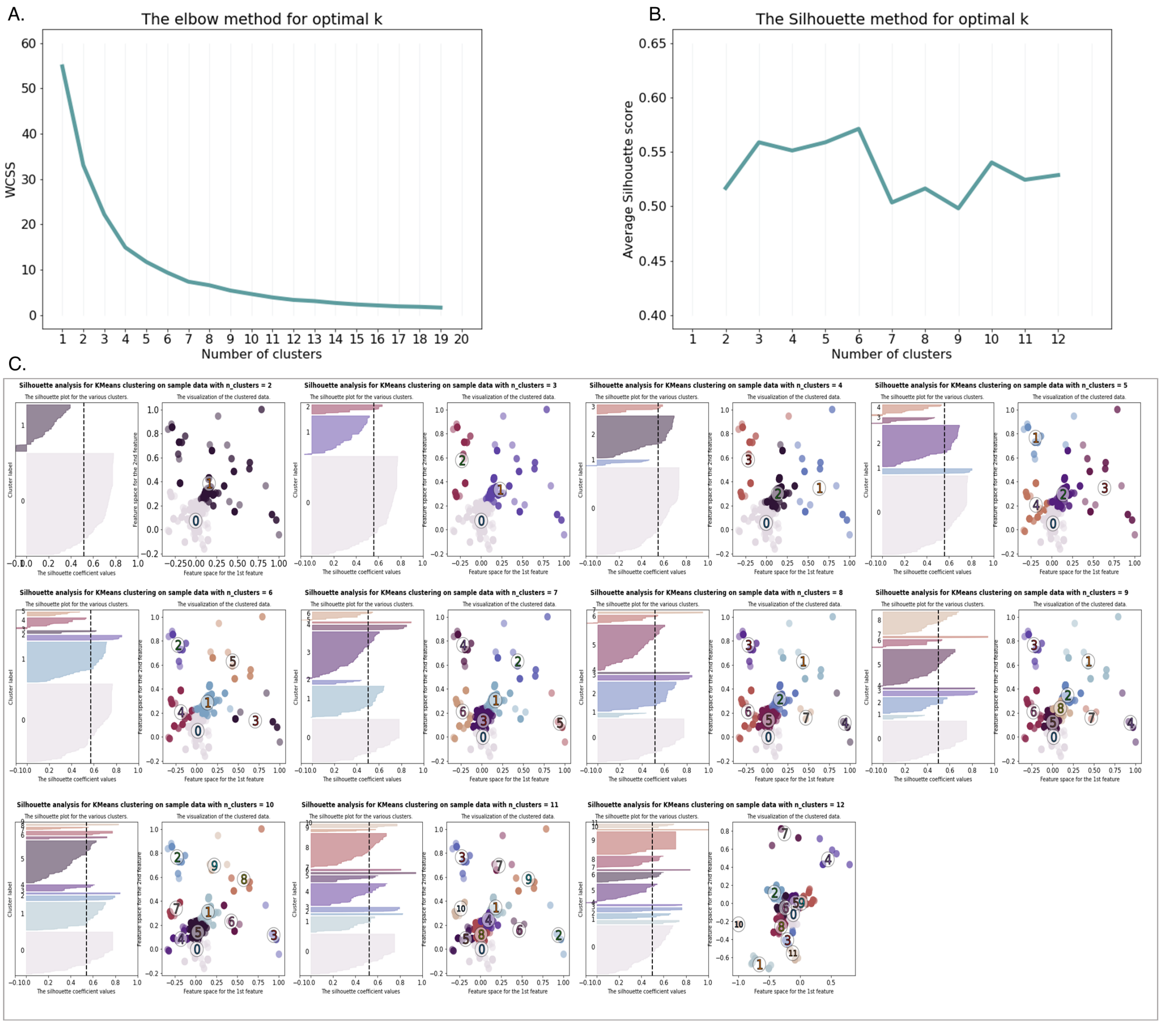


**Figure S11. Assessing the quality of k-means clusters fit on the data of GlyPa. A.** Elbow method for optimal value of clusters. **B.** Silhouette score analysis for clustering quality. The Silhouette score was calculated for the whole data set for 2-7 clusters. **C.** Silhouette analysis for 2-12 clusters.

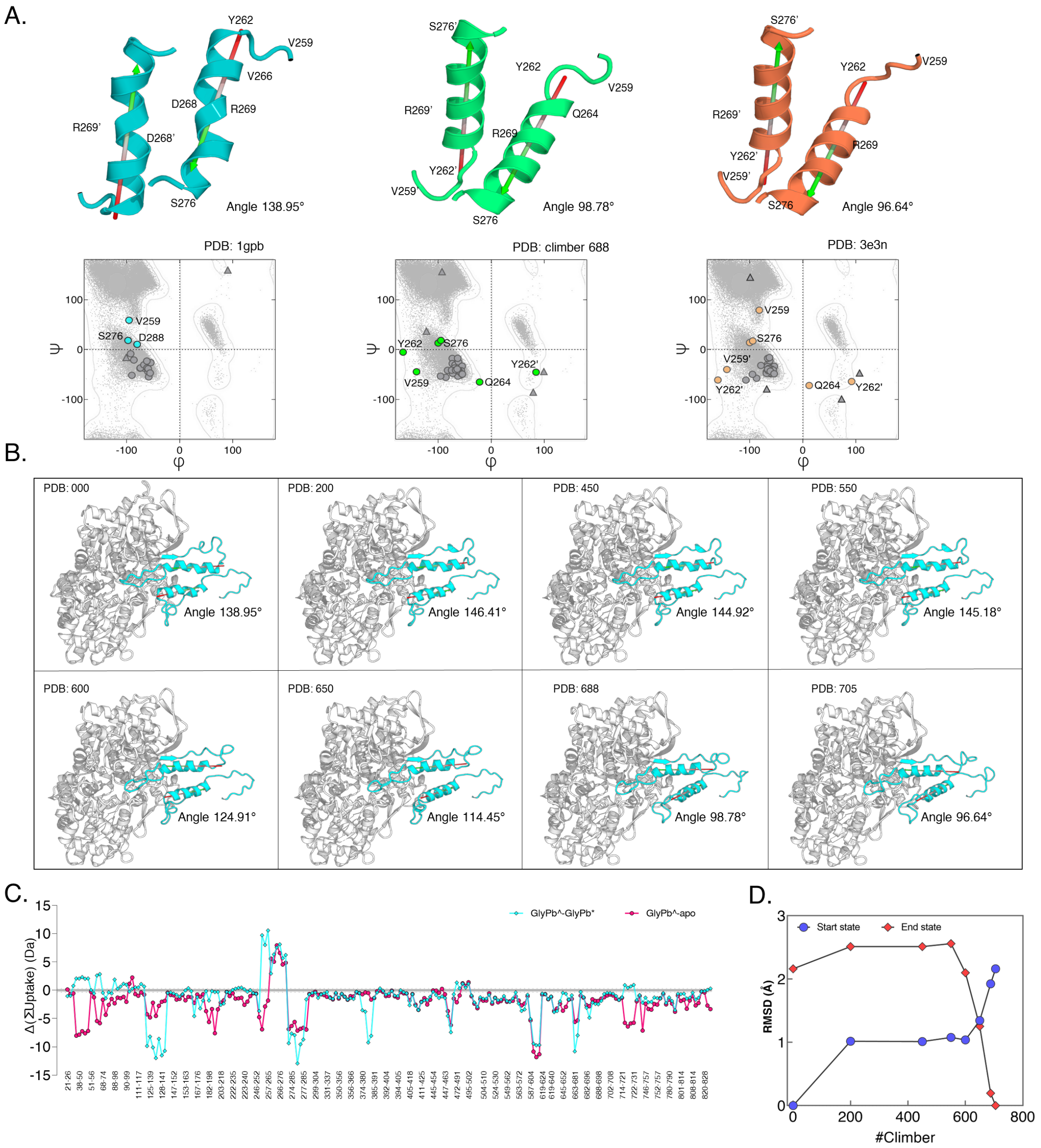


**Figure S12. Rotation and transition of the tower helix upon AMP activation of GlyPb. A.** Representation of the tower helix dynamics (residues 259-277), from T-state (blue), climber transition (green) and R-state (orange), with calculated angles between the α7 and α7' helices. Ramachandran plots are also included, calculated from the corresponding pdb files, with specific helix outliers highlighted. **B.** PDBs from the climber transition of GlyPb activation with AMP, including start pdb 000, 200, 450, 550, 600, 650, 688 and end state. The tower helices are colored cyan with the connecting 250 and 280s loops from both dimers. **C.** Difference plot representing the difference of the sum of uptake between the non-equilibrium GlyPb + AMP/AS data and the equilibrium GlyPb + AMP/AS data (pink), and the non-equilibrium GlyPb + AMP/AS data and GlyPb (cyan). **D.** The RMSD difference between the start (blue) and end (red) state with the 200, 450, 550, 600, 650, 688 PDBs.

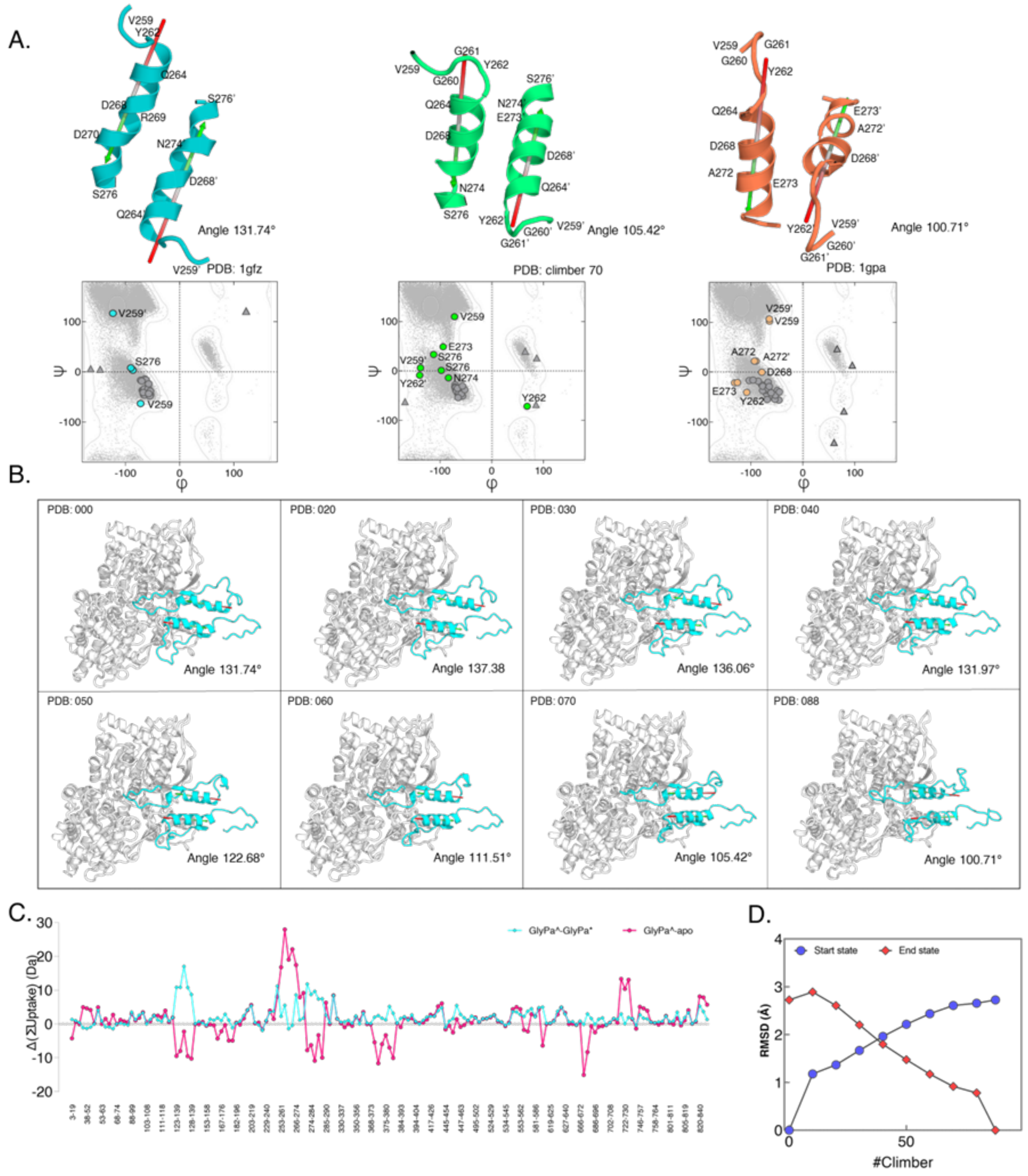


**Figure S13. Rotation and transition of the tower helix upon caffeine inhibition of GlyPa. A.** Representation of the tower helix dynamics (residues 259-277), from T-state (blue), climber transition (green) and R-state (orange), with calculated angles between the α7 and α7' helices. Ramachandran plots are also included, calculated from the corresponding pdb files, with specific helix outliers highlighted. **B.** PDBs from the climber transition of GlyPa inhibition with caffeine, including start pdb 000, 020, 030, 040, 050, 060, 070, and end state. The tower helices are colored cyan with the connecting 250 and 280s loops from both dimers. **C.** Difference plot representing the difference of the sum of uptake between the non-equilibrium GlyPa + caffeine data and the equilibrium GlyPa + caffeine data (pink), and the non-equilibrium GlyPa + caffeine data and GlyPa (cyan). **D.** The RMSD difference between the start (blue) and end (red) state with the 020, 030, 040, 050, 060, 070 PDBs.

A.

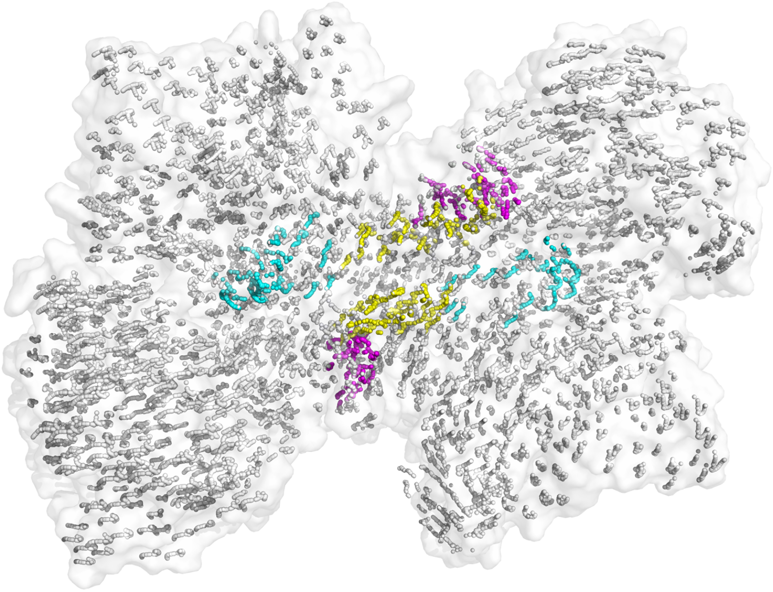

B.

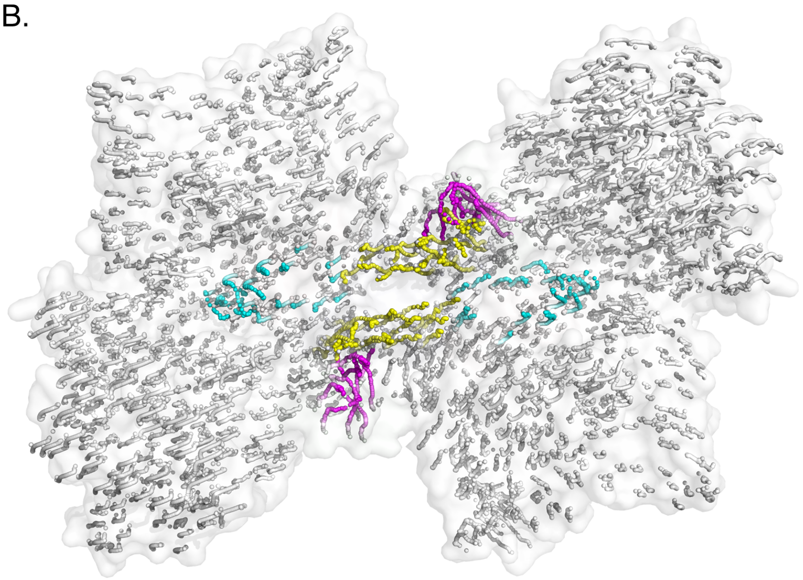

**Figure S14. Representation of the full transition upon activation/inhibition.** Each backbone nitrogen is represented as a sphere. Colors correspond to specific sections including the tower helix (yellow), 250 loop (pink) and 280 loop (blue). **A.** The movement of the backbone nitrogens upon AMP/AS activation. **B.** The movement of the backbone nitrogens upon caffeine inhibition.

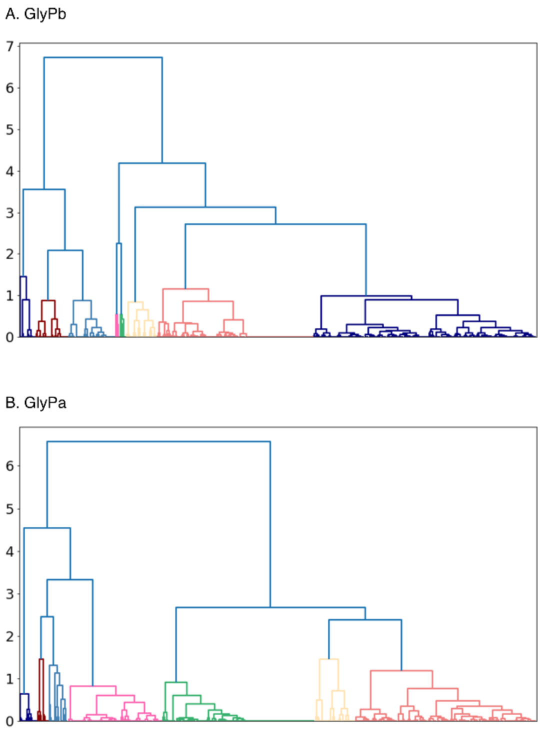


**Figure S15. Full dendrogram of hierarchical clustering.** The dendrogram was generated with ScyPy for all identified peptides monitored by non-equilibrium HDX, where above p=7 in truncate mode, 7 clusters of behaviors were observed. The dendrogram illustrates how each cluster is composed by drawing a U-shaped link between a non-singleton cluster and its children. The top of the U-link indicates a cluster merge. The two legs of the U-link indicate which clusters were merged. The length of the two legs of the U-link represents the distance between the child clusters. It is also the cophenetic distance between original observations in the two children clusters.

**Figure S16.** HDX uptake curves of GlyPb per peptide. Absolute data for three protein states shown (blue – apo GlyPb; orange – AMP-AS bound activated GlyPb; green – AMP-AS bound non-equilibrium activated GlyPb). Peptide sequence and amino acid number given above the plot. Y-axis shows absolute % deuteration at labeling time (ms) on X-axis.

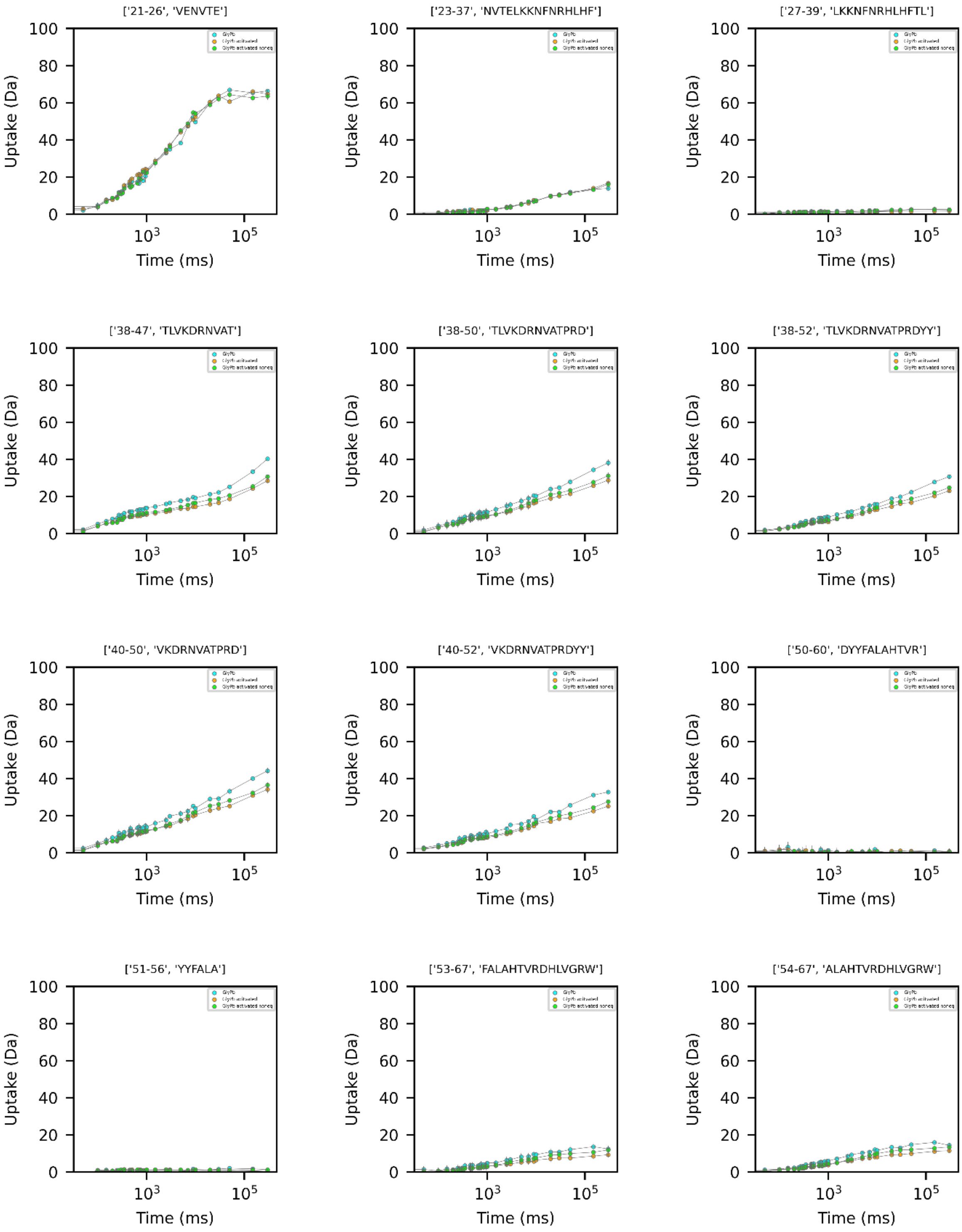

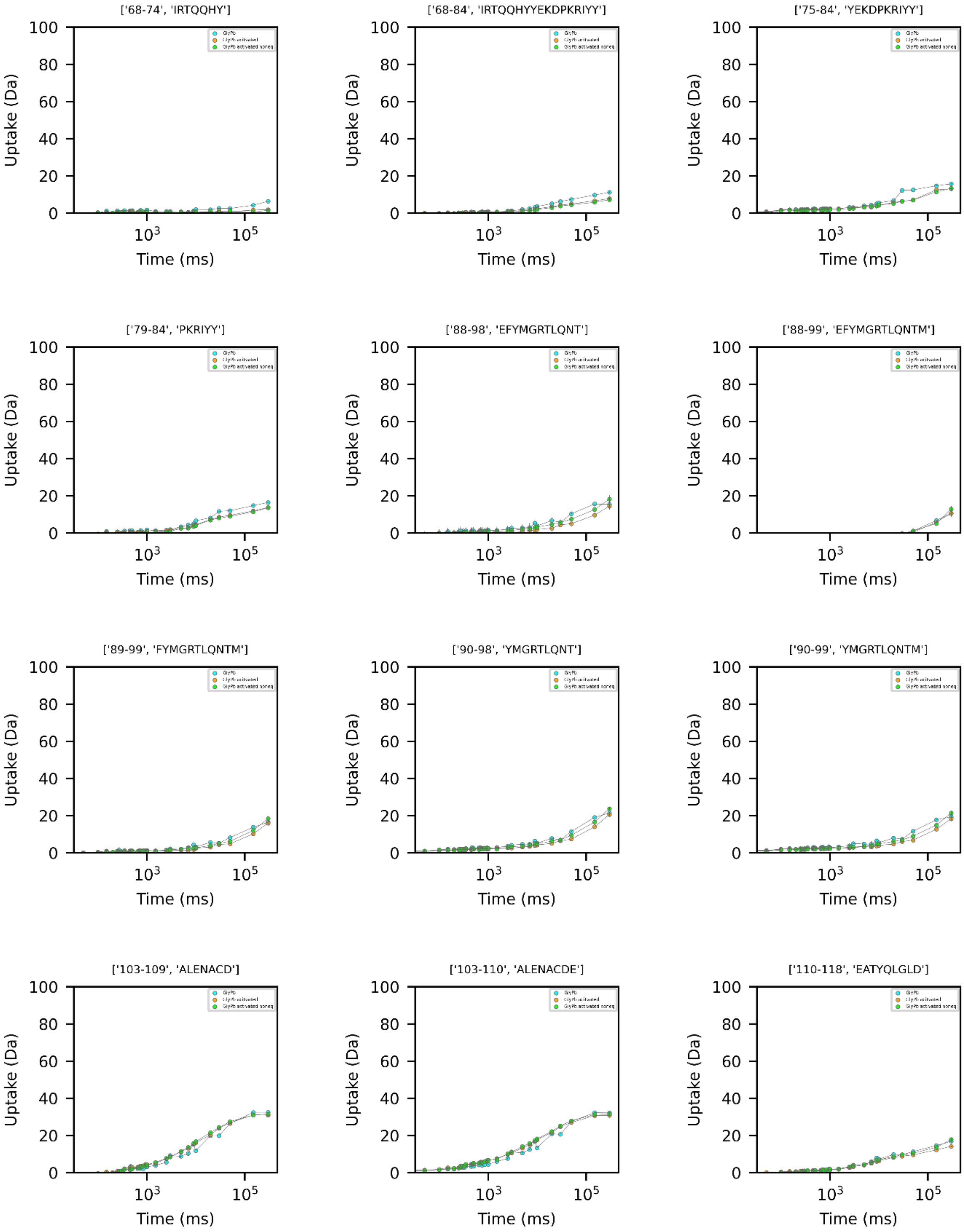
['68-74', 'IRTQQHY']
['68-84', 'IRTQQHYYEKDPKRIYY']
['75-84', 'YEKDPKRIYY']
['79-84', 'PKRIYY']
['88-98', 'EFYMGRTLQNT']
['88-99', 'EFYMGRTLQNTM']
['89-99', 'FYMGRTLQNTM']
['90-98', 'YMGRTLQNT']
['90-99', 'YMGRTLQNTM']
['103-109', 'ALENACD']
['103-110', 'ALENACDE']
['110-118', 'EATYQLGLD']
Uptake (Da)
Time (ms)
0
20
40
60
80
100
$10^3$
$10^5$

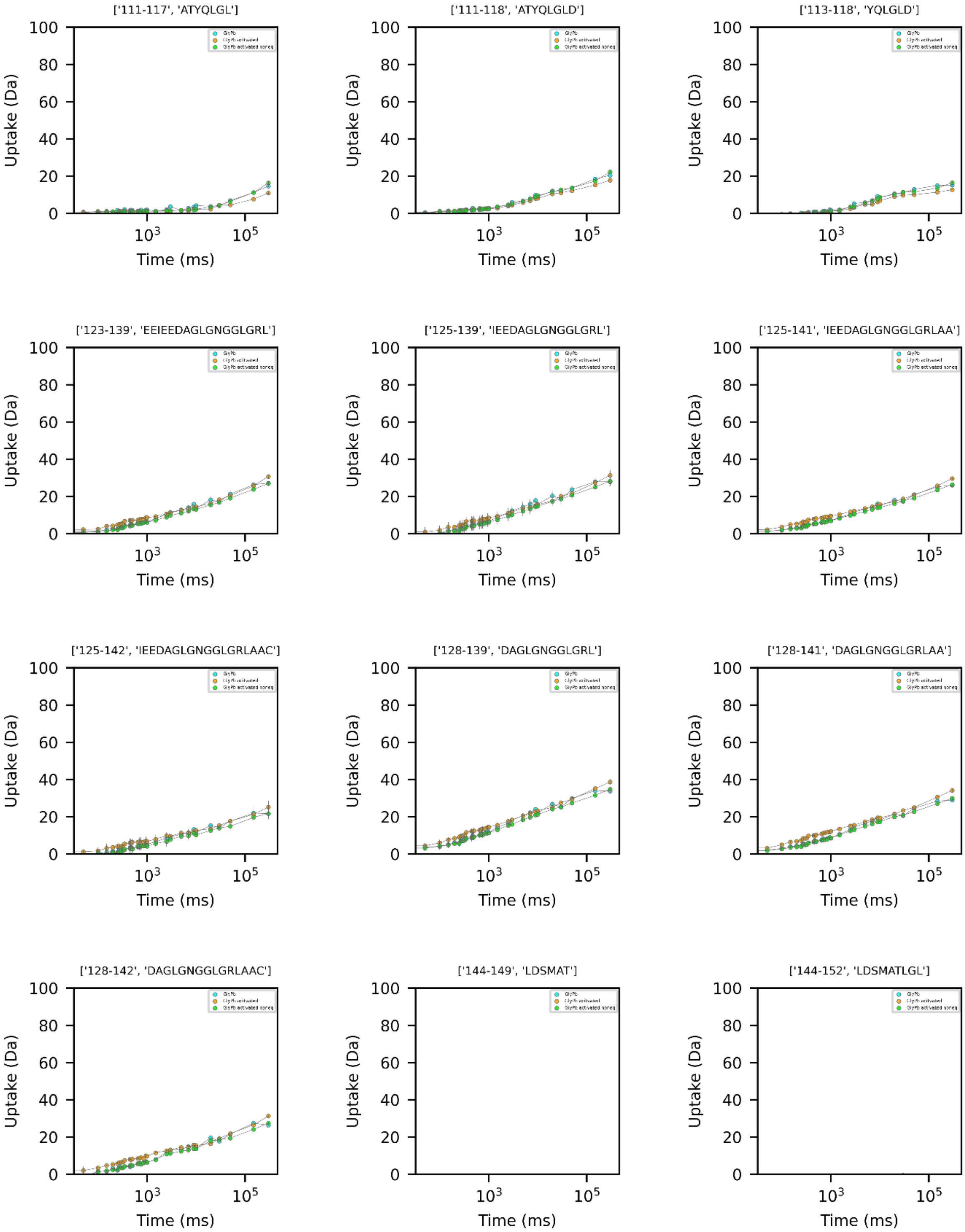
['111-117', 'ATYQLGL']
['111-118', 'ATYQLGLD']
['113-118', 'YQLGLD']
['123-139', 'EEIEEDAGLGNGGLGRL']
['125-139', 'IEEDAGLGNGGLGRL']
['125-141', 'IEEDAGLGNGGLGRLAA']
['125-142', 'IEEDAGLGNGGLGRLAAC']
['128-139', 'DAGLGNGGLGRL']
['128-141', 'DAGLGNGGLGRLAA']
['128-142', 'DAGLGNGGLGRLAAC']
['144-149', 'LDSMAT']
['144-152', 'LDSMATLGL']
Uptake (Da)
Time (ms)

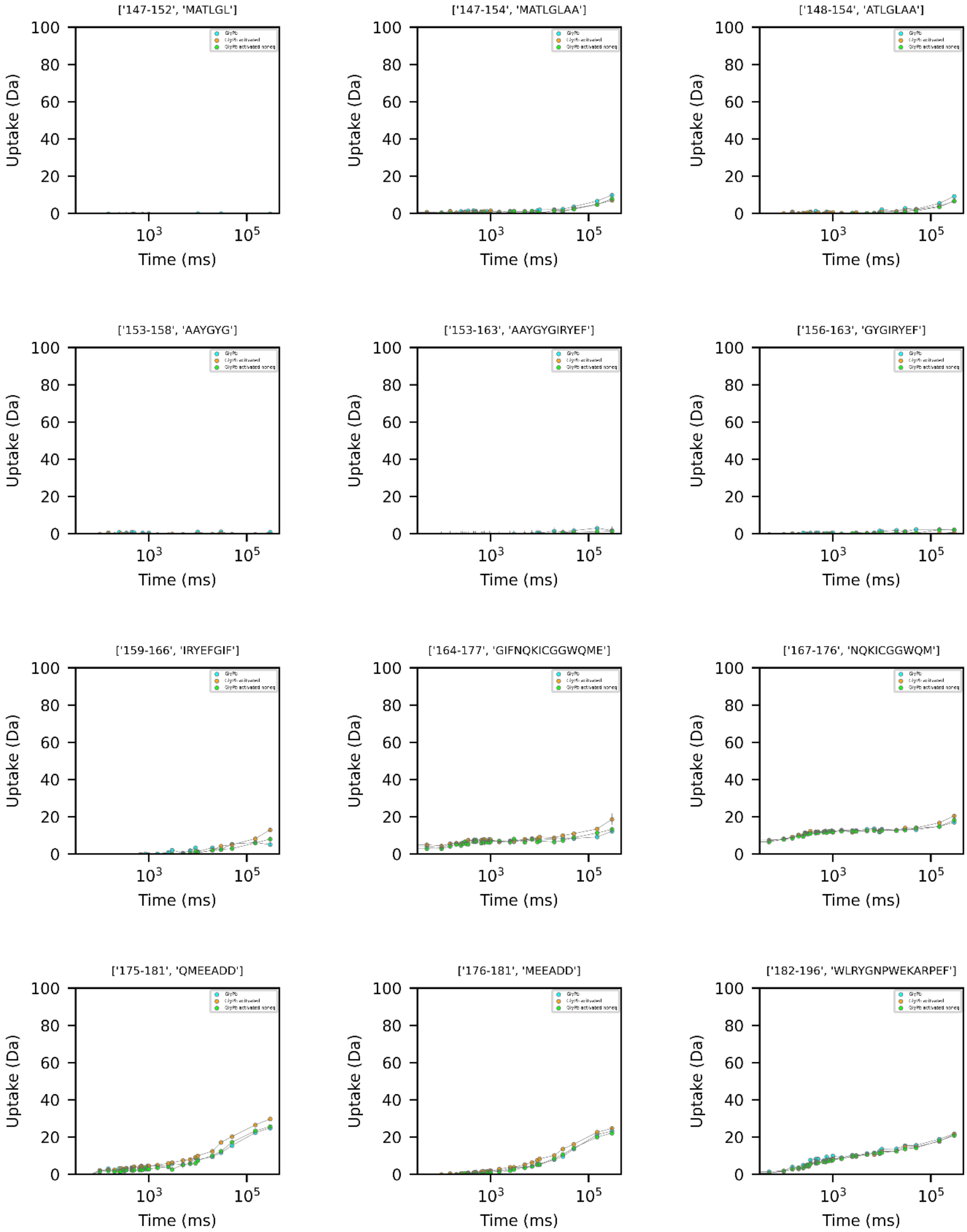
['147-152', 'MATLGL']
['147-154', 'MATLGLAA']
['148-154', 'ATLGLAA']
['153-158', 'AAYGYG']
['153-163', 'AAYGYGIRYEF']
['156-163', 'GYGIRYEF']
['159-166', 'IRYEFGIF']
['164-177', 'GIFNQKICGGWQME']
['167-176', 'NQKICGGWQM']
['175-181', 'QMEEADD']
['176-181', 'MEEADD']
['182-196', 'WLRYGNPWEKARPEF']
Uptake (Da)
Time (ms)
0
20
40
60
80
100
$10^3$
$10^5$

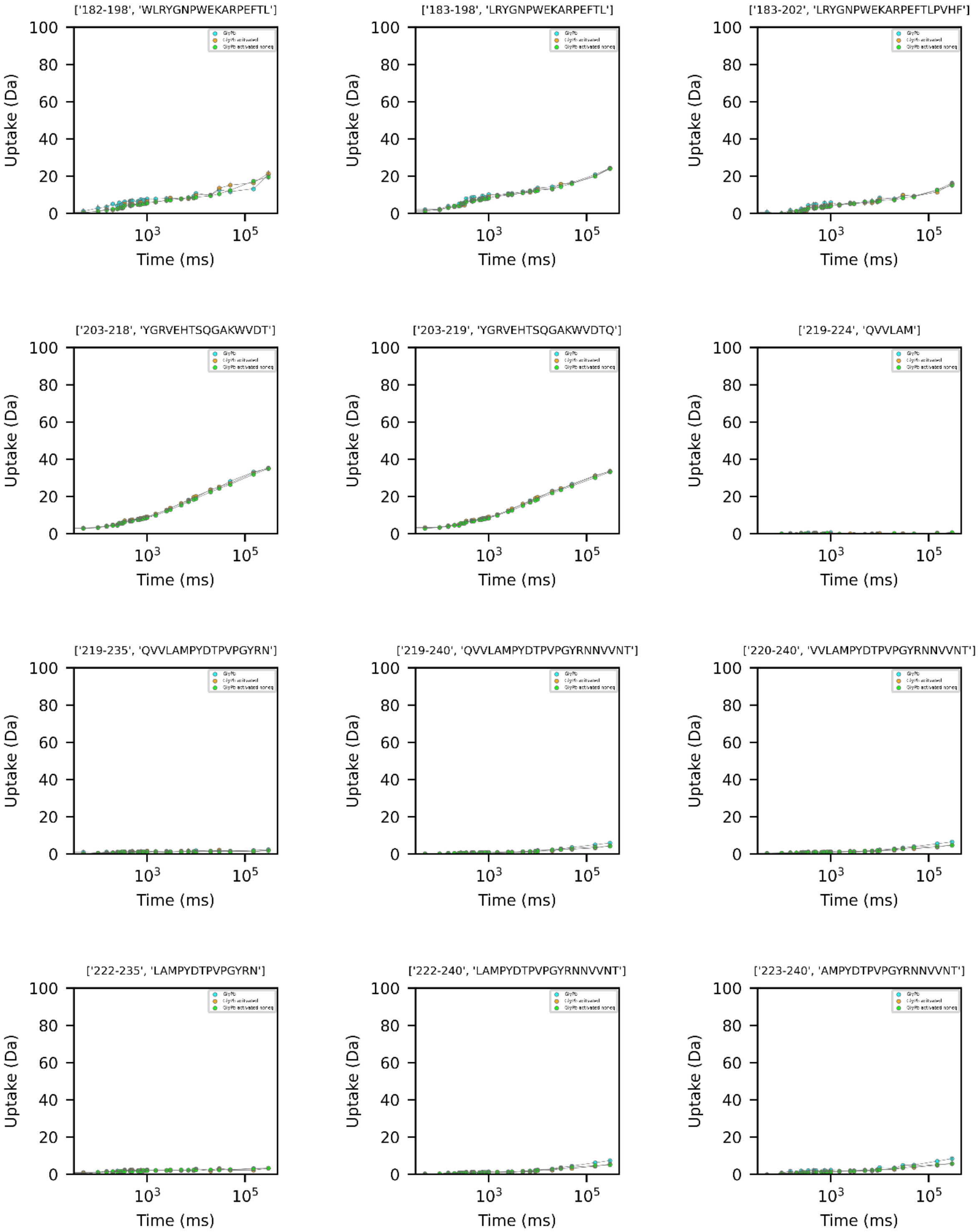
['182-198', 'WLRYGNPWEKARPEFTL']
['183-198', 'LRYGNPWEKARPEFTL']
['183-202', 'LRYGNPWEKARPEFTLPVHF']
['203-218', 'YGRVEHTSQGAKWVDT']
['203-219', 'YGRVEHTSQGAKWVDTQ']
['219-224', 'QVVLAM']
['219-235', 'QVVLAMPYDTPVPGYRN']
['219-240', 'QVVLAMPYDTPVPGYRNNVVNT']
['220-240', 'VVLAMPYDTPVPGYRNNVVNT']
['222-235', 'LAMPYDTPVPGYRN']
['222-240', 'LAMPYDTPVPGYRNNVVNT']
['223-240', 'AMPYDTPVPGYRNNVVNT']
Uptake (Da)
Time (ms)
0
20
40
60
80
100
$10^3$
$10^5$

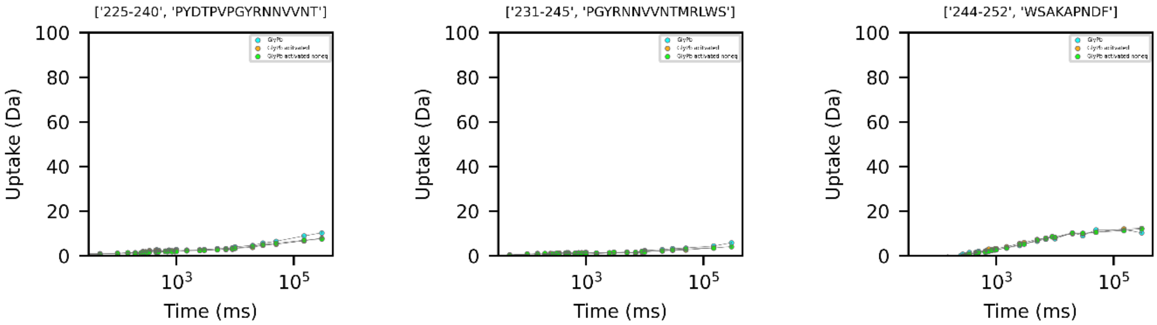
['225-240', 'PYDTPVPGYRNNVVNT']
['231-245', 'PGYRNNVVNTMRLWS']
['244-252', 'WSAKAPNDF']
Uptake (Da)
Time (ms)
100
80
60
40
20
0
$10^3$
$10^5$

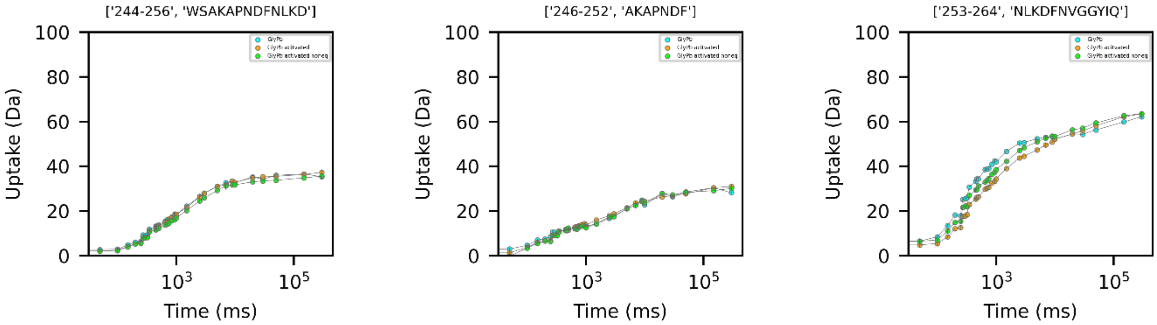
['244-256', 'WSAKAPNDFNLKD']
['246-252', 'AKAPNDF']
['253-264', 'NLKDFNVGGYIQ']
Uptake (Da)
Time (ms)
100
80
60
40
20
0
$10^3$
$10^5$

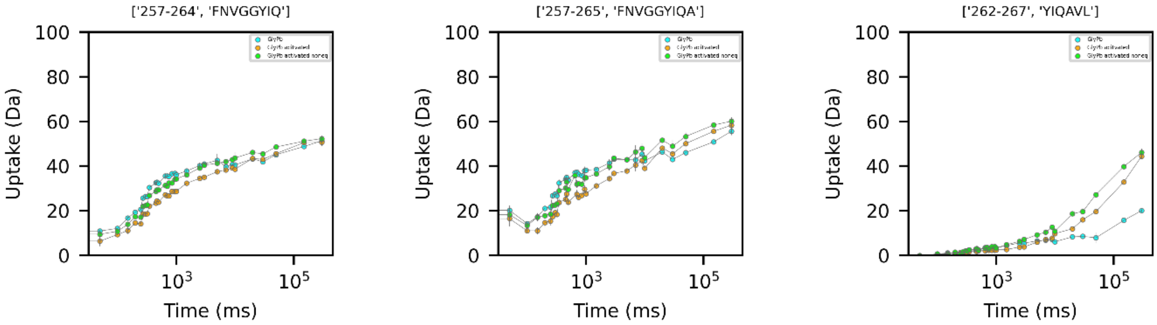
['257-264', 'FNVGGYIQ']
['257-265', 'FNVGGYIQA']
['262-267', 'YIQAVL']
Uptake (Da)
Time (ms)
100
80
60
40
20
0
$10^3$
$10^5$

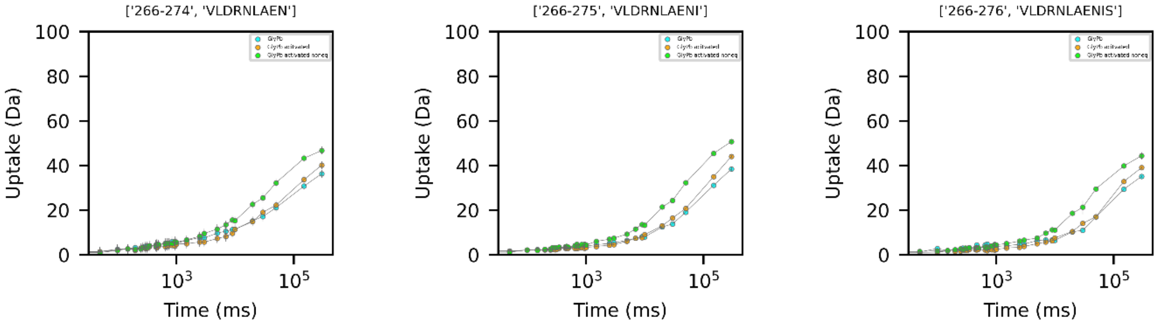
['266-274', 'VLDRNLAEN']
['266-275', 'VLDRNLAENI']
['266-276', 'VLDRNLAENIS']
Uptake (Da)
Time (ms)
100
80
60
40
20
0
$10^3$
$10^5$

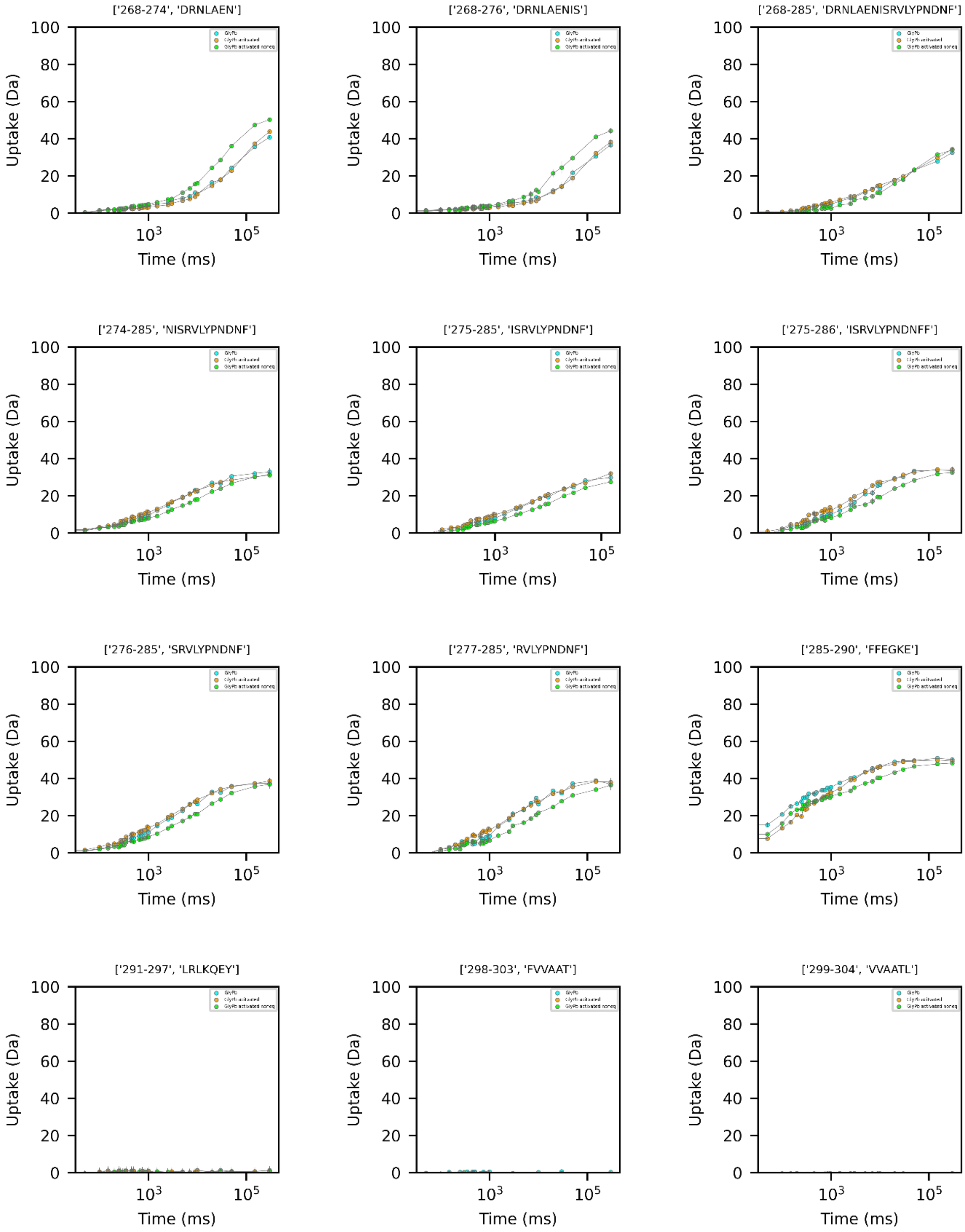
['268-274', 'DRNLAEN']
['268-276', 'DRNLAENIS']
['268-285', 'DRNLAENISRVLYPNDNF']
['274-285', 'NISRVLYPNDNF']
['275-285', 'ISRVLYPNDNF']
['275-286', 'ISRVLYPNDNFF']
['276-285', 'SRVLYPNDNF']
['277-285', 'RVLYPNDNF']
['285-290', 'FFEGKE']
['291-297', 'LRLKQEY']
['298-303', 'FVVAAT']
['299-304', 'VVAATL']
Uptake (Da)
Time (ms)

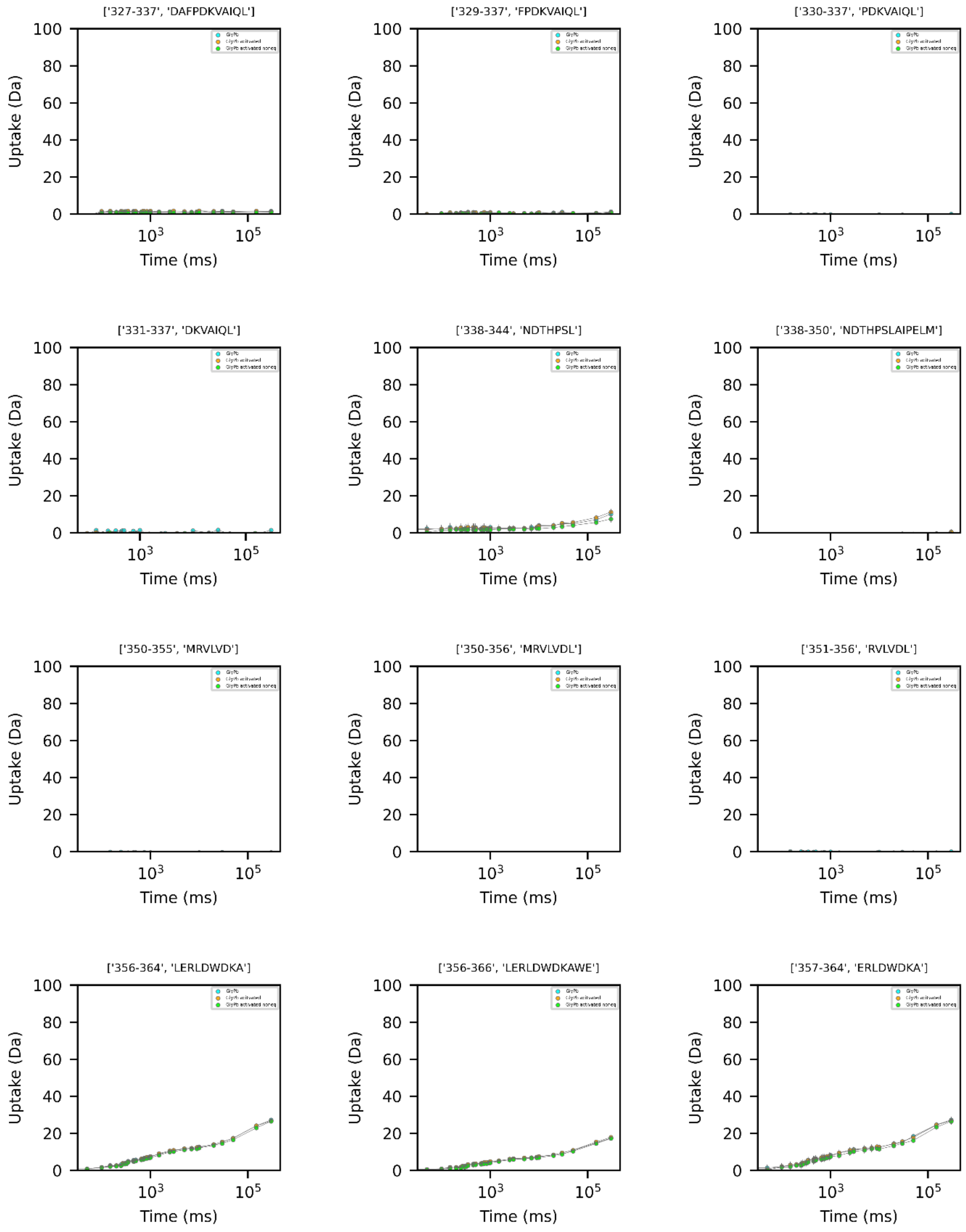
['327-337', 'DAFPDKVAIQL']
['329-337', 'FPDKVAIQL']
['330-337', 'PDKVAIQL']
['331-337', 'DKVAIQL']
['338-344', 'NDTHPSL']
['338-350', 'NDTHPSLAIPELM']
['350-355', 'MRVLVD']
['350-356', 'MRVLVDL']
['351-356', 'RVLVDL']
['356-364', 'LERLDWDKA']
['356-366', 'LERLDWDKAWE']
['357-364', 'ERLDWDKA']
Uptake (Da)
Time (ms)
100
80
60
40
20
0
$10^3$
$10^5$

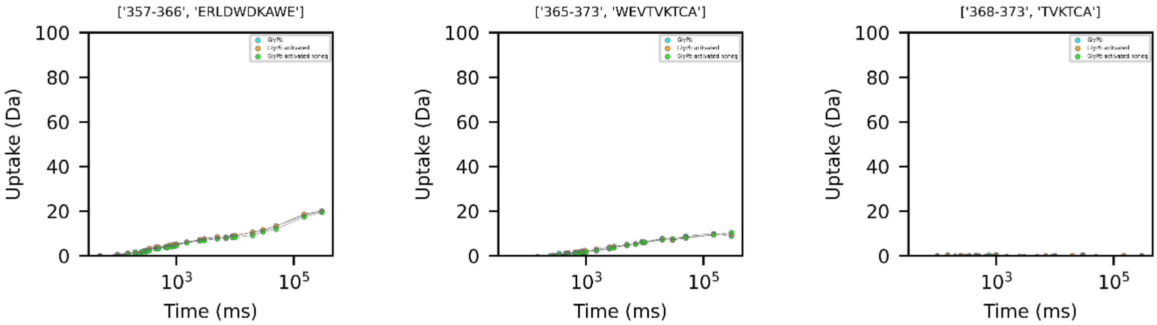


259

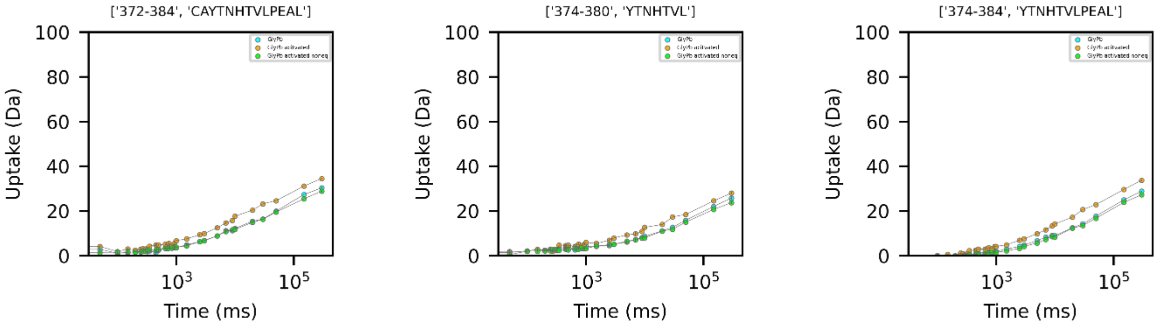


260

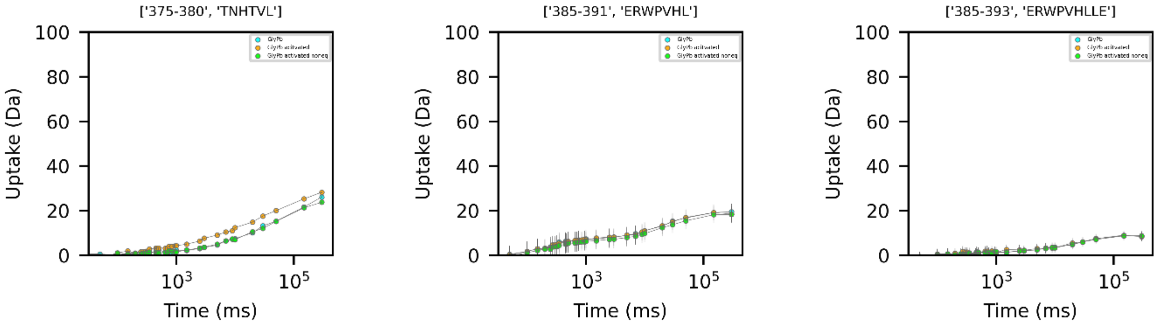


261

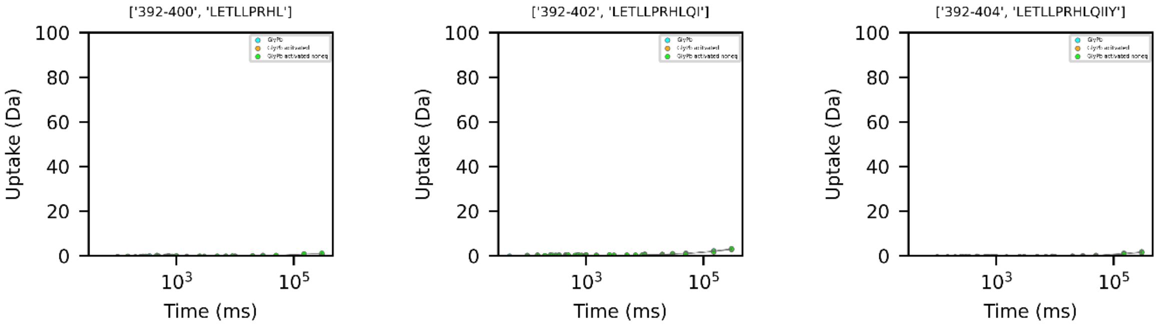


262

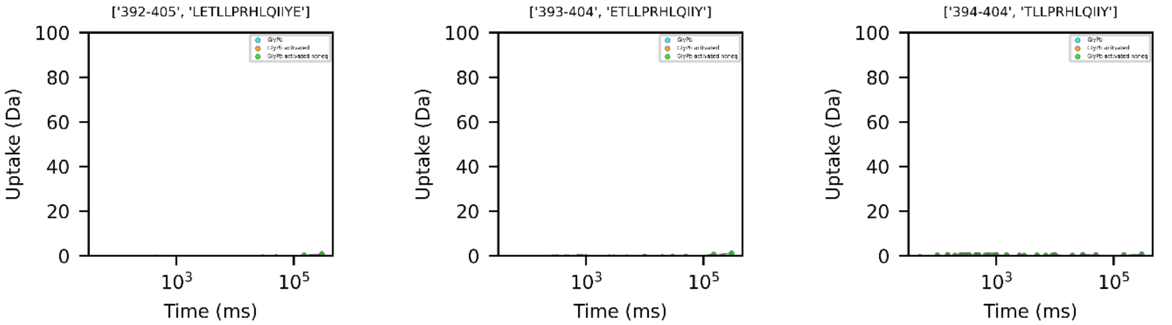
['392-405', 'LETLLPRHLQIIYE']
['393-404', 'ETLLPRHLQIIY']
['394-404', 'TLLPRHLQIIY']
Uptake (Da)
Time (ms)
0
20
40
60
80
100
$10^3$
$10^5$

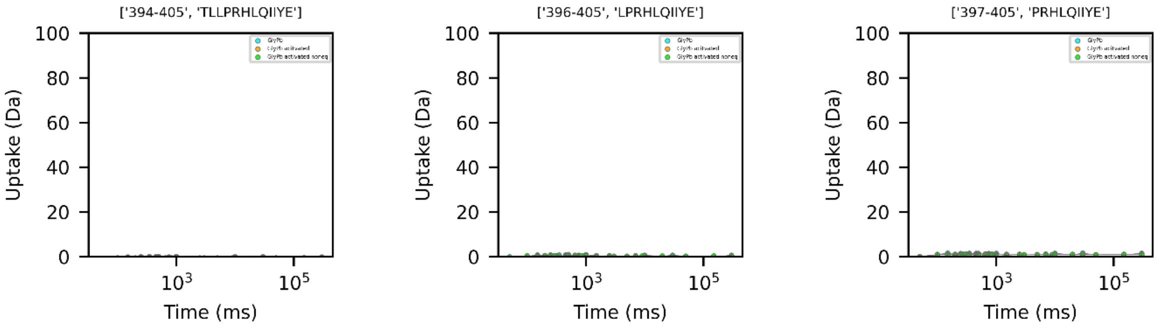
['394-405', 'TLLPRHLQIIYE']
['396-405', 'LPRHLQIIYE']
['397-405', 'PRHLQIIYE']
Uptake (Da)
Time (ms)
0
20
40
60
80
100
$10^3$
$10^5$

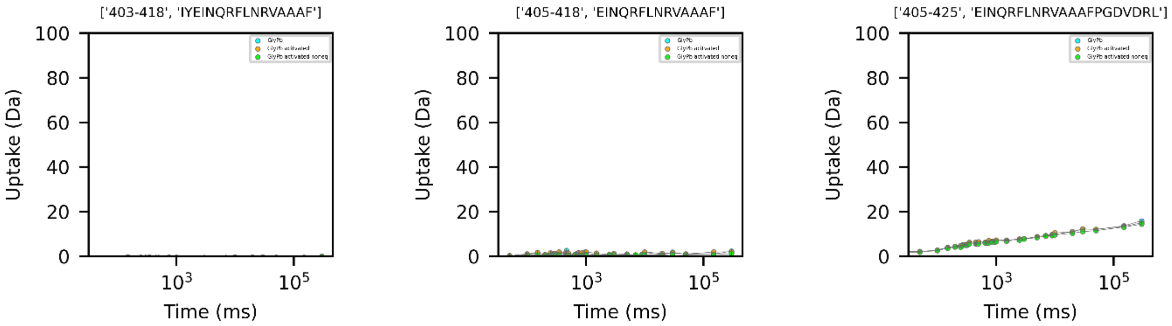
['403-418', 'IYEINQRFLNRVAAAF']
['405-418', 'EINQRFLNRVAAAF']
['405-425', 'EINQRFLNRVAAAFPGDVDRL']
Uptake (Da)
Time (ms)
0
20
40
60
80
100
$10^3$
$10^5$

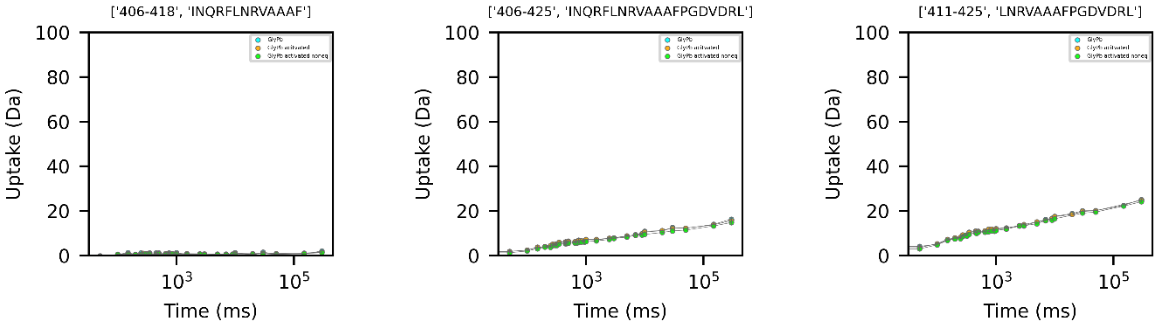
['406-418', 'INQRFLNRVAAAF']
['406-425', 'INQRFLNRVAAAFPGDVDRL']
['411-425', 'LNRVAAAFPGDVDRL']
Uptake (Da)
Time (ms)
0
20
40
60
80
100
$10^3$
$10^5$

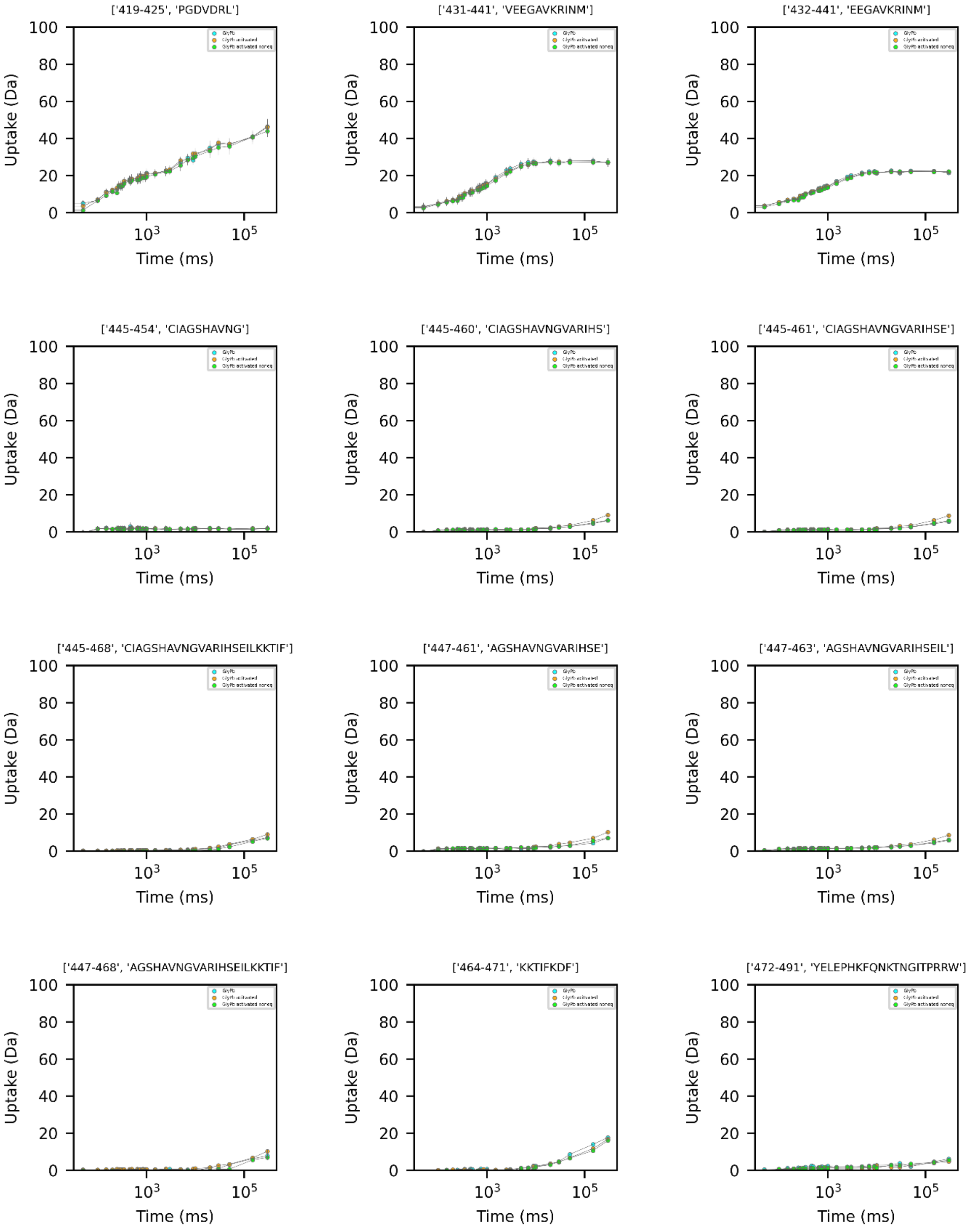
['419-425', 'PGDVDRL']
['431-441', 'VEEGAVKRINM']
['432-441', 'EEGAVKRINM']
['445-454', 'CIAGSHAVNG']
['445-460', 'CIAGSHAVNGVARIHS']
['445-461', 'CIAGSHAVNGVARIHSE']
['445-468', 'CIAGSHAVNGVARIHSEILKKTIF']
['447-461', 'AGSHAVNGVARIHSE']
['447-463', 'AGSHAVNGVARIHSEIL']
['447-468', 'AGSHAVNGVARIHSEILKKTIF']
['464-471', 'KKTIFKDF']
['472-491', 'YELEPHKFQNKTNGITPRRW']
Uptake (Da)
Time (ms)

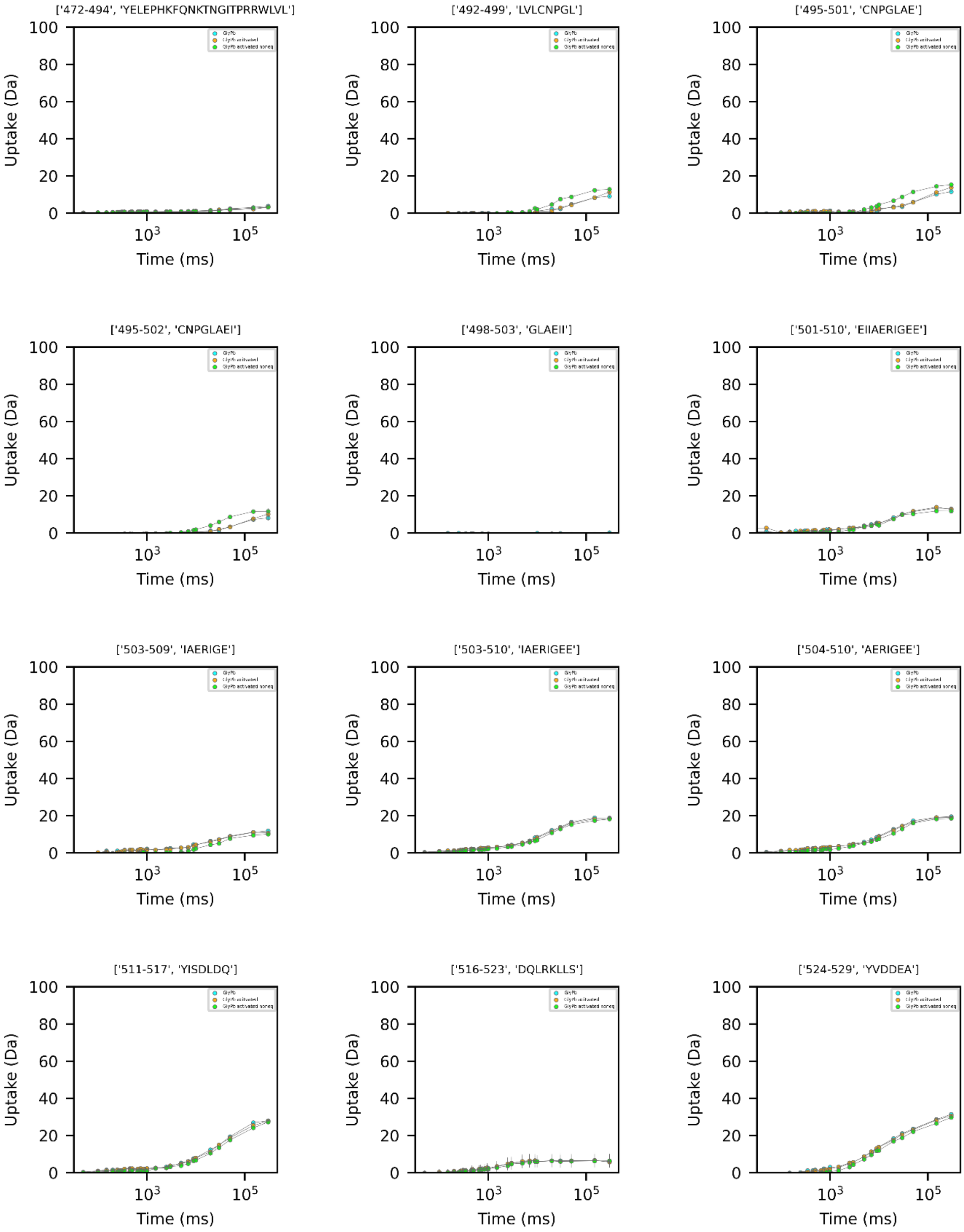
['472-494', 'YELEPHKFQNKTNGITPRRWLVL']
['492-499', 'LVLCNPGL']
['495-501', 'CNPGLAE']
['495-502', 'CNPGLAEI']
['498-503', 'GLAEII']
['501-510', 'EIIAERIGEE']
['503-509', 'IAERIGE']
['503-510', 'IAERIGEE']
['504-510', 'AERIGEE']
['511-517', 'YISDLDQ']
['516-523', 'DQLRKLLS']
['524-529', 'YVDDEA']
Uptake (Da)
Time (ms)
0
20
40
60
80
100
$10^3$
$10^5$

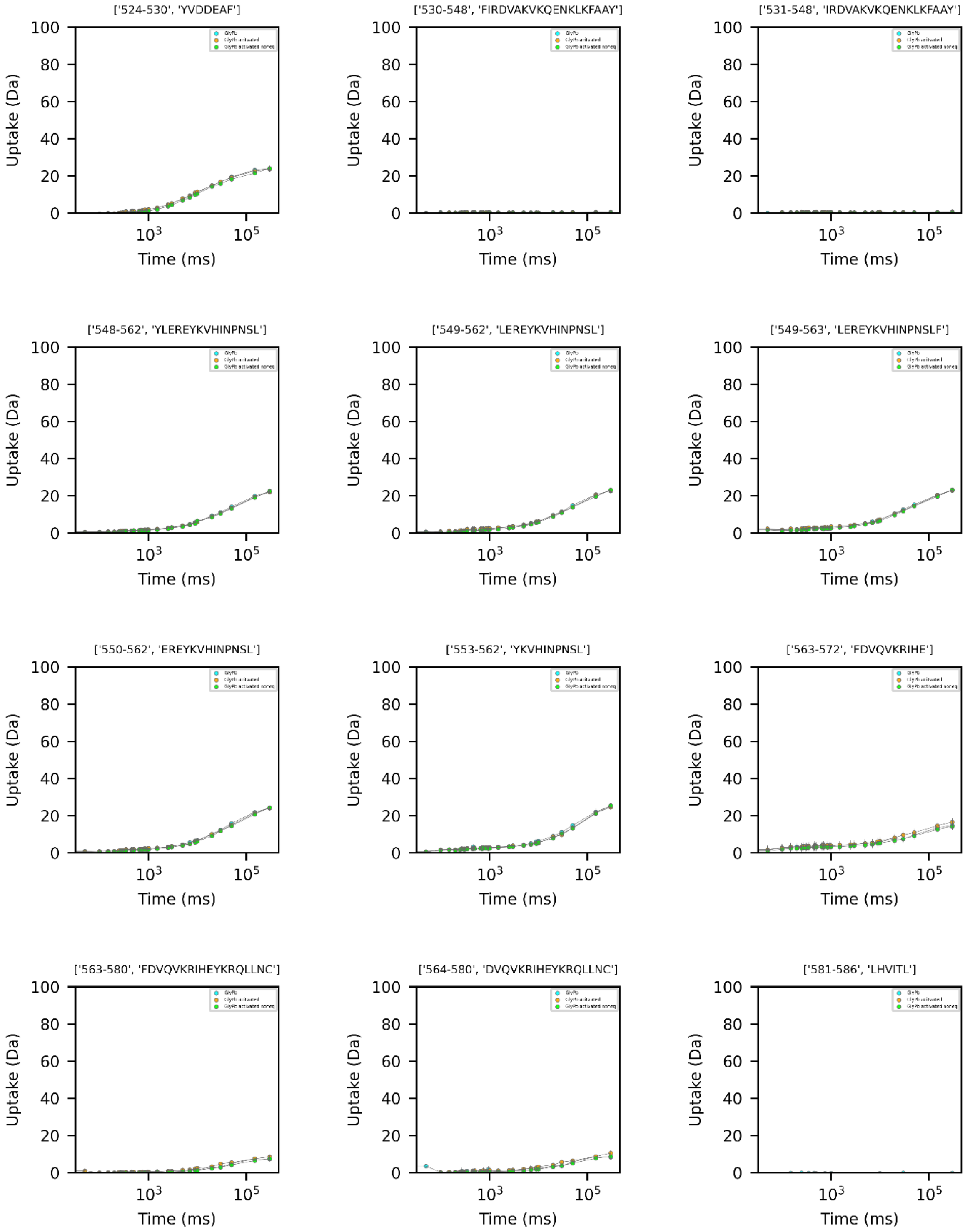
['524-530', 'YVDDEAF']
['530-548', 'FIRDVAKVKQENKLKFAAY']
['531-548', 'IRDVAKVKQENKLKFAAY']
['548-562', 'YLEREYKVHINPNSL']
['549-562', 'LEREYKVHINPNSL']
['549-563', 'LEREYKVHINPNSLF']
['550-562', 'EREYKVHINPNSL']
['553-562', 'YKVHINPNSL']
['563-572', 'FDVQVKRIHE']
['563-580', 'FDVQVKRIHEYKRQLLNC']
['564-580', 'DVQVKRIHEYKRQLLNC']
['581-586', 'LHVITL']
Uptake (Da)
Time (ms)
0
20
40
60
80
100
$10^3$
$10^5$

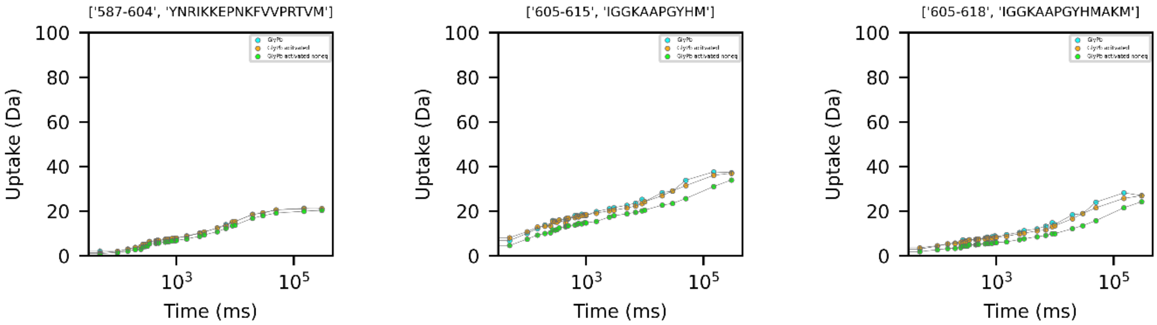
['587-604', 'YNRIKKEPNKFVVPRTVM']
['605-615', 'IGGKAAPGYHM']
['605-618', 'IGGKAAPGYHMAKM']
Uptake (Da)
Time (ms)

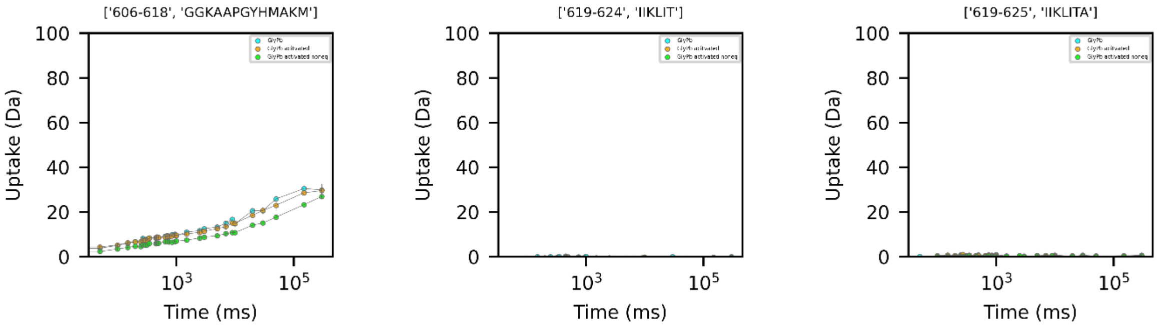
['606-618', 'GGKAAPGYHMAKM']
['619-624', 'IIKLIT']
['619-625', 'IIKLITA']
Uptake (Da)
Time (ms)

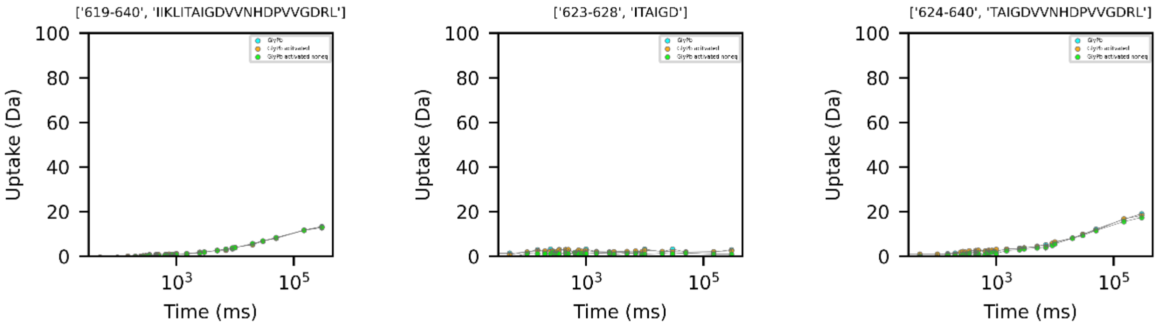
['619-640', 'IIKLITAIGDVVNHDPVVGDRL']
['623-628', 'ITAIGD']
['624-640', 'TAIGDVVNHDPVVGDRL']
Uptake (Da)
Time (ms)

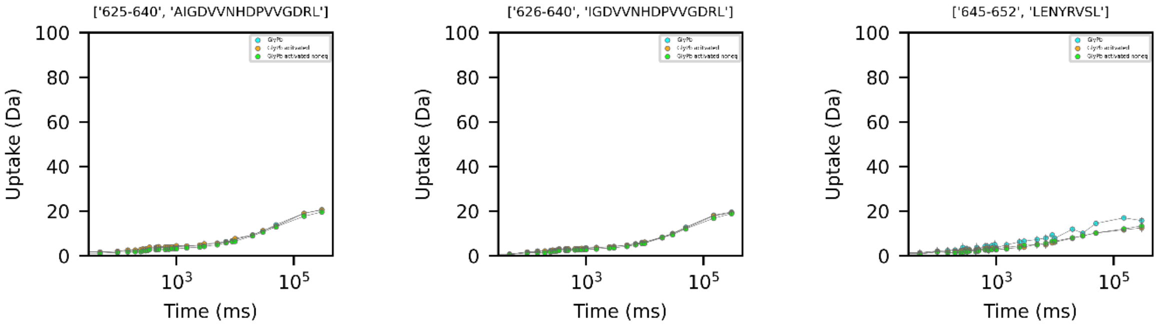
['625-640', 'AIGDVVNHDPVVGDRL']
['626-640', 'IGDVVNHDPVVGDRL']
['645-652', 'LENYRVSL']
Uptake (Da)
Time (ms)

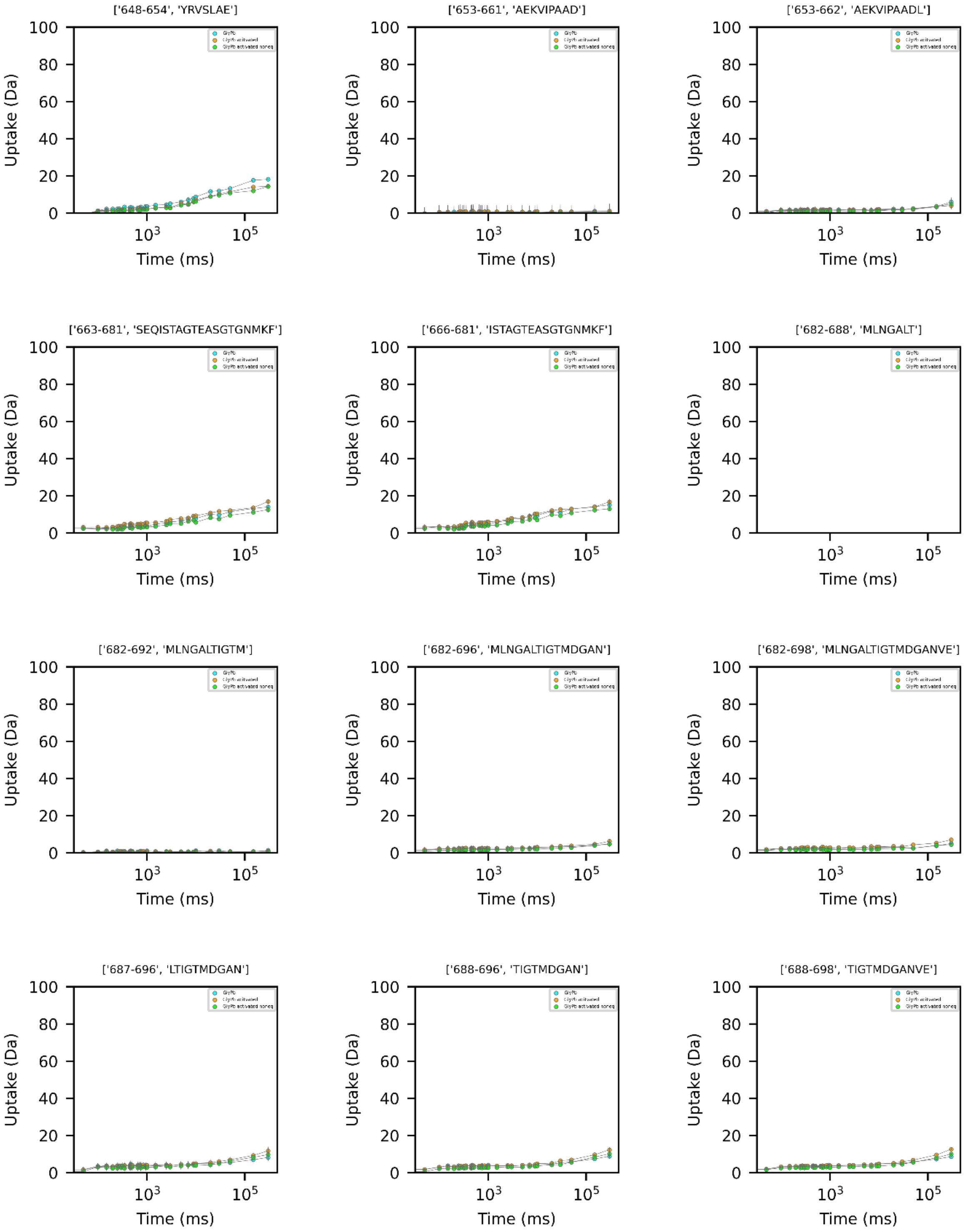
['648-654', 'YRVSLAE']
['653-661', 'AEKVIPAAD']
['653-662', 'AEKVIPAADL']
['663-681', 'SEQISTAGTEASGTGNMKF']
['666-681', 'ISTAGTEASGTGNMKF']
['682-688', 'MLNGALT']
['682-692', 'MLNGALTIGTM']
['682-696', 'MLNGALTIGTMDGAN']
['682-698', 'MLNGALTIGTMDGANVE']
['687-696', 'LTIGTMDGAN']
['688-696', 'TIGTMDGAN']
['688-698', 'TIGTMDGANVE']
Uptake (Da)
Time (ms)
0
20
40
60
80
100
$10^3$
$10^5$

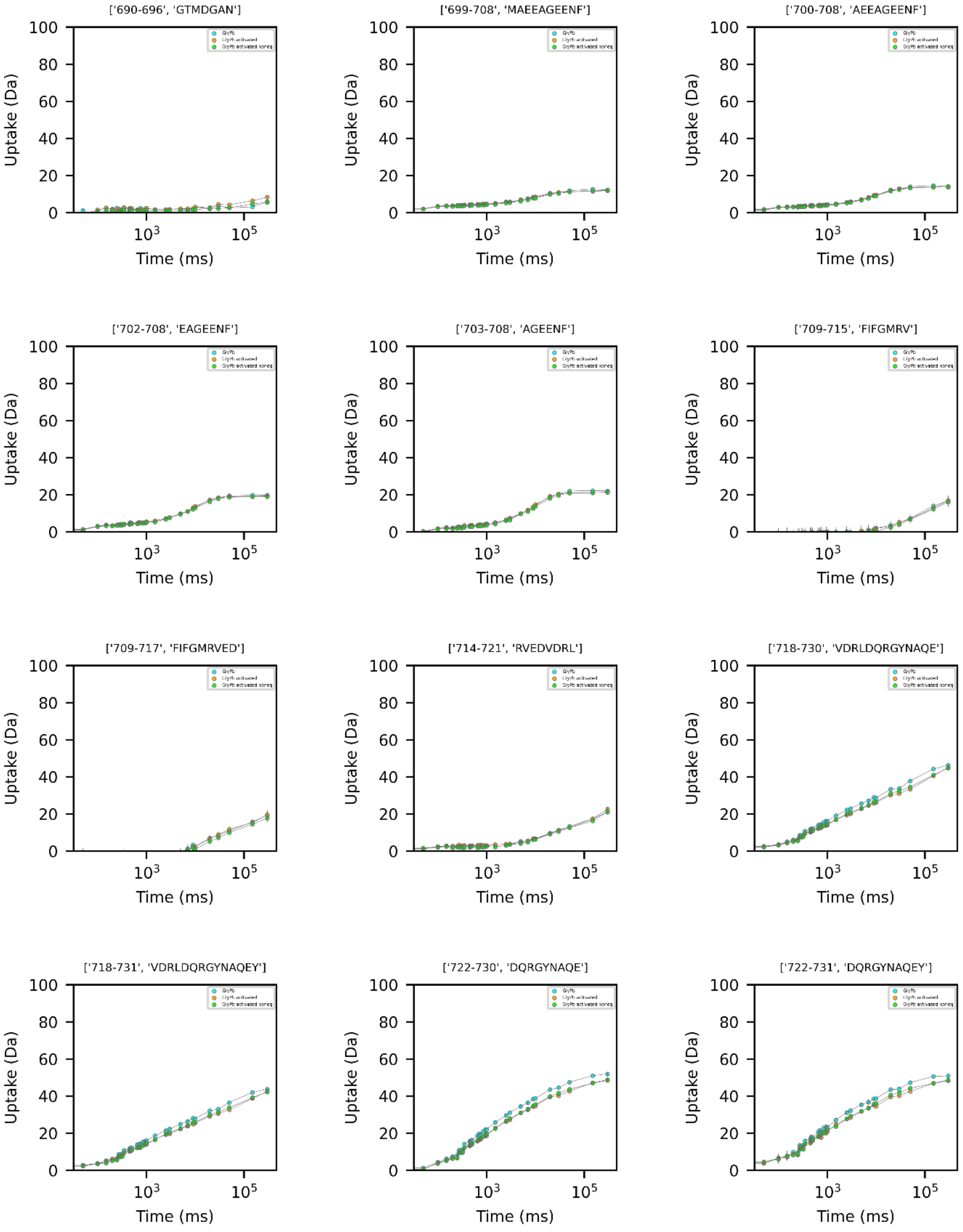
['690-696', 'GTMDGAN']
['699-708', 'MAEEAGEENF']
['700-708', 'AEEAGEENF']
['702-708', 'EAGEENF']
['703-708', 'AGEENF']
['709-715', 'FIFGMRV']
['709-717', 'FIFGMRVED']
['714-721', 'RVEDVDRL']
['718-730', 'VDRLDQRGYNAQE']
['718-731', 'VDRLDQRGYNAQEY']
['722-730', 'DQRGYNAQE']
['722-731', 'DQRGYNAQEY']
Uptake (Da)
Time (ms)

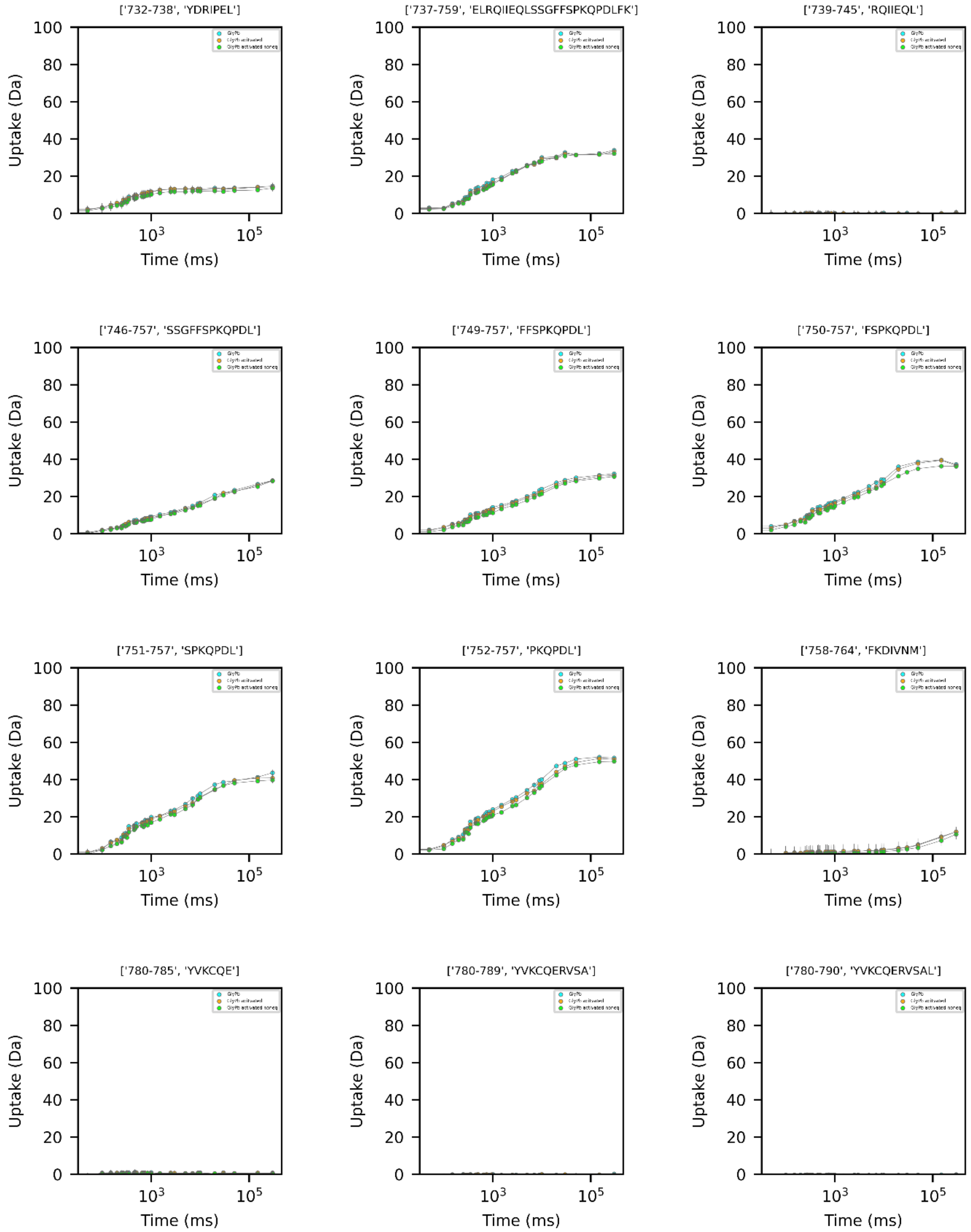
['732-738', 'YDRIPEL']
['737-759', 'ELRQIIEQLSSGFFSPKQPDLFK']
['739-745', 'RQIIEQL']
['746-757', 'SSGFFSPKQPDL']
['749-757', 'FFSPKQPDL']
['750-757', 'FSPKQPDL']
['751-757', 'SPKQPDL']
['752-757', 'PKQPDL']
['758-764', 'FKDIVNM']
['780-785', 'YVKCQE']
['780-789', 'YVKCQERVSA']
['780-790', 'YVKCQERVSAL']
Uptake (Da)
Time (ms)
$10^3$
$10^5$

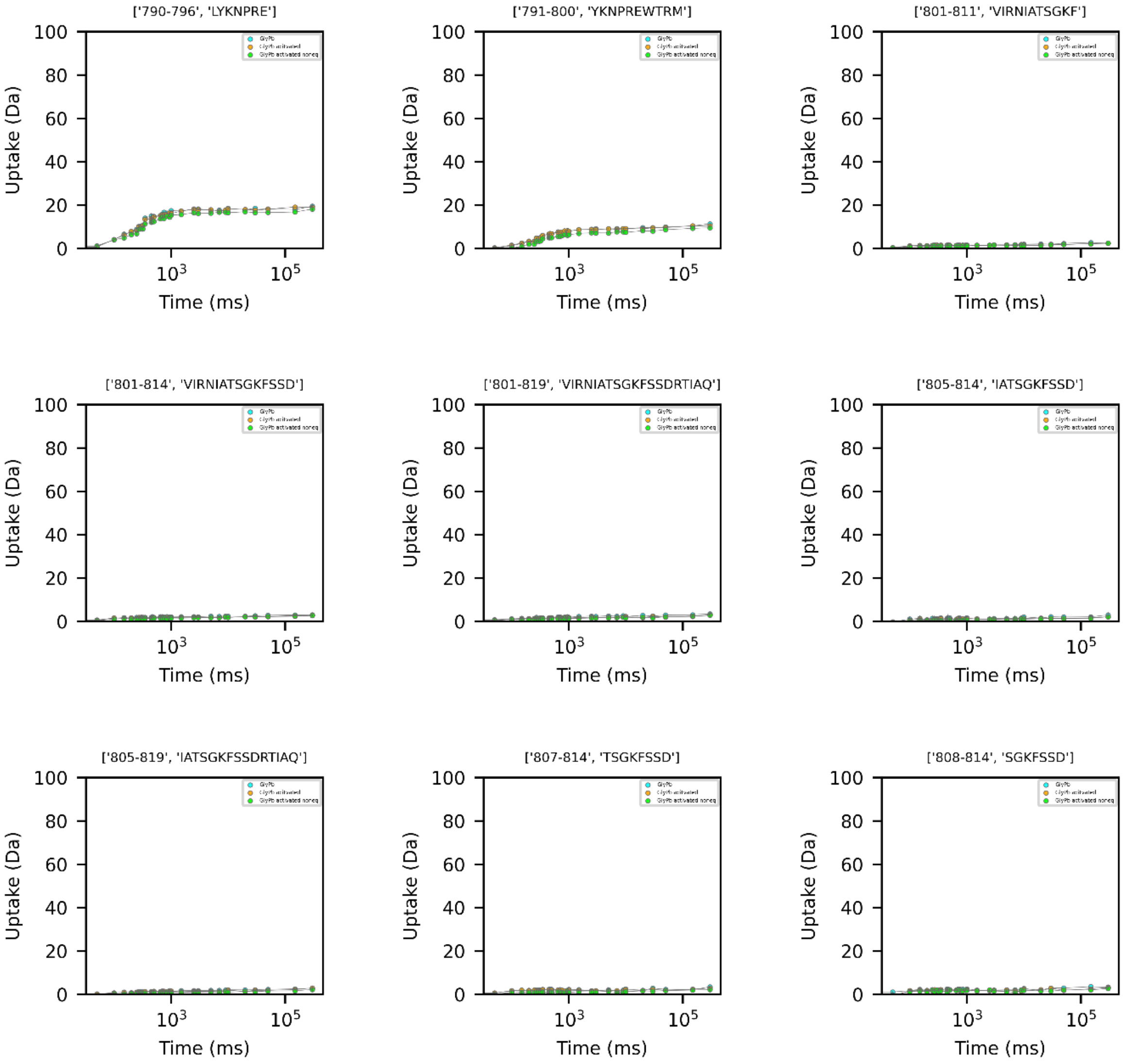
['790-796', 'LYKNPRE']
['791-800', 'YKNPREWTRM']
['801-811', 'VIRNIATSGKF']
['801-814', 'VIRNIATSGKFSSD']
['801-819', 'VIRNIATSGKFSSDRTIAQ']
['805-814', 'IATSGKFSSD']
['805-819', 'IATSGKFSSDRTIAQ']
['807-814', 'TSGKFSSD']
['808-814', 'SGKFSSD']
Uptake (Da)
Time (ms)
0
20
40
60
80
100
$10^3$
$10^5$

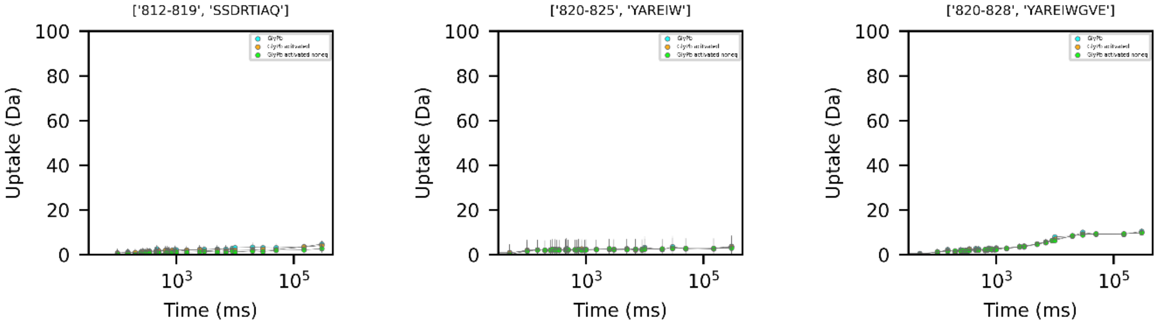
['812-819', 'SSDRTIAQ']
['820-825', 'YAREIW']
['820-828', 'YAREIWGVE']
Uptake (Da)
Time (ms)
100
80
60
40
20
0
$10^3$
$10^5$

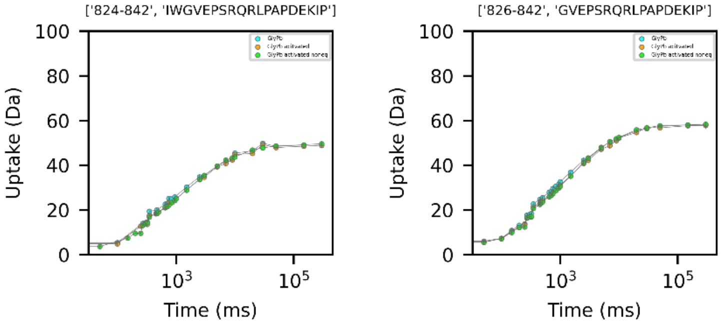
['824-842', 'IWGVEPSRQRLPAPDEKIP']
['826-842', 'GVEPSRQRLPAPDEKIP']
Uptake (Da)
Time (ms)
100
80
60
40
20
0
$10^3$
$10^5$

Figure S17. HDX uptake curves of GlyPa per peptide. Absolute data for three protein states shown (blue – apo GlyPa; orange – caffeine bound activated GlyPa; green – caffeine bound non-equilibrium activated GlyPa). Peptide sequence and amino acid number given above the plot. Y-axis shows absolute % deuteration at labeling time (ms) on X-axis.

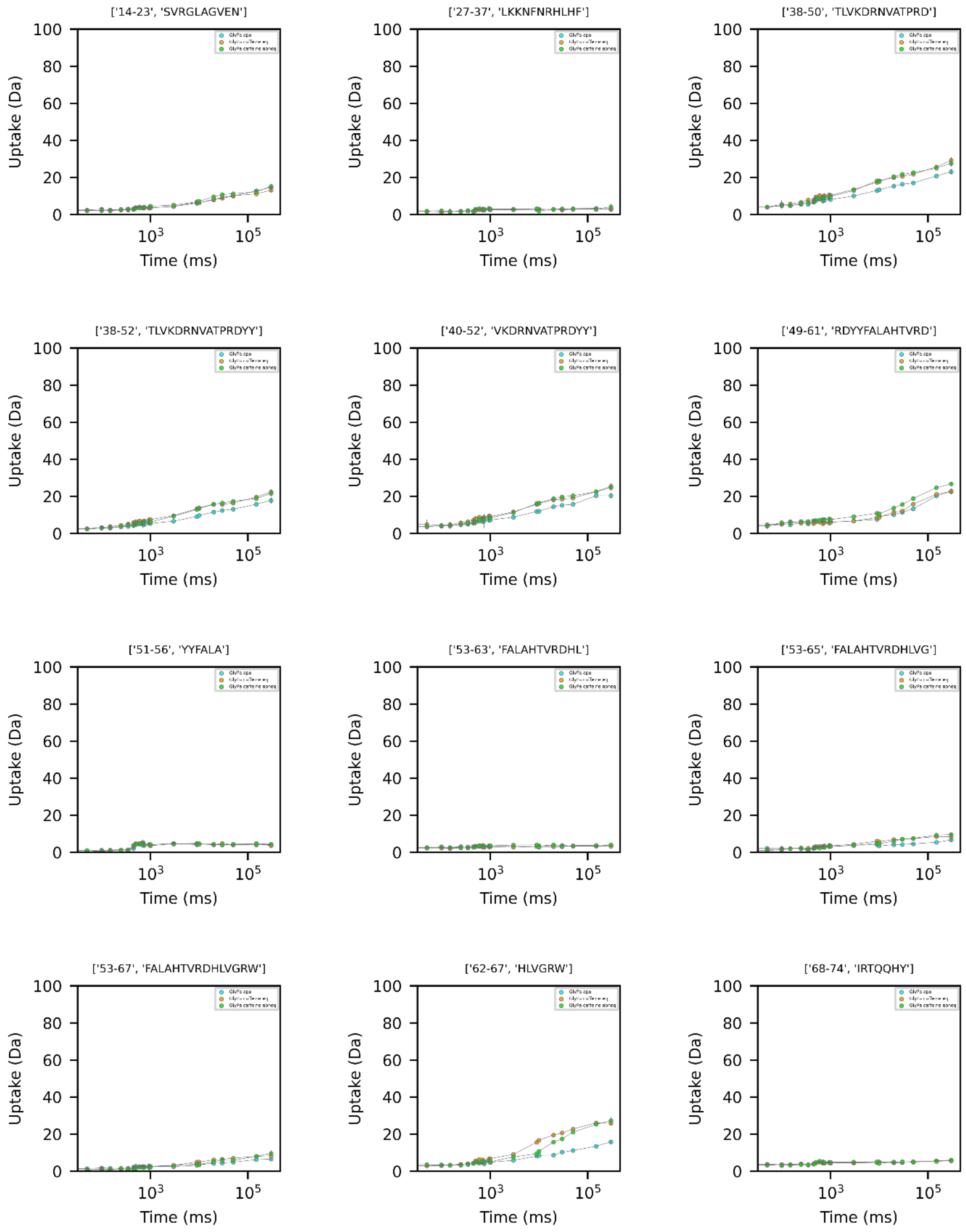

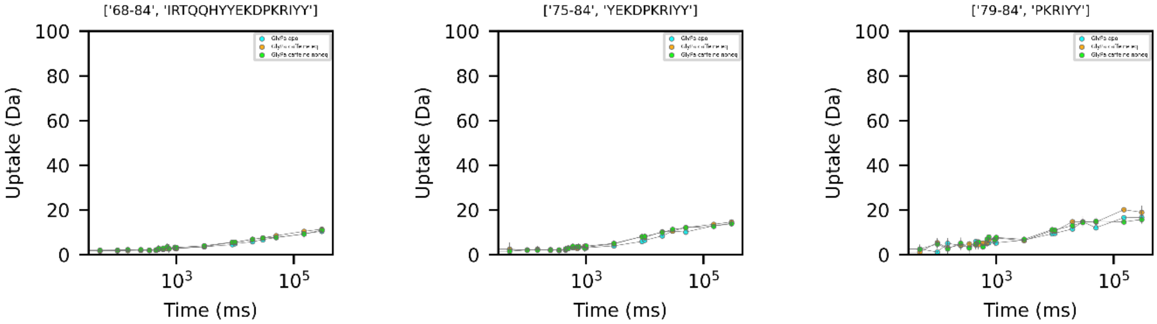
['68-84', 'IRTQQHYYEKDPKRIYY']
['75-84', 'YEKDPKRIYY']
['79-84', 'PKRIYY']
Uptake (Da)
Time (ms)
0
20
40
60
80
100
$10^3$
$10^5$

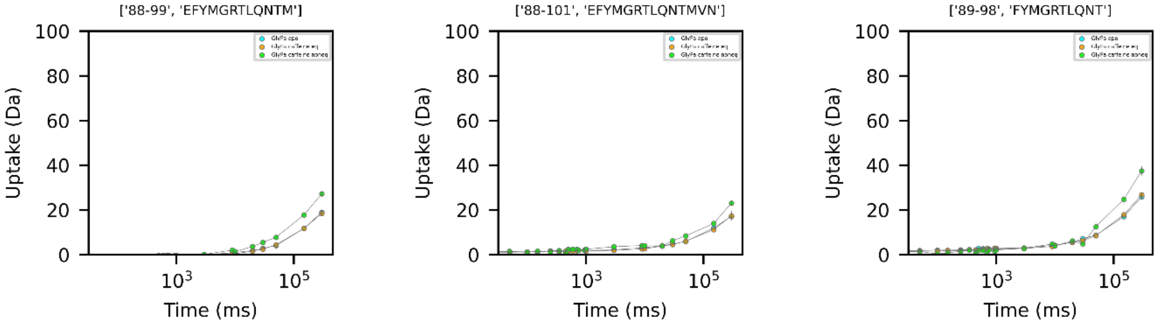
['88-99', 'EFYMGRTLQNTM']
['88-101', 'EFYMGRTLQNTMVN']
['89-98', 'FYMGRTLQNT']
Uptake (Da)
Time (ms)
0
20
40
60
80
100
$10^3$
$10^5$

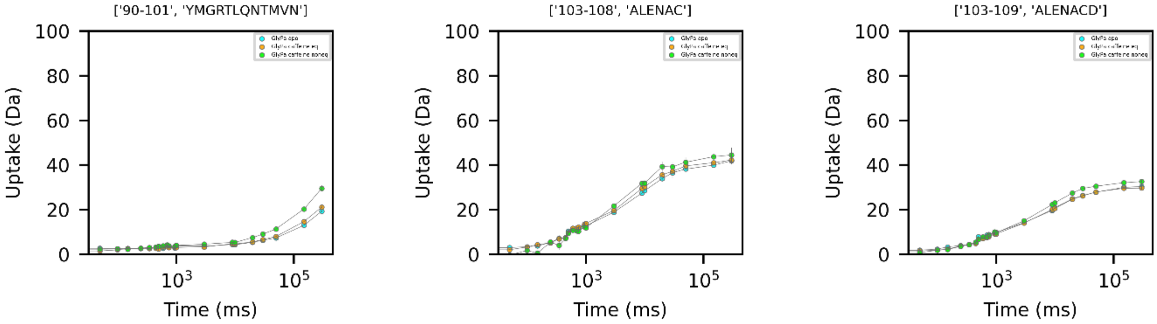
['90-101', 'YMGRTLQNTMVN']
['103-108', 'ALENAC']
['103-109', 'ALENACD']
Uptake (Da)
Time (ms)
0
20
40
60
80
100
$10^3$
$10^5$

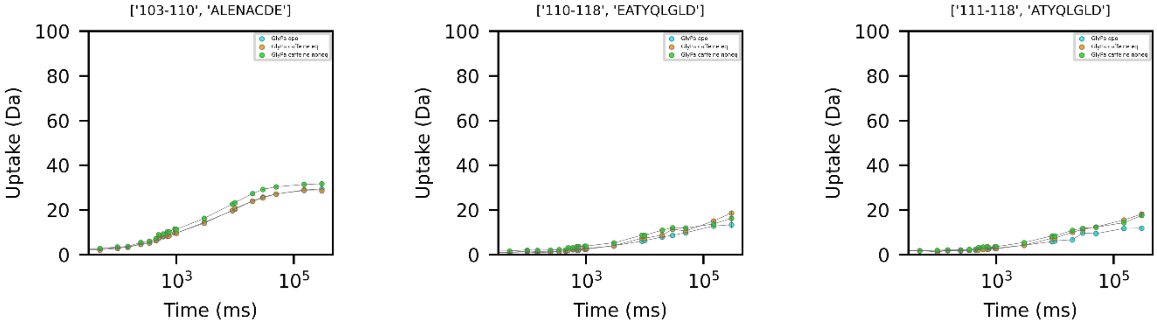
['103-110', 'ALENACDE']
['110-118', 'EATYQLGLD']
['111-118', 'ATYQLGLD']
Uptake (Da)
Time (ms)
0
20
40
60
80
100
$10^3$
$10^5$

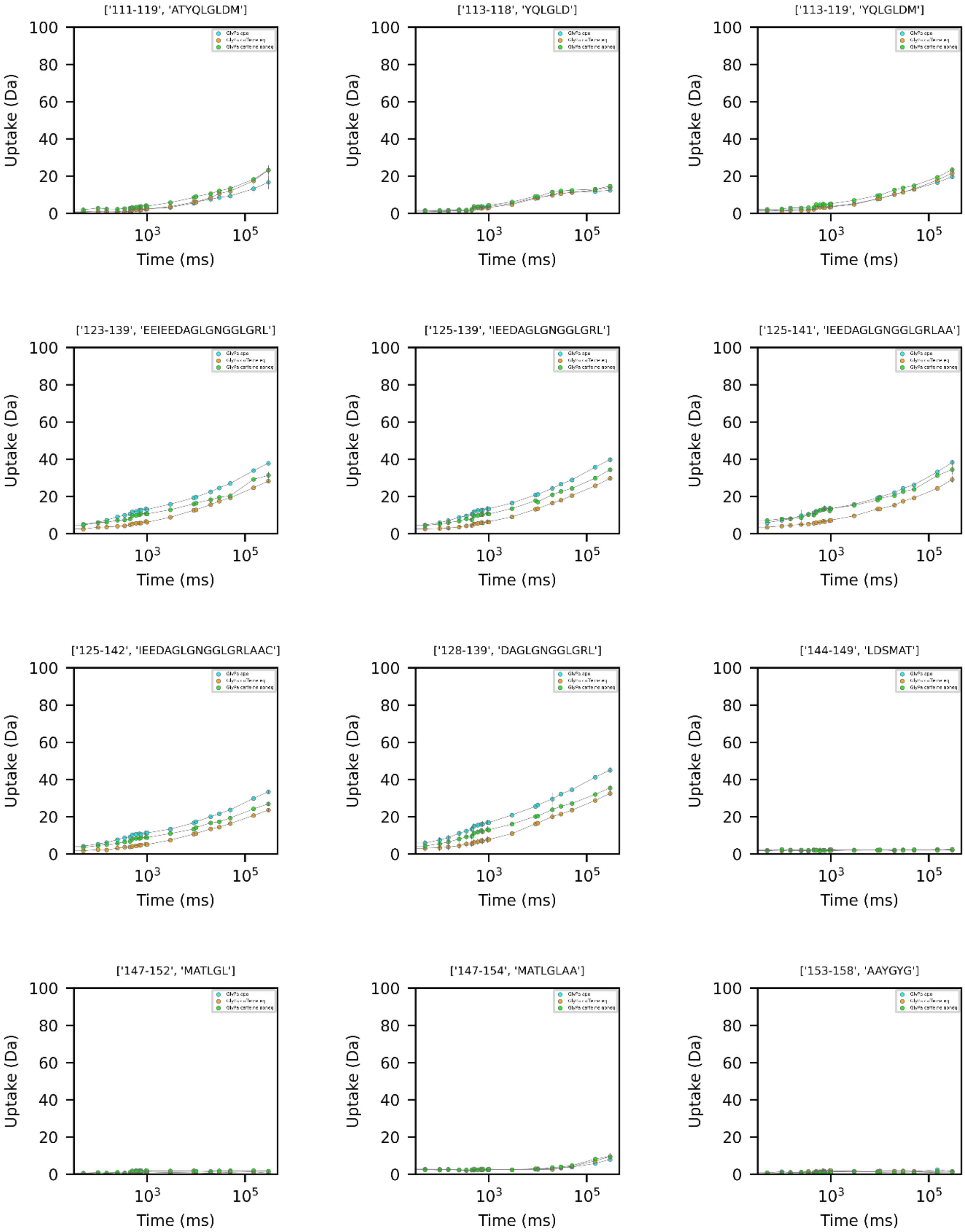
['111-119', 'ATYQLGLDM']
['113-118', 'YQLGLD']
['113-119', 'YQLGLDM']
['123-139', 'EEIEEDAGLGNGGLGRL']
['125-139', 'IEEDAGLGNGGLGRL']
['125-141', 'IEEDAGLGNGGLGRLAA']
['125-142', 'IEEDAGLGNGGLGRLAAC']
['128-139', 'DAGLGNGGLGRL']
['144-149', 'LDSMAT']
['147-152', 'MATLGL']
['147-154', 'MATLGLAA']
['153-158', 'AAYGYG']
Uptake (Da)
Time (ms)

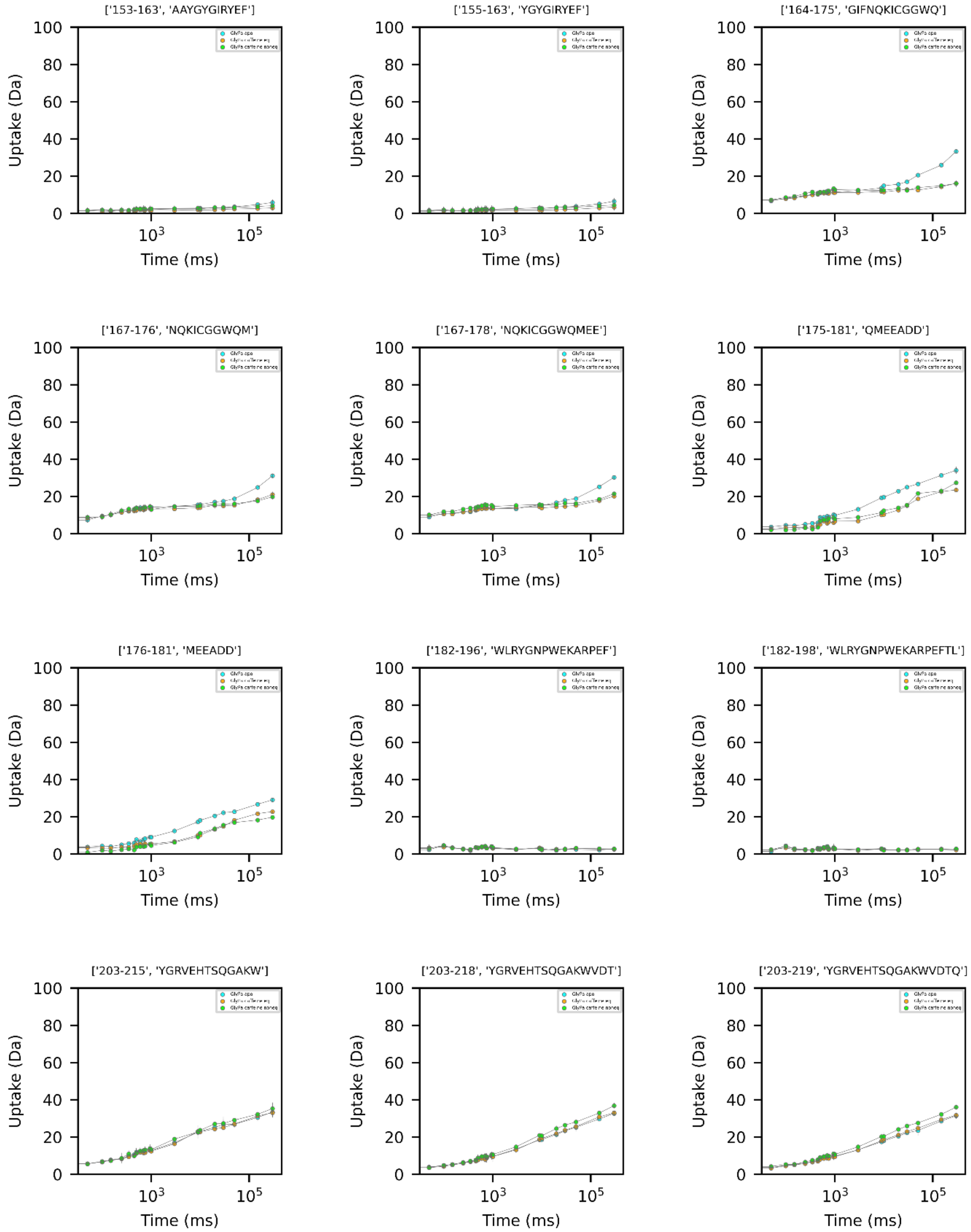
['153-163', 'AAYGYGIRYEF']
['155-163', 'YGYGIRYEF']
['164-175', 'GIFNQKICGGWQ']
['167-176', 'NQKICGGWQM']
['167-178', 'NQKICGGWQMEE']
['175-181', 'QMEEADD']
['176-181', 'MEEADD']
['182-196', 'WLRYGNPWEKARPEF']
['182-198', 'WLRYGNPWEKARPEFTL']
['203-215', 'YGRVEHTSQGAKW']
['203-218', 'YGRVEHTSQGAKWVDT']
['203-219', 'YGRVEHTSQGAKWVDTQ']
Uptake (Da)
Time (ms)

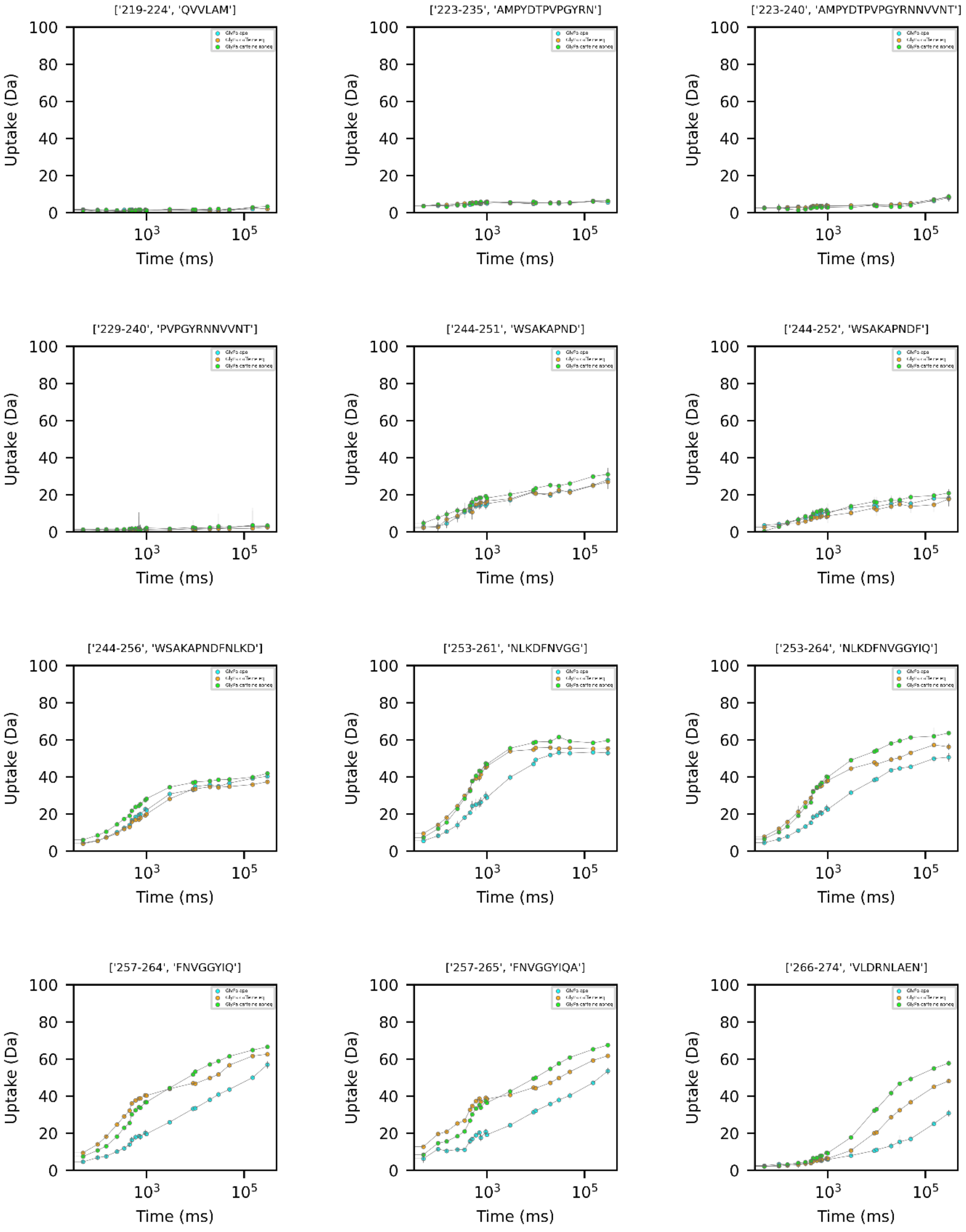
['219-224', 'QVVLAM']
['223-235', 'AMPYDTPVPGYRN']
['223-240', 'AMPYDTPVPGYRNNVVNT']
['229-240', 'PVPGYRNNVVNT']
['244-251', 'WSAKAPND']
['244-252', 'WSAKAPNDF']
['244-256', 'WSAKAPNDFNLKD']
['253-261', 'NLKDFNVGG']
['253-264', 'NLKDFNVGGYIQ']
['257-264', 'FNVGGYIQ']
['257-265', 'FNVGGYIQA']
['266-274', 'VLDRNLAEN']
Uptake (Da)
Time (ms)

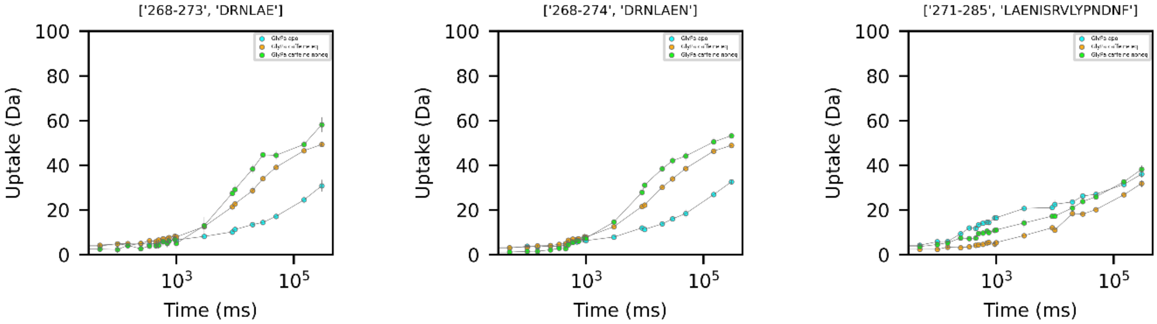
['268-273', 'DRNLAE']
['268-274', 'DRNLAEN']
['271-285', 'LAENISRVLYPNDNF']
Uptake (Da)
Time (ms)

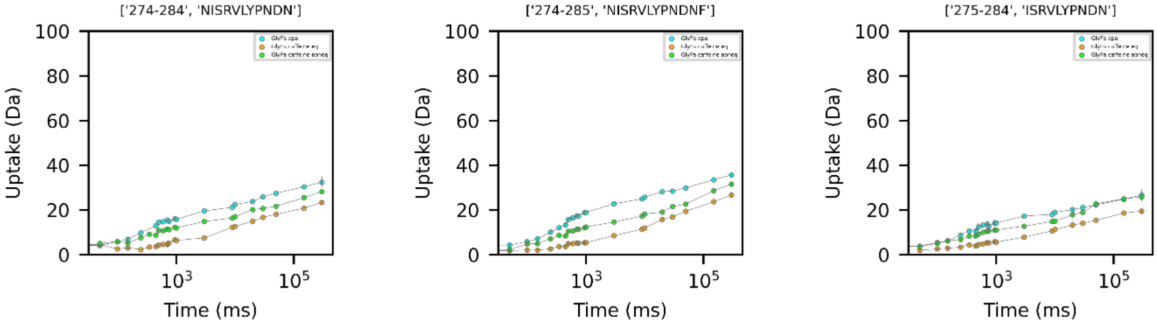
['274-284', 'NISRVLYPNDN']
['274-285', 'NISRVLYPNDNF']
['275-284', 'ISRVLYPNDN']
Uptake (Da)
Time (ms)

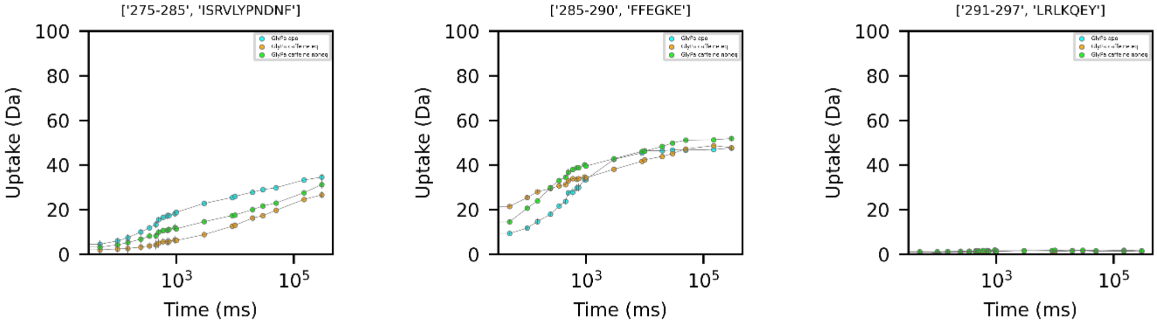
['275-285', 'ISRVLYPNDNF']
['285-290', 'FFEGKE']
['291-297', 'LRLKQEY']
Uptake (Da)
Time (ms)

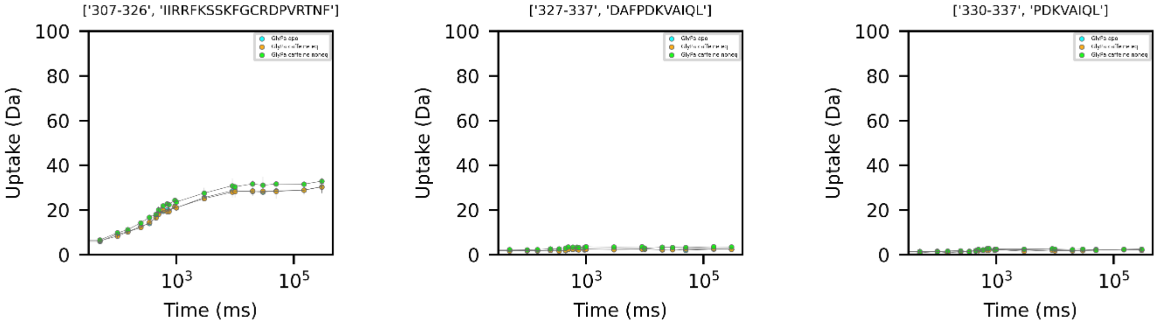
['307-326', 'IIRRFKSSKFGCRDPVRTNF']
['327-337', 'DAFPDKVAIQL']
['330-337', 'PDKVAIQL']
Uptake (Da)
Time (ms)

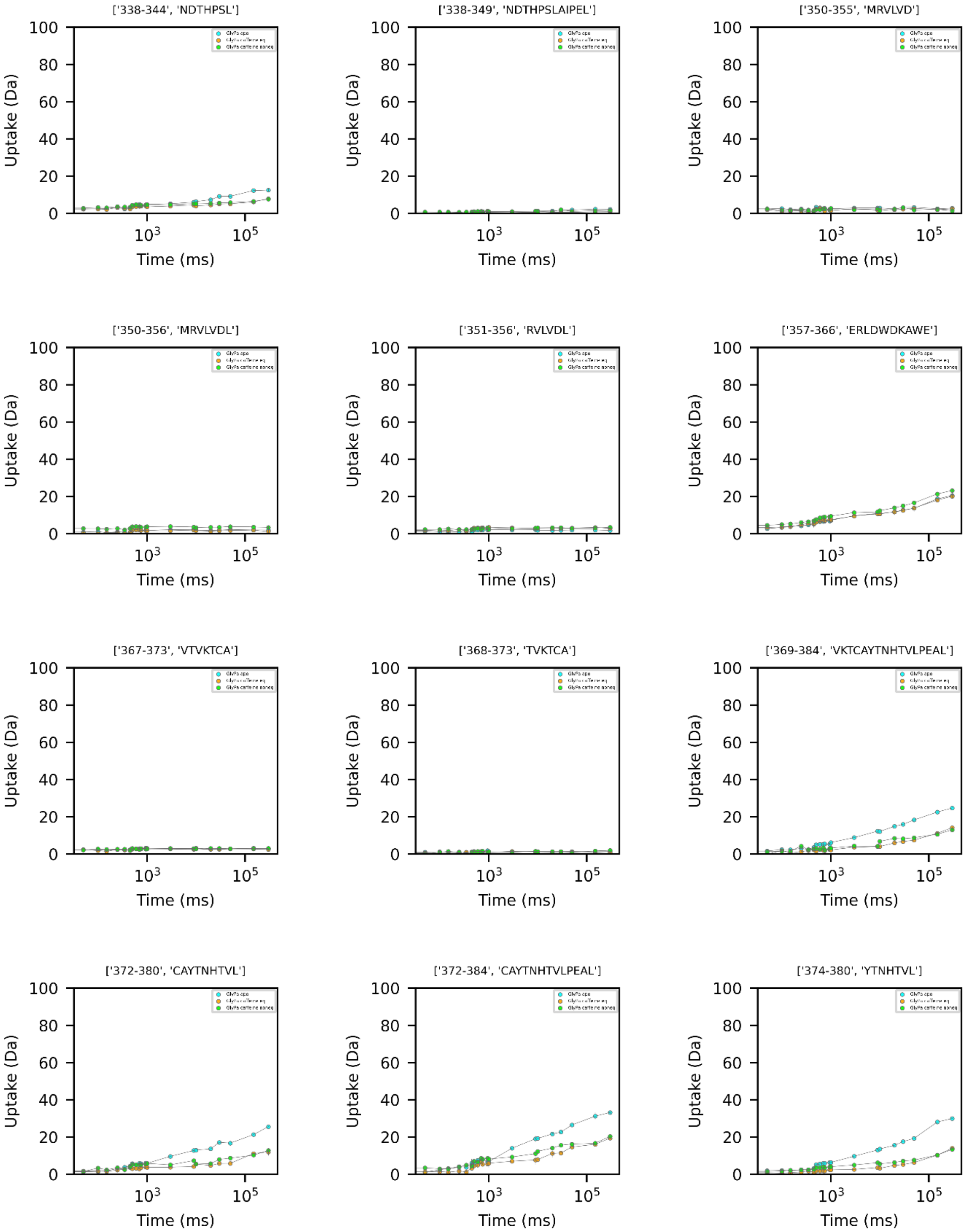

['338-344', 'NDTHPSL']
['338-349', 'NDTHPSLAIPEL']
['350-355', 'MRVLVD']
['350-356', 'MRVLVDL']
['351-356', 'RVLVDL']
['357-366', 'ERLDWDKAWE']
['367-373', 'VTVKTCA']
['368-373', 'TVKTCA']
['369-384', 'VKTCAYTNHTVLPEAL']
['372-380', 'CAYTNHTVL']
['372-384', 'CAYTNHTVLPEAL']
['374-380', 'YTNHTVL']
Uptake (Da)
Time (ms)
0
20
40
60
80
100
$10^3$
$10^5$

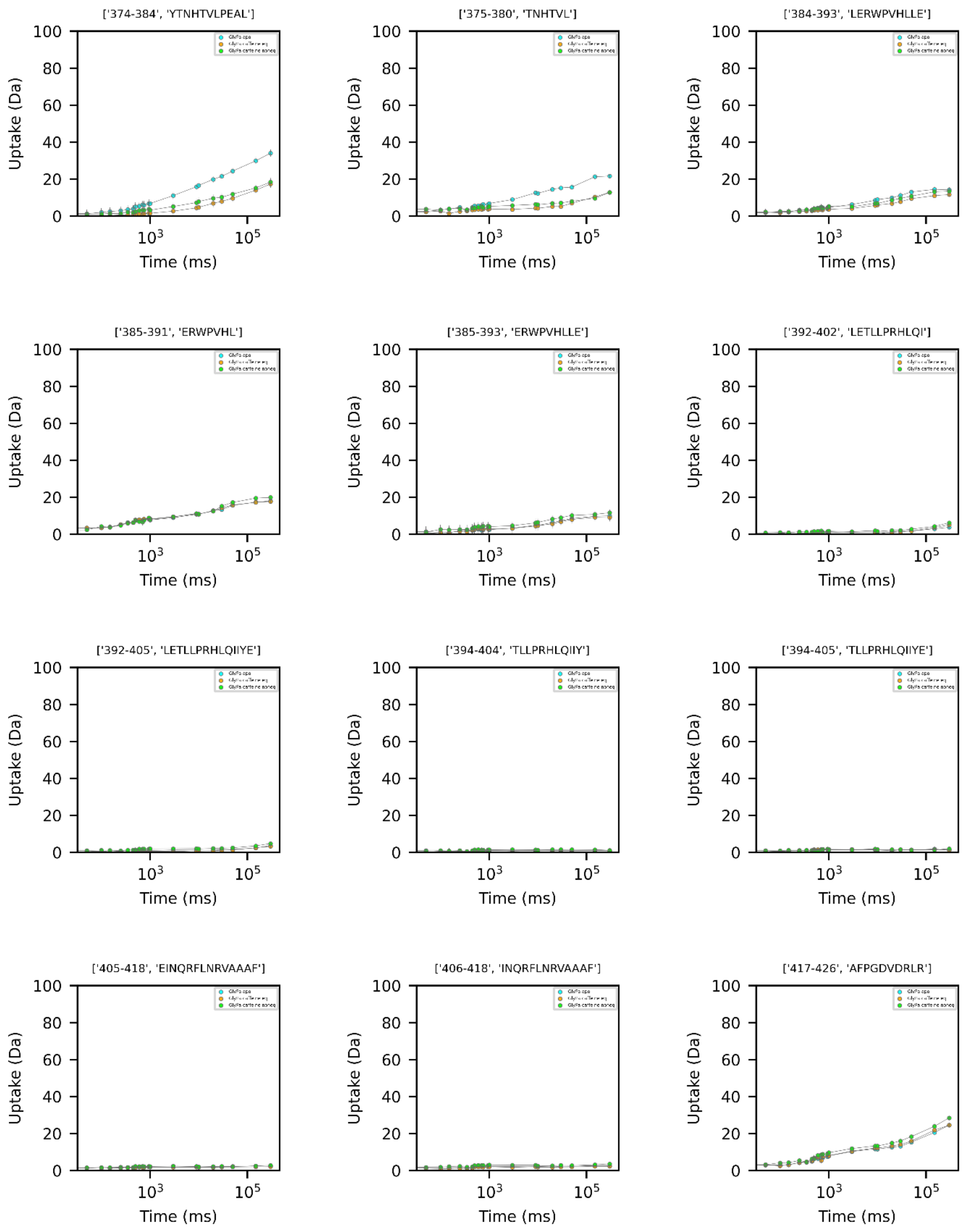
['374-384', 'YTNHTVLPEAL']
['375-380', 'TNHTVL']
['384-393', 'LERWPVHLLE']
['385-391', 'ERWPVHL']
['385-393', 'ERWPVHLLE']
['392-402', 'LETLLPRHLQI']
['392-405', 'LETLLPRHLQIIYE']
['394-404', 'TLLPRHLQIIY']
['394-405', 'TLLPRHLQIIYE']
['405-418', 'EINQRFLNRVAAAF']
['406-418', 'INQRFLNRVAAAF']
['417-426', 'AFPGDVDRLR']
Uptake (Da)
Time (ms)
0
20
40
60
80
100
$10^3$
$10^5$

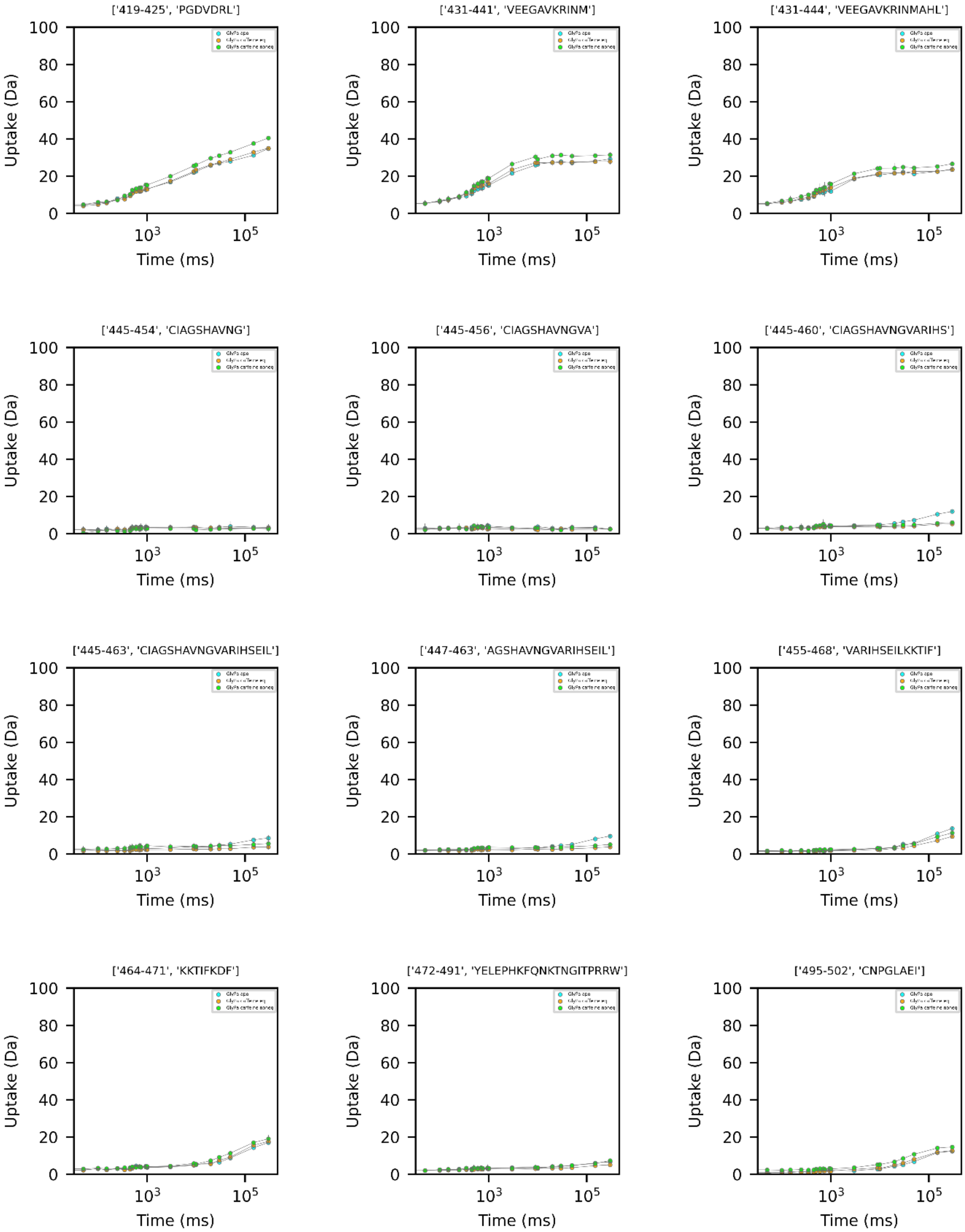
['419-425', 'PGDVDRL']
['431-441', 'VEEGAVKRINM']
['431-444', 'VEEGAVKRINMAHL']
['445-454', 'CIAGSHAVNG']
['445-456', 'CIAGSHAVNGVA']
['445-460', 'CIAGSHAVNGVARIHS']
['445-463', 'CIAGSHAVNGVARIHSEIL']
['447-463', 'AGSHAVNGVARIHSEIL']
['455-468', 'VARIHSEILKKTIF']
['464-471', 'KKTIFKDF']
['472-491', 'YELEPHKFQNKTNGITPRRW']
['495-502', 'CNPGLAEI']
Uptake (Da)
Time (ms)
0
20
40
60
80
100
$10^3$
$10^5$

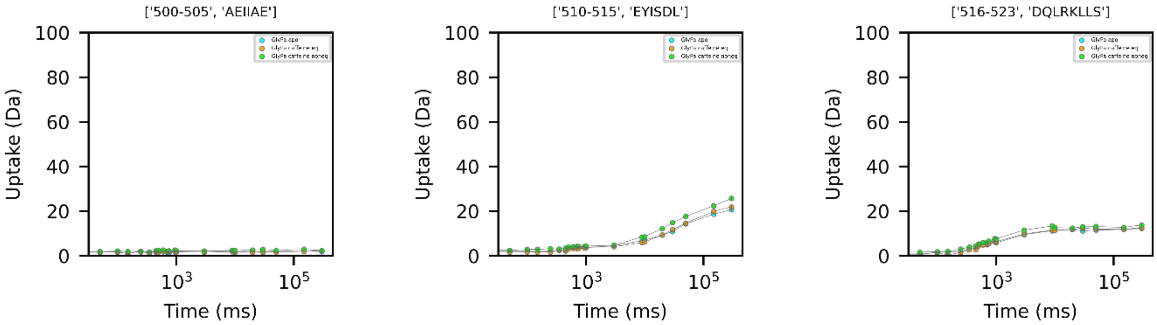
['500-505', 'AEIIAE']
['510-515', 'EYISDL']
['516-523', 'DQLRKLLS']
Uptake (Da)
Time (ms)

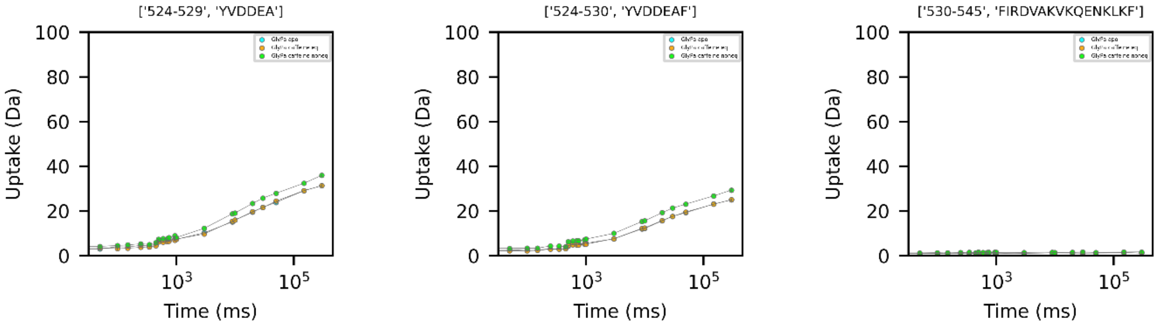
['524-529', 'YVDDEA']
['524-530', 'YVDDEAF']
['530-545', 'FIRDVAKVKQENKLKF']
Uptake (Da)
Time (ms)

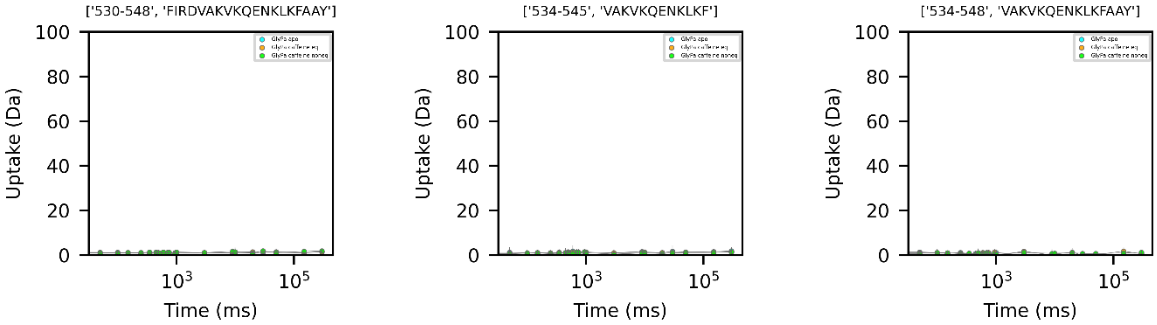
['530-548', 'FIRDVAKVKQENKLKFAAY']
['534-545', 'VAKVKQENKLKF']
['534-548', 'VAKVKQENKLKFAAY']
Uptake (Da)
Time (ms)

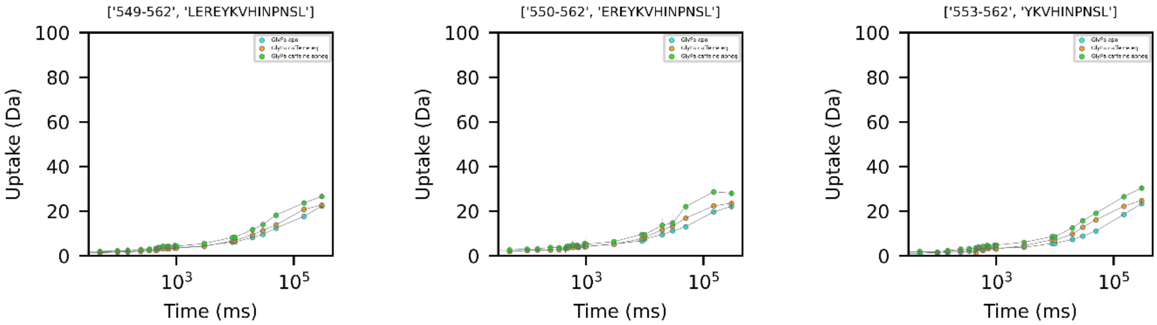
['549-562', 'LEREYKVHINPNSL']
['550-562', 'EREYKVHINPNSL']
['553-562', 'YKVHINPNSL']
Uptake (Da)
Time (ms)

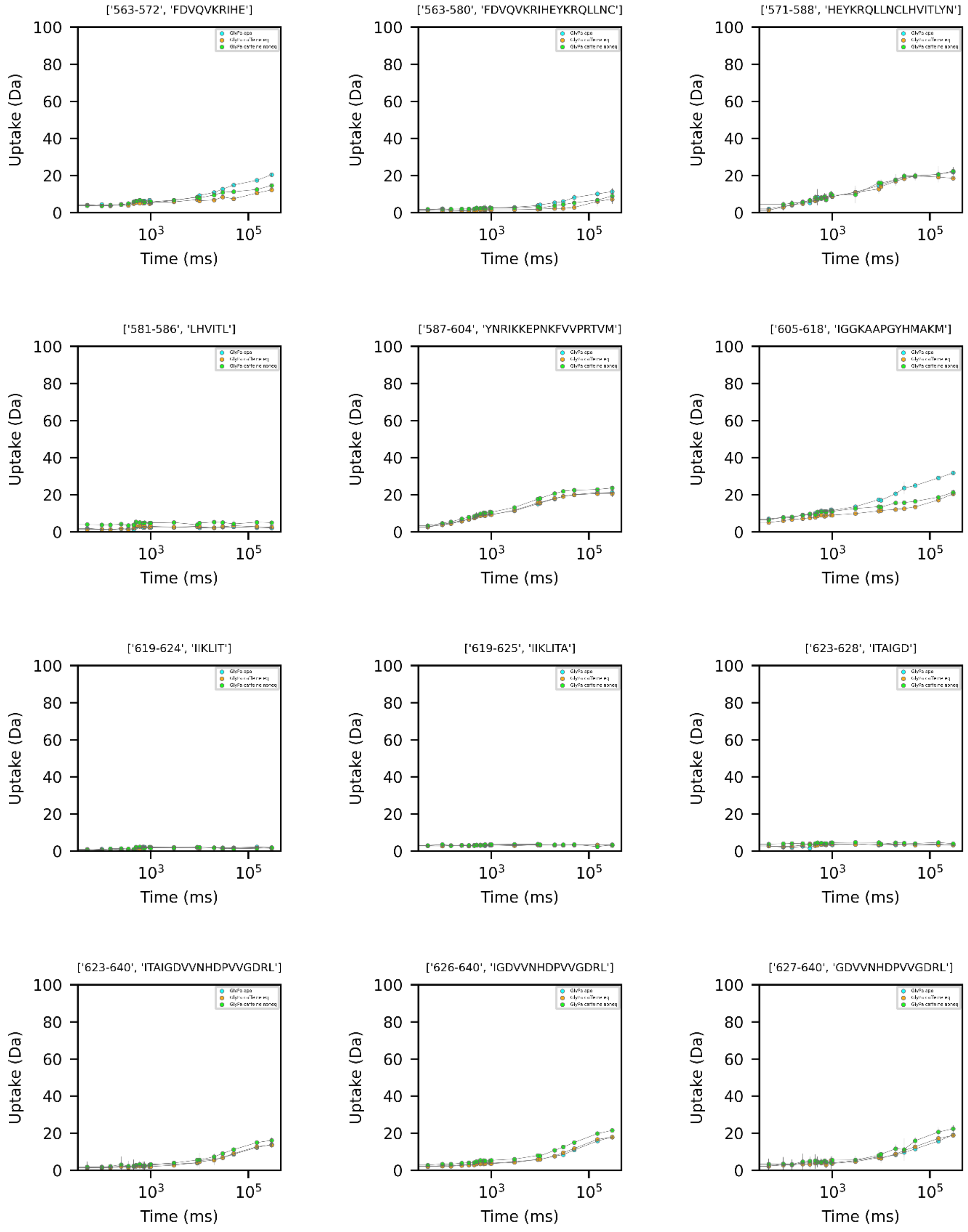
['563-572', 'FDVQVKRIHE']
['563-580', 'FDVQVKRIHEYKRQLLNC']
['571-588', 'HEYKRQLLNCLHVITLYN']
['581-586', 'LHVITL']
['587-604', 'YNRIKKEPNKFVVPRTVM']
['605-618', 'IGGKAAPGYHMAKM']
['619-624', 'IIKLIT']
['619-625', 'IIKLITA']
['623-628', 'ITAIGD']
['623-640', 'ITAIGDVVNHDPVVGDRL']
['626-640', 'IGDVVNHDPVVGDRL']
['627-640', 'GDVVNHDPVVGDRL']
Uptake (Da)
Time (ms)

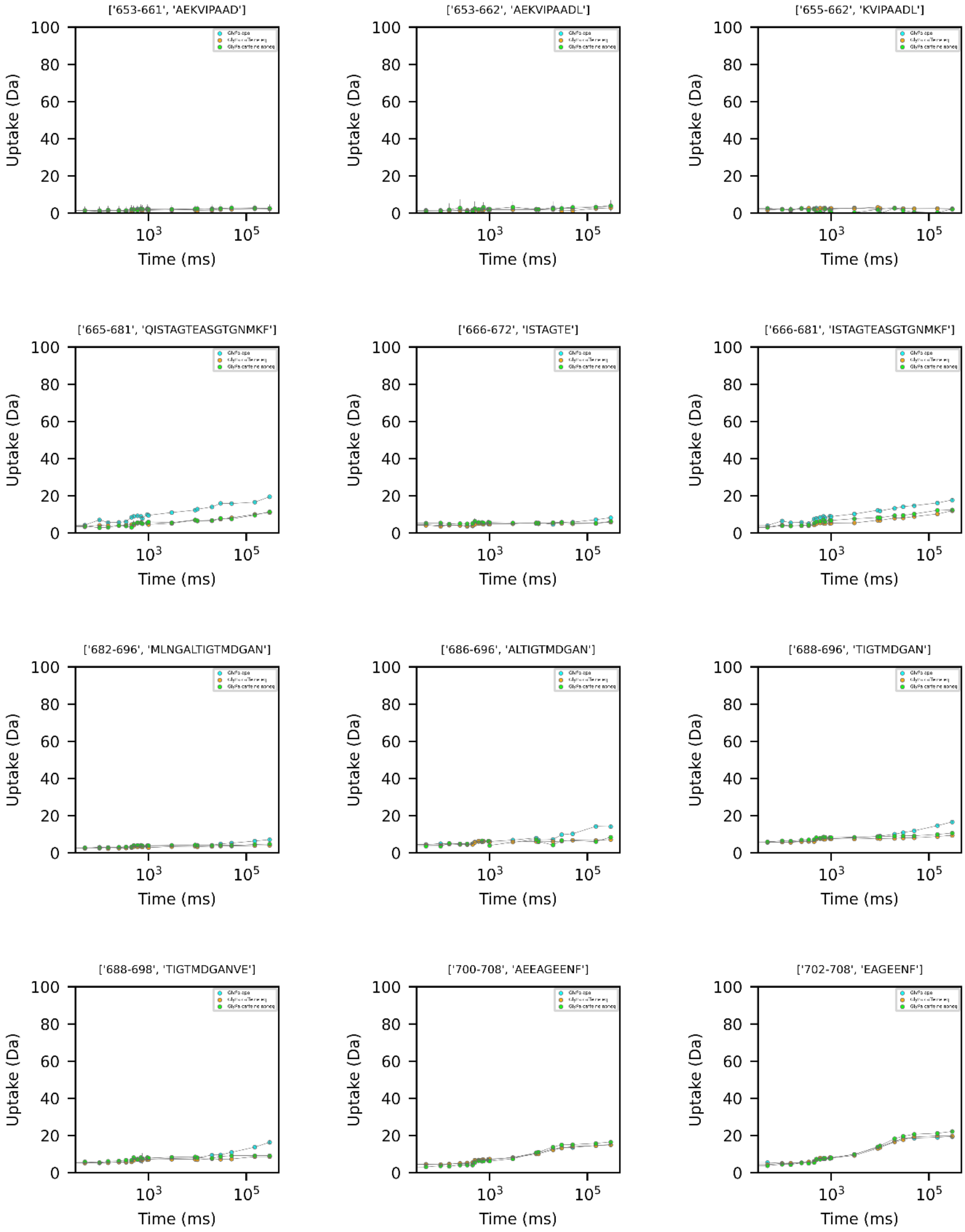
['653-661', 'AEKVIPAAD']
['653-662', 'AEKVIPAADL']
['655-662', 'KVIPAADL']
['665-681', 'QISTAGTEASGTGNMKF']
['666-672', 'ISTAGTE']
['666-681', 'ISTAGTEASGTGNMKF']
['682-696', 'MLNGALTIGTMDGAN']
['686-696', 'ALTIGTMDGAN']
['688-696', 'TIGTMDGAN']
['688-698', 'TIGTMDGANVE']
['700-708', 'AEEAGEENF']
['702-708', 'EAGEENF']
Uptake (Da)
Time (ms)

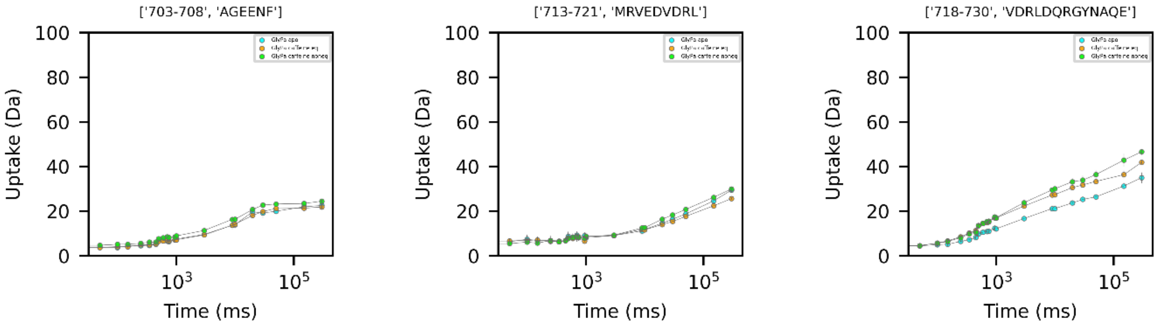
['703-708', 'AGEENF']
['713-721', 'MRVEDVDRL']
['718-730', 'VDRLDQRGYNAQE']
Uptake (Da)
Time (ms)

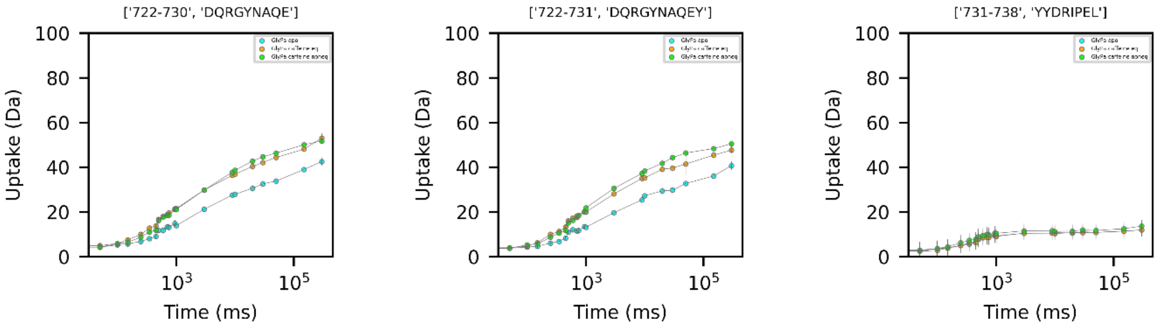
['722-730', 'DQRGYNAQE']
['722-731', 'DQRGYNAQEY']
['731-738', 'YYDRIPEL']
Uptake (Da)
Time (ms)

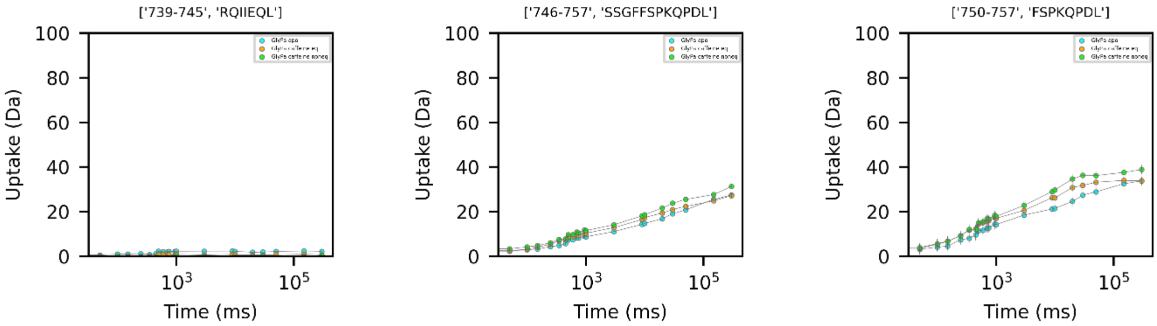
['739-745', 'RQIIEQL']
['746-757', 'SSGFFSPKQPDL']
['750-757', 'FSPKQPDL']
Uptake (Da)
Time (ms)

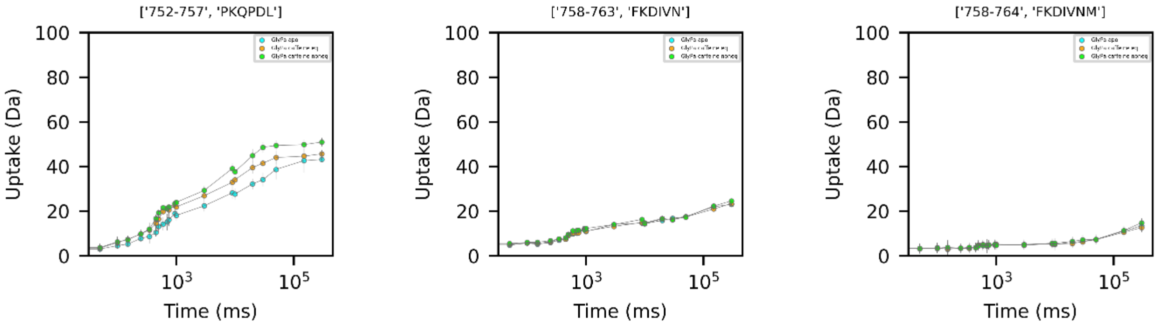
['752-757', 'PKQPDL']
['758-763', 'FKDIVN']
['758-764', 'FKDIVNM']
Uptake (Da)
Time (ms)

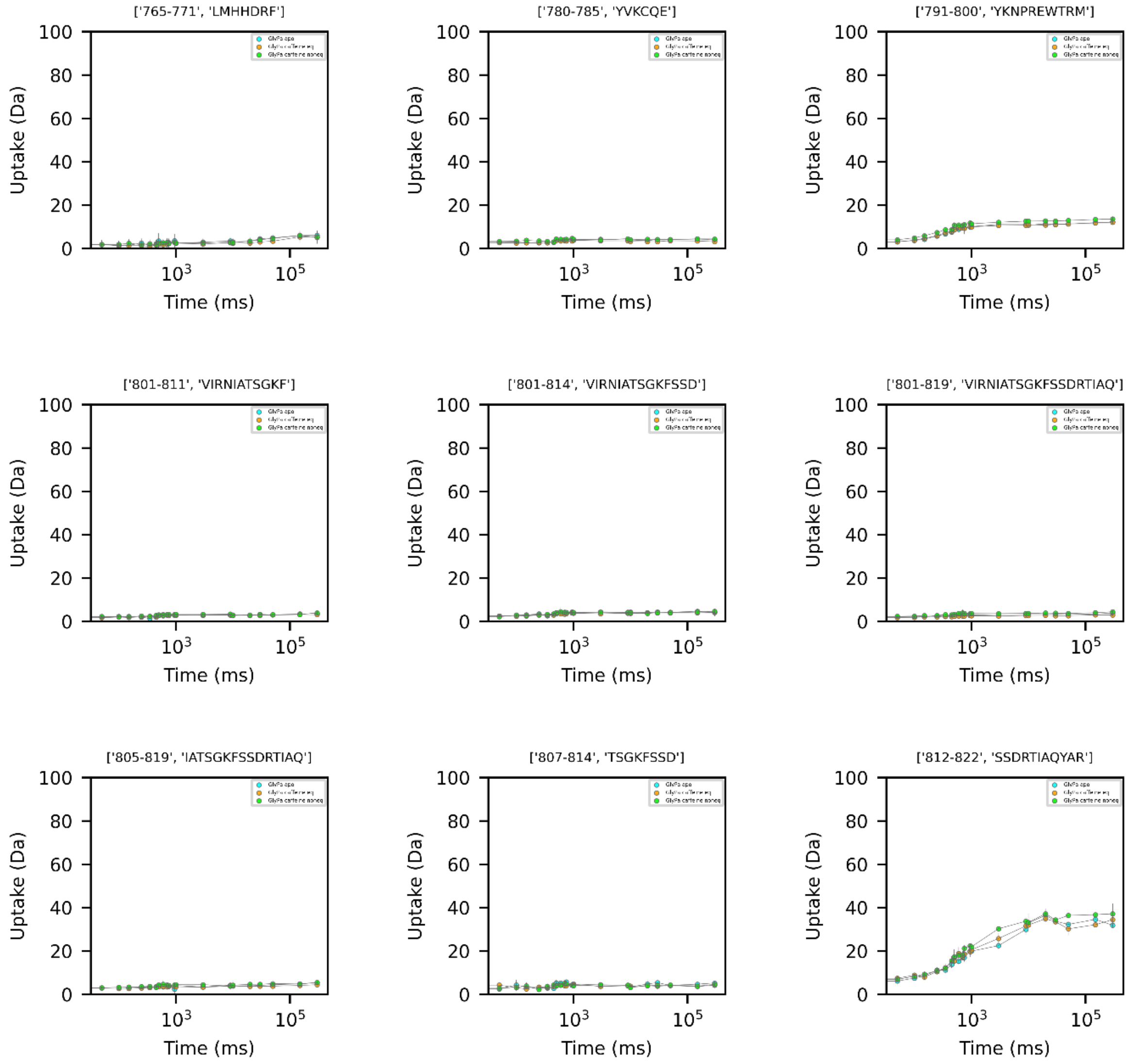
['765-771', 'LMHHDRF']
['780-785', 'YVKCQE']
['791-800', 'YKNPREWTRM']
['801-811', 'VIRNIATSGKF']
['801-814', 'VIRNIATSGKFSSD']
['801-819', 'VIRNIATSGKFSSDRTIAQ']
['805-819', 'IATSGKFSSDRTIAQ']
['807-814', 'TSGKFSSD']
['812-822', 'SSDRTIAQYAR']
Uptake (Da)
Time (ms)
0
20
40
60
80
100
$10^3$
$10^5$

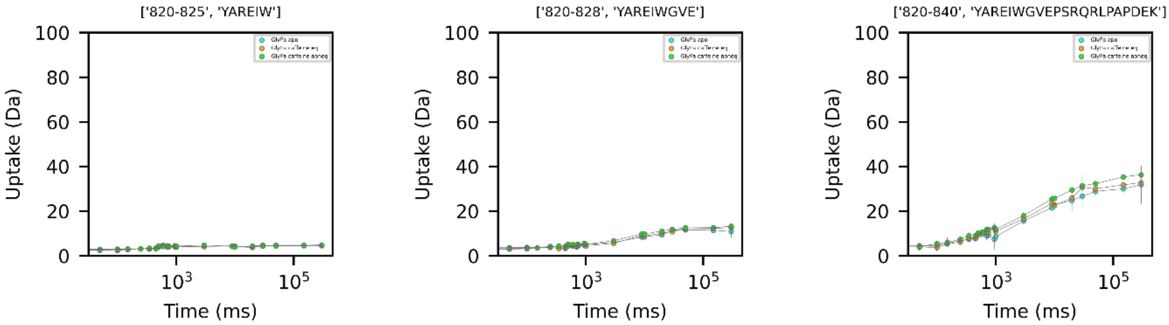
['820-825', 'YAREIW']
['820-828', 'YAREIWGVE']
['820-840', 'YAREIWGVEPSRQRLPAPDEK']
Uptake (Da)
Time (ms)
100
80
60
40
20
0
$10^3$
$10^5$

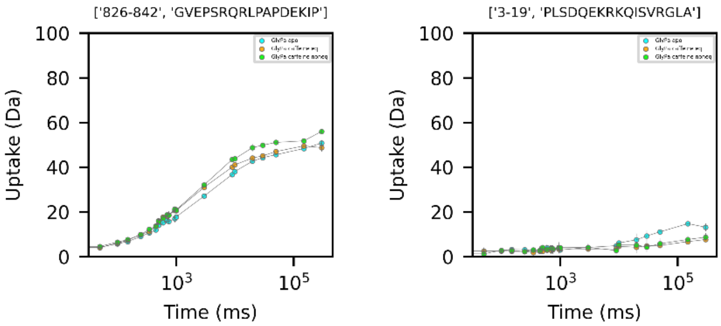
['826-842', 'GVEPSRQRLPAPDEKIP']
['3-19', 'PLSDQEKRKQISVRGLA']
Uptake (Da)
Time (ms)
100
80
60
40
20
0
$10^3$
$10^5$

**Table S2. Local hydrogen exchange rate ($k_{obs}$) of GlyPb peptides.** Exchange rates were calculated by fitting a stretched exponential model to HDX-MS data with 31 time points for 219 peptide segments, for three conditions: (i) inactive GlyPb (apo), (ii) fully activated GlyPb (denoted eq) equilibrated for one hour with 25 mM AMP, 25 mM AS and (iii) GlyPb activated at non-equilibrium with 25 mM AMP, 25 mM AS (denoted by non-eq).

| Sequence | Start | End | q | $k_{obs}$ apo ($s^{-1}$) | $k_{obs}$ eq ($s^{-1}$) | $k_{obs}$ non-eq ($s^{-1}$) |
|---|---|---|---|---|---|---|
| VENVTE | 21 | 26 | 5 | 3.60E-02 | 5.95E-02 | 7.47E-02 |
| NVTELKKNFNRHLHF | 23 | 37 | 14 | 1.60E-04 | 2.53E-04 | 3.84E-05 |
| LKKNFNRHLHFTL | 27 | 39 | 12 | 1.01E-06 | 7.87E-07 | 1.03E-06 |
| TLVKDRNVAT | 38 | 47 | 9 | 6.31E-05 | 6.46E-06 | 1.15E-05 |
| TLVKDRNVATPRD | 38 | 50 | 11 | 2.44E-04 | 1.32E-04 | 3.89E-04 |
| TLVKDRNVATPRDYY | 38 | 52 | 13 | 1.19E-04 | 1.70E-05 | 1.41E-04 |
| VKDRNVATPRD | 40 | 50 | 9 | 4.73E-04 | 1.14E-04 | 7.35E-04 |
| VKDRNVATPRDYY | 40 | 52 | 11 | 1.71E-04 | 7.90E-05 | 3.41E-05 |
| DYYFALAHTVR | 50 | 60 | 10 | 2.39E-07 | 4.70E-06 | 2.40E-07 |
| YYFALA | 51 | 56 | 5 | 8.03E-07 | 6.28E-07 | 7.85E-07 |
| FALAHTVRDHLVGRW | 53 | 67 | 14 | 2.57E-05 | 1.41E-06 | 4.88E-06 |
| ALAHTVRDHLVGRW | 54 | 67 | 13 | 3.20E-06 | 1.68E-06 | 5.57E-05 |
| IRTQQHY | 68 | 74 | 6 | 3.01E-06 | 3.75E-07 | 3.83E-07 |
| IRTQQHYYEKDPKRIYY | 68 | 84 | 15 | 3.92E-04 | 3.77E-05 | 3.93E-05 |
| YEKDPKRIYY | 75 | 84 | 8 | 1.92E-04 | 1.50E-04 | 5.36E-04 |
| PKRIYY | 79 | 84 | 4 | 1.25E-03 | 1.99E-03 | 9.62E-04 |
| EFYMGRTLQNT | 88 | 98 | 10 | 2.05E-04 | 1.83E-04 | 2.01E-04 |
| EFYMGRTLQNTM | 88 | 99 | 11 | 3.66E-04 | 3.32E-04 | 3.95E-04 |
| FYMGRTLQNTM | 89 | 99 | 10 | 9.23E-05 | 8.31E-05 | 1.57E-04 |
| YMGRTLQNT | 90 | 98 | 8 | 7.73E-05 | 3.77E-06 | 9.11E-06 |
| YMGRTLQNTM | 90 | 99 | 9 | 4.13E-05 | 2.29E-06 | 8.22E-06 |
| ALENACD | 103 | 109 | 6 | 3.04E-03 | 2.17E-03 | 6.53E-04 |
| ALENACDE | 103 | 110 | 7 | 8.64E-04 | 2.04E-03 | 3.21E-03 |
| EATYQLGLD | 110 | 118 | 8 | 5.49E-04 | 5.78E-04 | 4.20E-04 |
| ATYQLGL | 111 | 117 | 6 | 4.13E-05 | 3.54E-06 | 5.80E-05 |
| ATYQLGLD | 111 | 118 | 7 | 3.84E-04 | 4.91E-04 | 1.27E-03 |
| YQLGLD | 113 | 118 | 5 | 7.31E-04 | 1.49E-04 | 1.07E-04 |
| EEIEEDAGLGNGGLGRL | 123 | 139 | 16 | 3.06E-04 | 5.71E-05 | 1.34E-04 |
| IEEDAGLGNGGLGRL | 125 | 139 | 14 | 3.87E-04 | 1.08E-04 | 1.56E-04 |
| IEEDAGLGNGGLGRLAA | 125 | 141 | 16 | 3.74E-04 | 4.15E-05 | 6.02E-05 |
| IEEDAGLGNGGLGRLAAC | 125 | 142 | 17 | 8.63E-05 | 3.13E-05 | 4.97E-05 |
| DAGLGNGGLGRL | 128 | 139 | 11 | 4.26E-04 | 1.57E-04 | 4.77E-04 |
| DAGLGNGGLGRLAA | 128 | 141 | 13 | 2.98E-04 | 7.79E-05 | 9.48E-05 |
| DAGLGNGGLGRLAAC | 128 | 142 | 14 | 2.46E-04 | 8.08E-05 | 4.69E-04 |
| LDSMAT | 144 | 149 | 5 | 0.00E+00 | 0.00E+00 | 0.00E+00 |
| LDSMATLGL | 144 | 152 | 8 | 0.00E+00 | 0.00E+00 | 0.00E+00 |
| MATLGL | 147 | 152 | 5 | 0.00E+00 | 0.00E+00 | 0.00E+00 |
| MATLGLAA | 147 | 154 | 7 | 1.47E-06 | 1.16E-06 | 2.72E-06 |
| ATLGLAA | 148 | 154 | 6 | 1.73E-04 | 5.09E-05 | 2.31E-04 |
| AAYGYG | 153 | 158 | 5 | 0.00E+00 | 0.00E+00 | 0.00E+00 |
| AAYGYGIRYEF | 153 | 163 | 10 | 1.81E-04 | 0.00E+00 | 5.16E-05 |
| GYGIRYEF | 156 | 163 | 7 | 5.99E-07 | 2.22E-11 | 1.09E-04 |
| IRYEFGIF | 159 | 166 | 7 | 2.24E-05 | 3.54E-04 | 1.64E-04 |
| GIFNQKICGGWQME | 164 | 177 | 13 | 9.00E-07 | 7.95E-07 | 7.36E-07 |
| NQKICGGWQM | 167 | 176 | 9 | 1.69E-06 | 1.11E-06 | 1.49E-06 |
| QMEEADD | 175 | 181 | 6 | 8.51E-05 | 1.90E-04 | 1.91E-04 |
| MEEADD | 176 | 181 | 5 | 5.74E-04 | 1.21E-03 | 1.36E-03 |
| WLRYGNPWEKARPEF | 182 | 196 | 12 | 2.51E-06 | 5.08E-06 | 3.40E-06 |
| WLRYGNPWEKARPEFTL | 182 | 198 | 14 | 1.45E-06 | 1.14E-05 | 7.93E-06 |
| WLRYGNPWEKARPEFTLPVHF | 182 | 202 | 17 | 2.22E-11 | 0.00E+00 | 0.00E+00 |
| LRYGNPWEKARPEFTL | 183 | 198 | 13 | 5.01E-06 | 9.48E-06 | 8.19E-06 |
| LRYGNPWEKARPEFTLPVHF | 183 | 202 | 16 | 2.55E-06 | 4.26E-06 | 3.66E-06 |
| YGRVEHTSQGAKWVDT | 203 | 218 | 15 | 1.35E-03 | 1.23E-03 | 1.08E-03 |
| YGRVEHTSQGAKWVDTQ | 203 | 219 | 16 | 1.71E-03 | 5.90E-04 | 1.26E-03 |
| QVVLAM | 219 | 224 | 5 | 2.22E-11 | 9.59E-06 | 9.33E-06 |
| QVVLAMPYDTPVPGYRN | 219 | 235 | 13 | 8.37E-07 | 4.57E-06 | 8.39E-07 |
| QVVLAMPYDTPVPGYRNNVVNT | 219 | 240 | 18 | 1.69E-06 | 7.95E-07 | 1.60E-06 |
| VVLAMPYDTPVPGYRNNVVNT | 220 | 240 | 17 | 1.26E-06 | 5.50E-07 | 1.33E-06 |
| LAMPYDTPVPGYRN | 222 | 235 | 10 | 8.98E-07 | 7.64E-07 | 9.35E-07 |
| LAMPYDTPVPGYRNNVVNT | 222 | 240 | 15 | 3.11E-06 | 5.92E-06 | 2.25E-06 |
| AMPYDTPVPGYRNNVVNT | 223 | 240 | 14 | 9.30E-07 | 5.59E-07 | 1.35E-06 |
| PYDTPVPGYRNNVVNT | 225 | 240 | 12 | 1.26E-06 | 1.25E-06 | 1.39E-06 |
| PGYRNNVVNTMRLWS | 231 | 245 | 13 | 6.97E-07 | 6.78E-07 | 9.88E-07 |
| WSAKAPNDF | 244 | 252 | 7 | 1.13E-04 | 2.78E-05 | 4.03E-05 |
| WSAKAPNDFNLKD | 244 | 256 | 11 | 2.48E-03 | 2.39E-03 | 1.67E-03 |
| AKAPNDF | 246 | 252 | 5 | 2.86E-04 | 4.50E-04 | 8.71E-04 |
| NLKDFNVGGYIQ | 253 | 264 | 11 | 1.17E-02 | 1.62E-02 | 1.57E-02 |
| FNVGGYIQ | 257 | 264 | 7 | 3.09E-04 | 1.37E-03 | 1.51E-03 |
| FNVGGYIQA | 257 | 265 | 8 | 8.44E-06 | 8.71E-03 | 6.76E-03 |
| YIQAVL | 262 | 267 | 5 | 4.33E-05 | 1.39E-03 | 2.30E-03 |
| VLDRNLAEN | 266 | 274 | 8 | 3.72E-04 | 8.95E-04 | 1.91E-03 |
| VLDRNLAENI | 266 | 275 | 9 | 6.55E-04 | 1.19E-03 | 1.58E-03 |

| | | | | | | |
|---|---|---|---|---|---|---|
| VLDRNLAENIS | 266 | 276 | 10 | 2.08E-05 | 8.86E-04 | 1.61E-03 |
| DRNLAEN | 268 | 274 | 6 | 4.99E-04 | 1.01E-03 | 2.52E-03 |
| DRNLAENIS | 268 | 276 | 8 | 5.77E-04 | 9.40E-04 | 1.88E-03 |
| DRNLAENISRVLYPNDNF | 268 | 285 | 16 | 9.31E-04 | 6.78E-04 | 1.79E-03 |
| NISRVLYPNDNF | 274 | 285 | 10 | 1.74E-03 | 1.63E-03 | 1.08E-03 |
| ISRVLYPNDNF | 275 | 285 | 9 | 1.41E-03 | 1.04E-03 | 1.03E-03 |
| ISRVLYPNDNFF | 275 | 286 | 10 | 2.79E-03 | 1.39E-03 | 1.24E-03 |
| SRVLYPNDNF | 276 | 285 | 8 | 3.89E-03 | 4.85E-03 | 2.59E-03 |
| RVLYPNDNF | 277 | 285 | 7 | 4.81E-03 | 3.38E-03 | 2.48E-03 |
| FFEGKE | 285 | 290 | 5 | 2.10E-03 | 3.40E-03 | 1.07E-03 |
| LRLKQEY | 291 | 297 | 6 | 3.26E-07 | 2.95E-07 | 2.67E-07 |
| FVVAAT | 298 | 303 | 5 | 0.00E+00 | 0.00E+00 | 0.00E+00 |
| VVAATL | 299 | 304 | 5 | 0.00E+00 | 0.00E+00 | 0.00E+00 |
| DAFPDKVAIQL | 327 | 337 | 9 | 6.39E-07 | 9.15E-07 | 2.38E-07 |
| FPDKVAIQL | 329 | 337 | 7 | 1.52E-06 | 4.99E-06 | 3.14E-07 |
| PDKVAIQL | 330 | 337 | 6 | 0.00E+00 | 0.00E+00 | 0.00E+00 |
| DKVAIQL | 331 | 337 | 6 | 2.22E-11 | 0.00E+00 | 0.00E+00 |
| NDTHPSL | 338 | 344 | 5 | 8.34E-07 | 1.44E-06 | 6.34E-07 |
| NDTHPSLAIPELM | 338 | 350 | 10 | 0.00E+00 | 0.00E+00 | 0.00E+00 |
| MRVLVD | 350 | 355 | 5 | 0.00E+00 | 0.00E+00 | 0.00E+00 |
| MRVLVDL | 350 | 356 | 6 | 0.00E+00 | 0.00E+00 | 0.00E+00 |
| RVLVDL | 351 | 356 | 5 | 0.00E+00 | 0.00E+00 | 0.00E+00 |
| LERLDWDKA | 356 | 364 | 8 | 5.35E-05 | 5.16E-05 | 4.96E-05 |
| LERLDWDKAWE | 356 | 366 | 10 | 7.81E-06 | 6.87E-06 | 9.89E-06 |
| ERLDWDKA | 357 | 364 | 7 | 4.52E-05 | 4.14E-05 | 4.30E-05 |
| ERLDWDKAWE | 357 | 366 | 9 | 2.93E-05 | 2.08E-05 | 1.44E-05 |
| WEVTVKTCA | 365 | 373 | 8 | 1.01E-04 | 2.79E-05 | 2.70E-04 |
| TVKTCA | 368 | 373 | 5 | 0.00E+00 | 3.75E-06 | 0.00E+00 |
| CAYTNHTVLPEAL | 372 | 384 | 11 | 6.91E-04 | 1.88E-03 | 1.20E-03 |
| YTNHTVL | 374 | 380 | 6 | 1.40E-04 | 1.84E-04 | 1.32E-04 |
| YTNHTVLPEAL | 374 | 384 | 9 | 2.06E-03 | 2.07E-03 | 1.97E-03 |
| TNHTVL | 375 | 380 | 5 | 7.42E-04 | 8.58E-04 | 8.99E-04 |
| ERWPVHL | 385 | 391 | 5 | 1.28E-05 | 1.30E-05 | 1.59E-05 |
| ERWPVHLLE | 385 | 393 | 7 | 1.10E-05 | 1.31E-05 | 5.37E-05 |
| LETLLPRHL | 392 | 400 | 7 | 3.36E-05 | 1.33E-05 | 3.98E-05 |
| LETLLPRHLQI | 392 | 402 | 9 | 4.90E-06 | 2.04E-06 | 3.15E-06 |
| LETLLPRHLQIIY | 392 | 404 | 11 | 6.12E-05 | 4.36E-05 | 4.46E-05 |
| LETLLPRHLQIIYE | 392 | 405 | 12 | 1.97E-05 | 0.00E+00 | 0.00E+00 |
| ETLLPRHLQIIY | 393 | 404 | 10 | 2.81E-05 | 1.77E-05 | 2.22E-11 |
| TLLPRHLQIIY | 394 | 404 | 9 | 3.55E-06 | 1.20E-06 | 0.00E+00 |
| TLLPRHLQIIYE | 394 | 405 | 10 | 0.00E+00 | 0.00E+00 | 0.00E+00 |
| LPRHLQIIYE | 396 | 405 | 8 | 1.27E-06 | 1.48E-06 | 1.91E-07 |
| PRHLQIIYE | 397 | 405 | 7 | 3.82E-06 | 3.61E-06 | 3.61E-06 |
| IYEINQRFLNRVAAAF | 403 | 418 | 15 | 0.00E+00 | 0.00E+00 | 0.00E+00 |
| EINQRFLNRVAAAF | 405 | 418 | 13 | 6.15E-07 | 5.07E-06 | 3.42E-06 |
| EINQRFLNRVAAAFPGDVDRL | 405 | 425 | 19 | 1.68E-06 | 1.88E-06 | 1.57E-06 |
| INQRFLNRVAAAF | 406 | 418 | 12 | 4.89E-06 | 2.87E-06 | 5.72E-07 |
| INQRFLNRVAAAFPGDVDRL | 406 | 425 | 18 | 2.03E-06 | 1.74E-06 | 1.66E-06 |
| LNRVAAAFPGDVDRL | 411 | 425 | 13 | 1.88E-06 | 3.33E-06 | 2.17E-06 |
| PGDVDRL | 419 | 425 | 5 | 5.75E-04 | 5.73E-04 | 2.99E-04 |
| VEEGAVKRINM | 431 | 441 | 10 | 7.80E-05 | 1.17E-04 | 8.47E-05 |
| EEGAVKRINM | 432 | 441 | 9 | 1.49E-05 | 6.76E-06 | 3.24E-06 |
| CIAGSHAVNG | 445 | 454 | 9 | 8.01E-07 | 8.19E-07 | 8.33E-07 |
| CIAGSHAVNGVARIHS | 445 | 460 | 15 | 8.78E-07 | 3.27E-06 | 1.30E-06 |
| CIAGSHAVNGVARIHSE | 445 | 461 | 16 | 6.52E-07 | 3.91E-06 | 7.66E-07 |
| CIAGSHAVNGVARIHSEILKKTIF | 445 | 468 | 23 | 3.56E-05 | 6.86E-05 | 3.56E-04 |
| AGSHAVNGVARIHSE | 447 | 461 | 14 | 6.51E-07 | 4.54E-06 | 8.96E-07 |
| AGSHAVNGVARIHSEIL | 447 | 463 | 16 | 6.76E-07 | 1.52E-06 | 5.80E-07 |
| AGSHAVNGVARIHSEILKKTIF | 447 | 468 | 21 | 5.31E-05 | 1.08E-04 | 2.53E-04 |
| KKTIFKDF | 464 | 471 | 7 | 4.58E-04 | 7.73E-04 | 7.09E-04 |
| YELEPHKFQNKTNGITPRRW | 472 | 491 | 17 | 1.13E-06 | 1.10E-06 | 4.88E-07 |
| YELEPHKFQNKTNGITPRRWLVL | 472 | 494 | 20 | 6.80E-07 | 5.53E-07 | 5.20E-07 |
| LVLCNPGL | 492 | 499 | 6 | 6.41E-04 | 5.01E-04 | 5.67E-04 |
| CNPGLAE | 495 | 501 | 5 | 4.63E-05 | 1.15E-04 | 6.34E-04 |
| CNPGLAEI | 495 | 502 | 6 | 3.03E-04 | 4.13E-04 | 4.33E-04 |
| GLAEII | 498 | 503 | 5 | 0.00E+00 | 0.00E+00 | 0.00E+00 |
| EIIAERIGEE | 501 | 510 | 9 | 6.09E-04 | 5.51E-04 | 1.55E-04 |
| IAERIGE | 503 | 509 | 6 | 2.90E-06 | 4.03E-06 | 1.63E-05 |
| IAERIGEE | 503 | 510 | 7 | 2.75E-04 | 4.68E-05 | 6.81E-04 |
| AERIGEE | 504 | 510 | 6 | 1.12E-04 | 4.19E-04 | 7.79E-04 |
| YISDLDQ | 511 | 517 | 6 | 6.95E-04 | 5.49E-04 | 1.90E-03 |
| DQLRKLLS | 516 | 523 | 7 | 2.49E-06 | 1.05E-05 | 2.67E-06 |
| YVDDEA | 524 | 529 | 5 | 3.66E-03 | 5.61E-03 | 9.17E-04 |
| YVDDEAF | 524 | 530 | 6 | 2.33E-03 | 2.41E-03 | 2.66E-03 |
| FIRDVAKVKQENKLKFAAY | 530 | 548 | 18 | 2.02E-06 | 1.92E-06 | 5.89E-06 |
| IRDVAKVKQENKLKFAAY | 531 | 548 | 17 | 2.83E-07 | 1.59E-06 | 4.46E-06 |
| YLEREYKVHINPNSL | 548 | 562 | 13 | 3.21E-04 | 2.95E-04 | 7.74E-04 |
| LEREYKVHINPNSL | 549 | 562 | 12 | 2.77E-04 | 2.67E-04 | 3.24E-04 |
| LEREYKVHINPNSLF | 549 | 563 | 13 | 1.69E-04 | 1.41E-04 | 1.87E-04 |
| EREYKVHINPNSL | 550 | 562 | 11 | 3.57E-04 | 3.31E-04 | 3.88E-04 |
| YKVHINPNSL | 553 | 562 | 8 | 1.81E-04 | 1.55E-04 | 2.20E-04 |

| | | | | | | |
|---|---|---|---|---|---|---|
| FDVQVKRIHE | 563 | 572 | 9 | 1.89E-06 | 5.14E-06 | 3.13E-06 |
| FDVQVKRIHEYKRQLLNC | 563 | 580 | 17 | 1.30E-04 | 5.41E-05 | 1.25E-04 |
| DVQVKRIHEYKRQLLNC | 564 | 580 | 16 | 3.41E-05 | 2.68E-05 | 7.76E-05 |
| LHVITL | 581 | 586 | 5 | 0.00E+00 | 0.00E+00 | 0.00E+00 |
| YNRIKKEPNKFVVPRTVM | 587 | 604 | 15 | 1.88E-04 | 2.67E-04 | 2.49E-04 |
| IGGKAAPGYHM | 605 | 615 | 9 | 7.43E-05 | 3.85E-05 | 1.70E-05 |
| IGGKAAPGYHMAKM | 605 | 618 | 12 | 8.62E-05 | 4.72E-05 | 2.76E-05 |
| GGKAAPGYHMAKM | 606 | 618 | 11 | 1.15E-04 | 3.07E-05 | 3.59E-05 |
| IIKLIT | 619 | 624 | 5 | 0.00E+00 | 0.00E+00 | 0.00E+00 |
| IIKLITA | 619 | 625 | 6 | 4.10E-06 | 3.31E-06 | 1.74E-07 |
| IIKLITAIGDVVNHDPVVGDRL | 619 | 640 | 20 | 2.70E-04 | 9.62E-05 | 5.04E-04 |
| ITAIGD | 623 | 628 | 5 | 7.07E-07 | 7.28E-07 | 2.66E-07 |
| TAIGDVVNHDPVVGDRL | 624 | 640 | 15 | 8.13E-05 | 6.77E-05 | 2.74E-04 |
| AIGDVVNHDPVVGDRL | 625 | 640 | 14 | 2.66E-05 | 2.37E-05 | 7.50E-05 |
| IGDVVNHDPVVGDRL | 626 | 640 | 13 | 2.98E-05 | 2.84E-05 | 7.76E-05 |
| LENYRVSL | 645 | 652 | 7 | 2.66E-05 | 3.30E-06 | 3.25E-05 |
| YRVSLAE | 648 | 654 | 6 | 8.08E-05 | 3.71E-05 | 1.90E-05 |
| AEKVIPAAD | 653 | 661 | 7 | 2.51E-06 | 2.64E-06 | 8.61E-06 |
| AEKVIPAADL | 653 | 662 | 8 | 5.02E-07 | 9.88E-07 | 4.52E-07 |
| SEQISTAGTEASGTGNMKF | 663 | 681 | 18 | 2.14E-06 | 2.93E-06 | 2.05E-06 |
| ISTAGTEASGTGNMKF | 666 | 681 | 15 | 2.67E-06 | 4.99E-06 | 2.32E-06 |
| MLNGALT | 682 | 688 | 6 | 0.00E+00 | 0.00E+00 | 0.00E+00 |
| MLNGALTIGTM | 682 | 692 | 10 | 2.71E-06 | 2.76E-06 | 1.26E-06 |
| MLNGALTIGTMDGAN | 682 | 696 | 14 | 1.02E-06 | 1.07E-06 | 9.82E-07 |
| MLNGALTIGTMDGANVE | 682 | 698 | 16 | 1.12E-06 | 7.82E-07 | 1.09E-06 |
| LTIGTMDGAN | 687 | 696 | 9 | 1.20E-06 | 1.09E-06 | 9.60E-07 |
| TIGTMDGAN | 688 | 696 | 8 | 1.11E-06 | 7.16E-07 | 6.75E-07 |
| TIGTMDGANVE | 688 | 698 | 10 | 1.30E-06 | 5.98E-07 | 9.46E-07 |
| GTMDGAN | 690 | 696 | 6 | 7.44E-07 | 1.59E-06 | 9.82E-07 |
| MAEEAGEENF | 699 | 708 | 9 | 1.83E-06 | 2.56E-06 | 2.87E-06 |
| AEEAGEENF | 700 | 708 | 8 | 4.82E-05 | 4.40E-05 | 4.66E-05 |
| EAGEENF | 702 | 708 | 6 | 3.81E-04 | 1.97E-04 | 5.19E-04 |
| AGEENF | 703 | 708 | 5 | 1.47E-03 | 1.54E-03 | 2.65E-03 |
| FIFGMRV | 709 | 715 | 6 | 6.72E-04 | 6.63E-04 | 8.60E-04 |
| FIFGMRVED | 709 | 717 | 8 | 4.54E-04 | 5.12E-04 | 4.94E-04 |
| RVEDVDRL | 714 | 721 | 7 | 4.97E-05 | 6.69E-05 | 8.21E-05 |
| VDRLDQRGYNAQE | 718 | 730 | 12 | 1.01E-03 | 7.24E-04 | 7.06E-04 |
| VDRLDQRGYNAQEY | 718 | 731 | 13 | 1.00E-03 | 7.99E-04 | 6.70E-04 |
| DQRGYNAQE | 722 | 730 | 8 | 3.43E-03 | 2.33E-03 | 2.32E-03 |
| DQRGYNAQEY | 722 | 731 | 9 | 4.82E-03 | 1.63E-03 | 2.06E-03 |
| YDRIPEL | 732 | 738 | 5 | 6.36E-07 | 1.33E-06 | 1.52E-06 |
| ELRQIIEQLSSGFFSPKQPDLFK | 737 | 759 | 20 | 1.69E-04 | 2.56E-04 | 2.20E-04 |
| RQIIEQL | 739 | 745 | 6 | 2.22E-11 | 0.00E+00 | 0.00E+00 |
| SSGFFSPKQPDL | 746 | 757 | 9 | 3.79E-04 | 2.83E-04 | 4.20E-04 |
| FFSPKQPDL | 749 | 757 | 6 | 8.22E-04 | 6.45E-04 | 8.95E-04 |
| FSPKQPDL | 750 | 757 | 5 | 2.33E-03 | 1.84E-03 | 1.85E-03 |
| SPKQPDL | 751 | 757 | 4 | 3.15E-03 | 1.60E-03 | 2.85E-03 |
| PKQPDL | 752 | 757 | 3 | 1.69E-02 | 1.15E-02 | 8.36E-03 |
| FKDIVNM | 758 | 764 | 6 | 4.22E-05 | 3.79E-05 | 9.91E-05 |
| YVKCQE | 780 | 785 | 5 | 4.13E-06 | 1.36E-06 | 2.96E-07 |
| YVKCQERVSA | 780 | 789 | 9 | 0.00E+00 | 0.00E+00 | 0.00E+00 |
| YVKCQERVSAL | 780 | 790 | 10 | 0.00E+00 | 0.00E+00 | 0.00E+00 |
| LYKNPRE | 790 | 796 | 5 | 1.65E-06 | 1.75E-06 | 7.03E-07 |
| YKNPREWTRM | 791 | 800 | 8 | 9.95E-07 | 1.44E-06 | 1.39E-06 |
| VIRNIATSGKF | 801 | 811 | 10 | 9.75E-07 | 8.48E-07 | 7.37E-07 |
| VIRNIATSGKFSSD | 801 | 814 | 13 | 9.38E-07 | 9.72E-07 | 9.20E-07 |
| VIRNIATSGKFSSDRTIAQ | 801 | 819 | 18 | 1.01E-06 | 8.51E-07 | 9.47E-07 |
| IATSGKFSSD | 805 | 814 | 9 | 8.20E-07 | 6.45E-07 | 6.62E-07 |
| IATSGKFSSDRTIAQ | 805 | 819 | 14 | 8.36E-07 | 5.80E-07 | 7.78E-07 |
| TSGKFSSD | 807 | 814 | 7 | 6.11E-07 | 7.91E-07 | 8.81E-07 |
| SGKFSSD | 808 | 814 | 6 | 9.08E-07 | 8.97E-07 | 9.57E-07 |
| SSDRTIAQ | 812 | 819 | 7 | 9.46E-07 | 8.44E-07 | 7.85E-07 |
| YAREIW | 820 | 825 | 5 | 1.02E-06 | 9.90E-07 | 9.76E-07 |
| YAREIWGVE | 820 | 828 | 8 | 1.13E-04 | 2.86E-06 | 6.60E-05 |
| IWGVEPSRQRLPAPDEKIP | 824 | 842 | 14 | 1.11E-02 | 4.29E-03 | 5.05E-03 |
| GVEPSRQRLPAPDEKIP | 826 | 842 | 12 | 1.51E-02 | 1.71E-02 | 1.91E-02 |

**Table S3. Local hydrogen exchange rate ($k_{obs}$) of GlyPa peptides.** Exchange rates were calculated by fitting a stretched exponential model to HDX-MS data with 20 time points for 171 peptide segments, for three conditions: (i) inactive GlyPa (apo), (ii) fully activated GlyPa (denoted eq) equilibrated for one hour with 32 mM caffeine,(iii) GlyPa activated at non-equilibrium with 32 mM caffeine (denoted by non-eq).

| Sequence | Start | End | q | $k_{obs}$ apo ($s^{-1}$) | $k_{obs}$ eq ($s^{-1}$) | $k_{obs}$ non-eq ($s^{-1}$) |
|---|---|---|---|---|---|---|
| PLSDQEKRKQISVRGLA | 3 | 19 | 15 | 2.19E-09 | 1.21E-09 | 1.12E-09 |
| SVRGLAGVEN | 14 | 23 | 9 | 2.29E-09 | 2.33E-09 | 3.21E-09 |
| LKKNFNRHLHF | 27 | 37 | 10 | 1.02E-09 | 9.45E-10 | 1.17E-09 |
| TLVKDRNVATPRD | 38 | 50 | 11 | 6.28E-09 | 3.02E-08 | 5.30E-08 |
| TLVKDRNVATPRDYY | 38 | 52 | 13 | 4.91E-09 | 9.26E-09 | 2.53E-07 |
| VKDRNVATPRDYY | 40 | 52 | 11 | 1.38E-08 | 1.26E-08 | 2.40E-07 |
| RDYYFALAHTVRD | 49 | 61 | 12 | 3.44E-09 | 3.38E-09 | 4.60E-09 |
| YYFALA | 51 | 56 | 5 | 1.61E-09 | 1.11E-09 | 9.55E-10 |
| FALAHTVRDHL | 53 | 63 | 10 | 8.15E-10 | 3.63E-10 | 6.60E-10 |
| FALAHTVRDHLVG | 53 | 65 | 12 | 1.13E-09 | 8.23E-10 | 1.67E-09 |
| FALAHTVRDHLVGRW | 53 | 67 | 14 | 1.23E-09 | 2.36E-09 | 1.81E-09 |
| HLVGRW | 62 | 67 | 5 | 2.71E-09 | 1.70E-07 | 1.40E-07 |
| IRTQQHY | 68 | 74 | 6 | 5.42E-10 | 9.43E-10 | 7.00E-10 |
| IRTQQHYYEKDPKRIYY | 68 | 84 | 15 | 1.05E-09 | 2.06E-09 | 1.35E-09 |
| YEKDPKRIYY | 75 | 84 | 8 | 7.59E-09 | 8.34E-09 | 3.80E-09 |
| PKRIYY | 79 | 84 | 4 | 2.71E-09 | 2.03E-08 | 2.00E-09 |
| EFYMGRTLQNTM | 88 | 99 | 11 | 6.07E-07 | 6.15E-07 | 8.24E-07 |
| EFYMGRTLQNTMVN | 88 | 101 | 13 | 1.11E-09 | 3.25E-09 | 2.62E-09 |
| FYMGRTLQNT | 89 | 98 | 9 | 2.66E-09 | 1.23E-09 | 3.27E-08 |
| YMGRTLQNTMVN | 90 | 101 | 11 | 5.10E-09 | 2.26E-08 | 6.39E-09 |
| ALENAC | 103 | 108 | 5 | 1.22E-06 | 4.09E-06 | 8.75E-06 |
| ALENACD | 103 | 109 | 6 | 3.61E-07 | 5.89E-07 | 2.32E-06 |
| ALENACDE | 103 | 110 | 7 | 3.19E-07 | 4.37E-07 | 9.50E-07 |
| EATYQLGLD | 110 | 118 | 8 | 4.09E-08 | 3.58E-08 | 5.14E-08 |
| ATYQLGLD | 111 | 118 | 7 | 5.30E-09 | 1.19E-07 | 3.32E-08 |
| ATYQLGLDM | 111 | 119 | 8 | 4.02E-08 | 1.42E-07 | 4.67E-08 |
| YQLGLD | 113 | 118 | 5 | 8.62E-08 | 3.73E-08 | 2.24E-08 |
| YQLGLDM | 113 | 119 | 6 | 4.23E-08 | 6.45E-08 | 3.87E-08 |
| EEIEEDAGLGNGGLGRL | 123 | 139 | 16 | 1.12E-07 | 1.82E-07 | 5.71E-08 |
| IEEDAGLGNGGLGRL | 125 | 139 | 14 | 2.04E-07 | 1.22E-07 | 9.80E-08 |
| IEEDAGLGNGGLGRLAA | 125 | 141 | 16 | 8.96E-08 | 9.84E-08 | 1.50E-08 |
| IEEDAGLGNGGLGRLAAC | 125 | 142 | 17 | 5.40E-08 | 8.41E-08 | 2.86E-08 |
| DAGLGNGGLGRL | 128 | 139 | 11 | 3.98E-07 | 1.69E-07 | 1.03E-07 |
| LDSMAT | 144 | 149 | 5 | 5.25E-10 | 5.83E-10 | 6.26E-10 |
| MATLGL | 147 | 152 | 5 | 4.78E-10 | 8.72E-10 | 2.95E-10 |
| MATLGLAA | 147 | 154 | 7 | 4.31E-10 | 1.66E-09 | 5.53E-10 |
| AAYGYG | 153 | 158 | 5 | 4.27E-10 | 3.83E-10 | 1.57E-10 |
| AAYGYGIRYEF | 153 | 163 | 10 | 7.00E-10 | 9.23E-10 | 1.01E-09 |
| YGYGIRYEF | 155 | 163 | 8 | 4.92E-10 | 6.16E-10 | 5.54E-10 |
| GIFNQKICGGWQ | 164 | 175 | 11 | 3.46E-09 | 0.001128767 | 8.44E-10 |
| NQKICGGWQM | 167 | 176 | 9 | 1.35E-09 | 1.49E-09 | 1.46E-09 |
| NQKICGGWQMEE | 167 | 178 | 11 | 9.78E-10 | 7.05E-10 | 9.35E-10 |
| QMEEADD | 175 | 181 | 6 | 4.71E-07 | 5.82E-08 | 1.82E-07 |
| MEEADD | 176 | 181 | 5 | 3.70E-07 | 5.62E-08 | 2.39E-07 |
| WLRYGNPWEKARPEF | 182 | 196 | 12 | 8.12E-10 | 4.77E-10 | 8.79E-10 |
| WLRYGNPWEKARPEFTL | 182 | 198 | 14 | 8.62E-10 | 2.05E-09 | 3.62E-10 |
| YGRVEHTSQGAKW | 203 | 215 | 12 | 1.86E-07 | 1.61E-07 | 2.94E-07 |
| YGRVEHTSQGAKWVDT | 203 | 218 | 15 | 1.14E-07 | 6.35E-07 | 2.62E-07 |
| YGRVEHTSQGAKWVDTQ | 203 | 219 | 16 | 2.28E-07 | 3.72E-07 | 6.07E-07 |
| QVVLAM | 219 | 224 | 5 | 5.06E-10 | 8.08E-10 | 4.51E-10 |
| AMPYDTPVPGYRN | 223 | 235 | 9 | 1.09E-09 | 1.20E-09 | 3.99E-10 |
| AMPYDTPVPGYRNNVVNT | 223 | 240 | 14 | 1.16E-09 | 1.18E-09 | 1.58E-09 |
| PVPGYRNNVVNT | 229 | 240 | 9 | 7.44E-10 | 6.06E-10 | 9.80E-10 |
| WSAKAPND | 244 | 251 | 6 | 4.41E-09 | 2.21E-09 | 2.63E-09 |
| WSAKAPNDF | 244 | 252 | 7 | 7.48E-09 | 1.99E-09 | 1.50E-09 |
| WSAKAPNDFNLKD | 244 | 256 | 11 | 5.93E-07 | 3.07E-07 | 1.55E-06 |
| NLKDFNVGG | 253 | 261 | 8 | 1.65E-05 | 0.000776095 | 2.71E-05 |
| NLKDFNVGGYIQ | 253 | 264 | 11 | 4.76E-06 | 7.39E-06 | 3.81E-05 |
| FNVGGYIQ | 257 | 264 | 7 | 2.88E-06 | 3.73E-06 | 4.43E-05 |
| FNVGGYIQA | 257 | 265 | 8 | 1.37E-06 | 3.13E-06 | 2.70E-05 |
| VLDRNLAEN | 266 | 274 | 8 | 1.27E-07 | 6.01E-06 | 1.56E-05 |
| DRNLAE | 268 | 273 | 5 | 1.46E-08 | 4.01E-06 | 1.46E-05 |
| DRNLAEN | 268 | 274 | 6 | 1.48E-07 | 4.03E-06 | 1.38E-05 |
| LAENISRVLYPNDNF | 271 | 285 | 13 | 4.07E-08 | 2.79E-07 | 2.68E-07 |
| NISRVLYPNDN | 274 | 284 | 9 | 1.49E-08 | 1.65E-07 | 5.77E-09 |
| NISRVLYPNDNF | 274 | 285 | 10 | 5.02E-08 | 2.33E-07 | 2.92E-08 |
| ISRVLYPNDN | 275 | 284 | 8 | 1.94E-08 | 3.36E-08 | 1.49E-08 |
| ISRVLYPNDNF | 275 | 285 | 9 | 4.84E-08 | 2.16E-07 | 3.82E-08 |
| FFEGKE | 285 | 290 | 5 | 3.27E-07 | 2.46E-07 | 3.27E-07 |
| LRLKQEY | 291 | 297 | 6 | 8.17E-10 | 3.79E-09 | 7.52E-10 |
| IIRRFKSSKFGCRDPVRTNF | 307 | 326 | 18 | 2.72E-09 | 2.17E-09 | 9.20E-09 |
| DAFPDKVAIQL | 327 | 337 | 9 | 8.83E-10 | 5.15E-10 | 8.73E-10 |
| PDKVAIQL | 330 | 337 | 6 | 8.36E-10 | 7.81E-10 | 9.24E-10 |
| NDTHPSL | 338 | 344 | 5 | 1.49E-09 | 5.27E-10 | 5.14E-10 |

| | | | | | | |
|---|---|---|---|---|---|---|
| NDTHPSLAIPEL | 338 | 349 | 9 | 9.22E-10 | 3.14E-10 | 3.24E-10 |
| MRVLVD | 350 | 355 | 5 | 5.08E-10 | 3.71E-10 | 9.15E-10 |
| MRVLVDL | 350 | 356 | 6 | 8.40E-10 | 8.69E-10 | 1.07E-09 |
| RVLVDL | 351 | 356 | 5 | 9.07E-10 | 3.54E-10 | 7.23E-10 |
| ERLDWDKAWE | 357 | 366 | 9 | 3.80E-09 | 2.54E-09 | 3.16E-09 |
| VTVKTCA | 367 | 373 | 6 | 1.93E-10 | 8.78E-10 | 1.06E-09 |
| TVKTCA | 368 | 373 | 5 | 8.75E-10 | 4.33E-10 | 7.94E-10 |
| VKTCAYTNHTVLPEAL | 369 | 384 | 14 | 1.98E-07 | 1.10E-08 | 2.15E-09 |
| CAYTNHTVL | 372 | 380 | 8 | 3.60E-08 | 1.69E-09 | 6.37E-10 |
| CAYTNHTVLPEAL | 372 | 384 | 11 | 7.92E-07 | 1.02E-07 | 4.90E-09 |
| YTNHTVL | 374 | 380 | 6 | 1.43E-07 | 5.25E-09 | 1.12E-09 |
| YTNHTVLPEAL | 374 | 384 | 9 | 2.89E-07 | 1.08E-07 | 2.18E-08 |
| TNHTVL | 375 | 380 | 5 | 5.06E-08 | 8.23E-10 | 7.18E-10 |
| LERWPVHLLE | 384 | 393 | 8 | 1.70E-08 | 1.51E-09 | 2.03E-09 |
| ERWPVHL | 385 | 391 | 5 | 1.36E-09 | 1.52E-09 | 6.05E-09 |
| ERWPVHLLE | 385 | 393 | 7 | 2.34E-09 | 1.41E-09 | 1.90E-09 |
| LETLLPRHLQI | 392 | 402 | 9 | 9.05E-10 | 8.30E-10 | 1.14E-09 |
| LETLLPRHLQIIYE | 392 | 405 | 12 | 6.95E-10 | 1.13E-09 | 6.60E-10 |
| TLLPRHLQIIY | 394 | 404 | 9 | 6.16E-10 | 1.64E-09 | 7.72E-10 |
| TLLPRHLQIIYE | 394 | 405 | 10 | 7.70E-10 | 6.96E-10 | 7.67E-10 |
| EINQRFLNRVAAAF | 405 | 418 | 13 | 7.46E-10 | 7.98E-10 | 8.59E-10 |
| INQRFLNRVAAAF | 406 | 418 | 12 | 8.06E-10 | 8.40E-10 | 8.21E-10 |
| AFPGDVDRLR | 417 | 426 | 8 | 1.52E-08 | 1.37E-08 | 2.44E-08 |
| PGDVDRL | 419 | 425 | 5 | 1.34E-07 | 2.11E-07 | 7.45E-07 |
| VEEGAVKRINM | 431 | 441 | 10 | 4.97E-08 | 1.63E-08 | 5.18E-08 |
| VEEGAVKRINMAHL | 431 | 444 | 13 | 6.37E-09 | 2.24E-09 | 8.31E-09 |
| CIAGSHAVNG | 445 | 454 | 9 | 9.99E-10 | 7.77E-10 | 8.76E-10 |
| CIAGSHAVNGVA | 445 | 456 | 11 | 5.42E-10 | 7.31E-10 | 5.16E-10 |
| CIAGSHAVNGVARIHS | 445 | 460 | 15 | 8.90E-10 | 1.12E-09 | 1.09E-09 |
| CIAGSHAVNGVARIHSEIL | 445 | 463 | 18 | 1.02E-09 | 8.68E-10 | 9.94E-10 |
| AGSHAVNGVARIHSEIL | 447 | 463 | 16 | 7.06E-10 | 8.74E-10 | 1.07E-09 |
| VARIHSEILKKTIF | 455 | 468 | 13 | 3.69E-09 | 1.64E-09 | 2.63E-09 |
| KKTIFKDF | 464 | 471 | 7 | 2.36E-09 | 7.72E-09 | 1.32E-09 |
| YELEPHKFQNKTNGITPRRW | 472 | 491 | 17 | 1.18E-09 | 1.09E-09 | 1.13E-09 |
| CNPGLAEI | 495 | 502 | 6 | 5.21E-09 | 1.27E-08 | 4.73E-09 |
| AEIIAE | 500 | 505 | 5 | 4.84E-10 | 4.68E-10 | 7.84E-10 |
| EYISDL | 510 | 515 | 5 | 3.84E-08 | 1.06E-07 | 7.34E-08 |
| DQLRKLLS | 516 | 523 | 7 | 1.52E-09 | 3.75E-08 | 7.51E-09 |
| YVDDEA | 524 | 529 | 5 | 2.26E-07 | 5.36E-07 | 5.03E-07 |
| YVDDEAF | 524 | 530 | 6 | 1.13E-07 | 9.09E-08 | 4.92E-07 |
| FIRDVAKVKQENKLKF | 530 | 545 | 15 | 6.71E-10 | 2.09E-09 | 5.95E-10 |
| FIRDVAKVKQENKLKFAAY | 530 | 548 | 18 | 2.08E-09 | 5.91E-10 | 6.76E-10 |
| VAKVKQENKLKF | 534 | 545 | 11 | 6.97E-10 | 6.28E-10 | 3.21E-10 |
| VAKVKQENKLKFAAY | 534 | 548 | 14 | 1.86E-10 | 3.42E-09 | 4.98E-10 |
| LEREYKVHINPNSL | 549 | 562 | 12 | 1.94E-08 | 7.03E-08 | 1.07E-07 |
| EREYKVHINPNSL | 550 | 562 | 11 | 3.08E-08 | 1.27E-07 | 1.06E-07 |
| YKVHINPNSL | 553 | 562 | 8 | 1.24E-07 | 2.30E-07 | 2.07E-07 |
| FDVQVKRIHE | 563 | 572 | 9 | 3.47E-09 | 6.18E-10 | 8.67E-10 |
| FDVQVKRIHEYKRQLLNC | 563 | 580 | 17 | 2.03E-09 | 5.10E-10 | 6.12E-10 |
| HEYKRQLLNCLHVITLYN | 571 | 588 | 17 | 1.64E-08 | 9.24E-07 | 3.03E-08 |
| LHVITL | 581 | 586 | 5 | 3.42E-10 | 7.64E-10 | 1.09E-09 |
| YNRIKKEPNKFVVPRTVM | 587 | 604 | 15 | 4.14E-08 | 1.09E-08 | 1.30E-07 |
| IGGKAAPGYHMAKM | 605 | 618 | 12 | 3.36E-08 | 1.04E-09 | 1.57E-09 |
| IIKLIT | 619 | 624 | 5 | 5.42E-10 | 2.88E-10 | 2.54E-10 |
| IIKLITA | 619 | 625 | 6 | 9.92E-10 | 1.20E-09 | 1.29E-09 |
| ITAIGD | 623 | 628 | 5 | 1.11E-09 | 1.16E-09 | 1.27E-09 |
| ITAIGDVVNHDPVVGDRL | 623 | 640 | 16 | 6.49E-09 | 8.90E-09 | 1.44E-08 |
| IGDVVNHDPVVGDRL | 626 | 640 | 13 | 5.61E-09 | 1.10E-08 | 1.46E-08 |
| GDVVNHDPVVGDRL | 627 | 640 | 12 | 2.54E-09 | 2.48E-09 | 4.23E-08 |
| AEKVIPAAD | 653 | 661 | 7 | 1.02E-09 | 8.12E-10 | 8.17E-10 |
| AEKVIPAADL | 653 | 662 | 8 | 9.80E-10 | 9.23E-10 | 9.39E-10 |
| KVIPAADL | 655 | 662 | 6 | 8.95E-10 | 1.09E-09 | 3.83E-10 |
| QISTAGTEASGTGNMKF | 665 | 681 | 16 | 1.61E-09 | 2.92E-06 | 1.22E-09 |
| ISTAGTE | 666 | 672 | 6 | 6.81E-10 | 1.17E-09 | 3.16E-10 |
| ISTAGTEASGTGNMKF | 666 | 681 | 15 | 1.04E-09 | 5.45E-10 | 1.40E-09 |
| MLNGALTIGTMDGAN | 682 | 696 | 14 | 1.19E-09 | 1.00E-09 | 1.02E-09 |
| ALTIGTMDGAN | 686 | 696 | 10 | 9.95E-10 | 4.41E-10 | 8.50E-09 |
| TIGTMDGAN | 688 | 696 | 8 | 9.05E-10 | 4.03E-10 | 1.55E-09 |
| TIGTMDGANVE | 688 | 698 | 10 | 1.03E-09 | 1.38E-09 | 1.39E-09 |
| AEEAGEENF | 700 | 708 | 8 | 1.44E-09 | 1.68E-09 | 1.89E-08 |
| EAGEENF | 702 | 708 | 6 | 2.34E-08 | 4.27E-09 | 9.67E-08 |
| AGEENF | 703 | 708 | 5 | 2.05E-08 | 1.43E-07 | 1.51E-07 |
| MRVEDVDRL | 713 | 721 | 8 | 2.64E-08 | 1.33E-08 | 3.63E-08 |
| VDRLDQRGYNAQE | 718 | 730 | 12 | 2.37E-07 | 9.72E-07 | 1.64E-06 |
| DQRGYNAQE | 722 | 730 | 8 | 6.32E-07 | 2.60E-06 | 2.73E-06 |
| DQRGYNAQEY | 722 | 731 | 9 | 6.69E-07 | 3.04E-06 | 6.89E-06 |
| YYDRIPEL | 731 | 738 | 6 | 6.99E-10 | 6.37E-10 | 9.23E-10 |
| RQIIEQL | 739 | 745 | 6 | 8.27E-10 | 3.80E-10 | 2.22E-14 |
| SSGFFSPKQPDL | 746 | 757 | 9 | 6.20E-08 | 2.74E-08 | 5.88E-08 |
| FSPKQPDL | 750 | 757 | 5 | 1.53E-07 | 7.32E-07 | 4.21E-07 |
| PKQPDL | 752 | 757 | 3 | 2.84E-06 | 2.35E-06 | 5.88E-06 |

| | | | | | | |
|---|---|---|---|---|---|---|
| FKDIVN | 758 | 763 | 5 | 1.27E-08 | 2.87E-09 | 2.47E-09 |
| FKDIVNM | 758 | 764 | 6 | 9.48E-10 | 7.77E-10 | 8.89E-10 |
| LMHHDRF | 765 | 771 | 6 | 1.11E-09 | 1.05E-09 | 1.14E-09 |
| YVKCQE | 780 | 785 | 5 | 5.99E-10 | 8.52E-10 | 9.90E-10 |
| YKNPREWTRM | 791 | 800 | 8 | 1.20E-09 | 1.11E-09 | 1.37E-09 |
| VIRNIATSGKF | 801 | 811 | 10 | 8.93E-10 | 1.05E-09 | 9.28E-10 |
| VIRNIATSGKFSSD | 801 | 814 | 13 | 1.09E-09 | 9.93E-10 | 8.93E-10 |
| VIRNIATSGKFSSDRTIAQ | 801 | 819 | 18 | 9.23E-10 | 9.89E-10 | 5.91E-10 |
| IATSGKFSSDRTIAQ | 805 | 819 | 14 | 1.07E-09 | 5.76E-10 | 1.02E-09 |
| TSGKFSSD | 807 | 814 | 7 | 1.21E-09 | 6.58E-10 | 9.64E-10 |
| SSDRTIAQYAR | 812 | 822 | 10 | 1.33E-06 | 2.39E-06 | 1.75E-06 |
| YAREIW | 820 | 825 | 5 | 1.08E-09 | 1.08E-09 | 1.07E-09 |
| YAREIWGVE | 820 | 828 | 8 | 1.46E-09 | 2.42E-09 | 2.51E-09 |
| YAREIWGVEPSRQRLPAPDEK | 820 | 840 | 17 | 6.18E-07 | 7.45E-07 | 3.15E-06 |
| GVEPSRQRLPAPDEKIP | 826 | 842 | 12 | 3.76E-06 | 3.37E-06 | 1.89E-05 |
| PSRQRLPAPDEKIP | 829 | 842 | 9 | 4.95E-06 | 8.63E-06 | 1.18E-05 |
